# Annotation of ribosomal protein mass peaks in MALDI-TOF mass spectra of bacterial species and their phylogenetic significance

**DOI:** 10.1101/2020.07.15.203653

**Authors:** Wenfa Ng

## Abstract

Although MALDI-TOF mass spectrometry based microbial identification has achieved a level of accuracy that facilitate its use in classifying microbes to the species and strain level, questions remain on the identities of the mass peaks profiled from individual microbial species. Specifically, in the popular approach of comparing the mass spectrum of known and unknown microbes for identification purposes, the identities of the mass peaks were not taken into consideration. This study sought to determine if ribosomal proteins could account for some of the mass peaks profiled in MALDI-TOF mass spectra of different bacterial species. Using calculated molecular mass of ribosomal proteins for annotating mass peaks in bacterial species’ MALDI-TOF mass spectra downloaded from the SpectraBank database, this study revealed that ribosomal proteins could account for the low molecular weight mass peaks of <10000 Da. However, contrary to published reports, ribosomal proteins could not account for most of the mass peaks profiled. In particular, the data revealed that between 1 and 6 ribosomal protein mass peaks could be annotated in each mass spectrum. Annotated ribosomal proteins were S16, S17, S18, S20 and S21 from the small ribosome subunit, and L27, L28, L29, L30, L31, L31 Type B, L32, L33, L34, L35 and L36 from the large ribosome subunit. The ribosomal proteins with the most number of mass peak annotations were L36 and L29, with L34, L33, and L31 completing the list of ribosomal proteins with large number of annotations. Given the highly conserved nature of most ribosomal proteins, possible phylogenetic significance of the annotated ribosomal proteins were investigated through reconstruction of maximum likelihood phylogenetic trees. Results revealed that except for ribosomal protein L34, L31, L36 and S18, all annotated ribosomal proteins hold phylogenetic significance under the criteria of recapitulation of phylogenetic cluster groups present in the phylogeny of 16S rRNA. Phylogenetic significance of the annotated ribosomal proteins was further verified by the phylogenetic tree constructed based on the concatenated amino acid sequence of L29, S16, S20, S17, L27 and L35. Finally, analysis of the structure of the annotated ribosomal proteins did not reveal a high conservation of structure of the ribosomal proteins. Collectively, small low molecular weight (<10000 Da) ribosomal proteins could annotate some of the mass peaks in MALDI-TOF mass spectra of various bacterial species, and most of the ribosomal proteins hold phylogenetic significance. However, structural analysis did not identify a conserved structure for the annotated ribosomal proteins. Annotation of ribosomal protein mass peaks in MALDI-TOF mass spectra highlighted the deep biological basis inherent in the mass spectrometry-based microbial identification method. Subject areas biochemistry, biotechnology, microbiology, evolution, ecology

**Significance of the work:** While MALDI-TOF MS has been successfully used in identification of different microbes to the species and strain level through the comparison of mass spectra of known and unknown microbes, the approach (known as mass spectrum fingerprinting) remains lacking in the biological basis that underpins the technique. This study sought to uncover some of the biological basis that underpins MALDI-TOF MS microbial identification through the annotation of profiled mass peaks with ribosomal proteins. Previous studies have linked different ribosomal proteins to mass peaks in MALDI-TOF mass spectra of bacteria; however, broad spectrum verification of the finding across multiple species across different genera remain lacking. Using a collection of MALDI-TOF mass spectra of 110 bacterial species and strains catalogued in SpectraBank, this study sought to annotate ribosomal protein mass peaks in the mass spectra. Results revealed that small, low molecular weight ribosomal proteins of molecular mass < 10000 Da could annotate between 1 and 6 mass peaks in the catalogued mass spectra. This was smaller than the number of ribosomal proteins mass peaks postulated by previous studies. Overall, 16 ribosomal proteins (S16, S17, S18, S20, S21, L27, L28, L29, L30, L31, L31 Type B, L32, L33, L34, L35, and L36) were annotated with the most number of mass peaks annotations coming from L36 and L29. Reconstruction of phylogenetic trees of the annotated ribosomal proteins revealed that most of the ribosomal proteins hold phylogenetic significance with respect to the phylogeny of 16S rRNA. This provided further evidence that a deep biological basis is present in the approach of using mass spectrometry profiling of biomolecules for identifying bacterial species.

**Highlights:**

1. Ribosomal protein mass peaks were annotated in MALDI-TOF mass spectra of bacterial species across multiple genera.
2. Annotated ribosomal proteins were S16, S17, S18, S20, S21 for the small ribosome subunit, and L27, L28, L29, L30, L31, L31 Type B, L32, L33, L34, L35, L36 for the large ribosome subunit.
3. Between 1 and 6 ribosomal protein mass peaks were annotated per mass spectrum, a number significantly lower than that implied by other studies.
4. Annotated ribosomal proteins were small, low molecular weight ribosomal proteins of molecular mass < 10000 Da.
5. Phylogenetic tree reconstruction revealed the phylogenetic significance of most annotated ribosomal proteins except ribosomal protein L34, L31, L36 and S18.
6. Multi-locus sequence typing of L29, S16, S20, S17, L27 and L35 further showed the phylogenetic significance of ribosomal proteins in recapitulating the phylogeny of 16S rRNA.
7. Structural analysis of annotated ribosomal proteins did not find conserved structure. Thus, the reasons for the annotation of particular ribosomal proteins over others remain unknown.

## Introduction

Modern mass spectrometry tools have provided an unprecedented view of the cellular proteome such that the information obtained (i.e., mass spectra) could be used for various purposes such as the identification of proteins or tracing the phylogeny of a microbial species.^1,2^ Specifically, the soft ionization mass spectrometry technique of matrix-assisted laser desorption/ionization time-of-flight mass spectrometry (MALDI-TOF MS) has been utilized in the identification of various microbial species ranging from bacteria, fungi and archaea to the species and sometimes strain level with high accuracy.^3-7^

Briefly, whole cell samples of microorganisms are smeared onto a metal target plate, mixed with special MALDI matrixes that co-crystallized the cellular proteins, and a pulsed laser fired at the matrix-sample mixture to ionize the biomolecules for analysis by the mass spectrometer. The obtained mass spectrum provides a view of the cellular proteome with respect to the MALDI ionization technique and has been shown to be useful for discriminating between different species and strains of microbes. Specifically, a species of microbe would show a different set of mass peaks compared to another, and which thus allows the identification of different species and strains through the identification of species-specific mass peaks.

Given the myriad mass peaks profiled in a single mass spectrum and the large number of species in an identification exercise, computational algorithmic tools and software has been developed and deployed to aid the identification of microbial species.^8^ The most common approach used is known as mass spectrum fingerprinting.^8-10^ In this approach, the observation of the existence of unique mass spectrum for individual microbial species is utilized to build a reference database of mass spectra of known microorganisms. By comparing the mass spectrum of unknown microorganisms with those of known ones in a reference database, it is possible to identify the unknown microbe through the existence of species-specific mass peaks. Note that the mass spectrum fingerprinting approach does not require knowledge of the identities of the mass peaks profiled in the mass spectrum. What helps in identification is the existence of species-specific mass peaks in the profiled mass spectrum. Or from another perspective, a unique set of mass peaks exist for individual microbial species and helps in the building of unique mass spectrum fingerprints.

Besides mass spectrum fingerprinting, another approach exists for the identification of microorganisms through MALDI-TOF MS. Using a proteome database search approach, the method attempts to assign unique biomarkers for individual species and thus helps in identification.^11-13^ Typically, housekeeping proteins such as ribosomal proteins are profiled for their phylogenetic potential in identification of microbial species through the proteome database approach.^3,13,14^ However, compared to mass spectrum fingerprinting approach, the proteome database search approach suffers from a tedious workflow at first calculating the molecular masses of the candidate proteins, followed by a search for mass peaks in the mass spectrum that matched the calculated molecular masses of proteins. Thus, given the large size of the proteome of a microbial species, the proteome database search approach entails large investment of time and effort in calculating the molecular masses of proteins captured in the cellular proteome of the organism. With large number of proteomes available in proteomic database, the proteome database approach is simple in concept but difficult in implementation considering the time and effort needed to search for specific biomarker proteins and their assignment to mass peaks in a mass spectrum.

Thus, most vendors of MALDI-TOF MS microbial identification systems equipped their instruments with software that utilizes the mass spectrum fingerprinting approach for identifying microbial species. However, the instrument vendors typically require the users to purchase a costly reference database that is central to any identification effort based on the MALDI-TOF MS workflow. Currently, few academic studies have attempted to fully annotate the mass spectrum obtained from microbial species through the MALDI-TOF MS approach. Thus, the underlying biological basis of mass spectrum fingerprinting and, by extension, MALDI-TOF MS microbial identification remains nebulous.

Recently, efforts have been underway in understanding the biological basis of mass spectrometry-based microbial identification such as MALDI-TOF MS. Such efforts typically seek to understand the specific class of proteins that could help annotate a relatively large fraction of mass peaks in a mass spectrum. One example is ribosomal proteins. Many studies have attempted to annotate ribosomal proteins in the mass spectrum of microbial species and understand their phylogenetic significance.^3,13^ Knowledge gained in the process have helped anchor ribosomal proteins as important proteins able to confer phylogenetic significance to MALDI-TOF mass spectrum of microbial species.

Other efforts aimed at providing a more detailed look into the biological basis of MALDI-TOF MS microbial identification surveyed mass spectra of microorganisms curated in publicly accessible mass spectrum database. Specifically, conserved mass peaks of microbial species have been found at the species and genus levels that confirmed the biological basis of the MALDI-TOF MS microbial identification approach.^15^ Future annotation of the conserved mass peaks would provide additional layers of information for distilling the biological mysteries of the set of biomolecules ionized and profiled by the MALDI-TOF mass spectrometer.

Using mass spectra curated by the SpectraBank database (https://www.usc.es/gl/investigacion/grupos/lhica/spectrabank/Database.html), this study attempted to annotate potential ribosomal protein mass peaks in the mass spectra of 110 species and strains profiled in the database. Beyond annotation of ribosomal proteins, the study also sought to understand the phylogenetic significance of the annotated ribosomal proteins from both a sequence and structure perspective. Thus, the study would start off with the calculation of the molecular masses of the set of ribosomal proteins in the proteome of microbial species profiled in the SpectraBank database. Proteomic information used would be downloaded from the UniProt database. This would be followed by manual annotation of the ribosomal protein mass peaks of the peak list of the various microbial species catalogued in the SpectraBank database. Following this, phylogenetic analysis would be conducted to determine the level of concordance between the 16S rRNA phylogenetic tree and that of the annotated ribosomal proteins of microbes with a particular ribosomal protein mass peak. Structural analysis of the annotated ribosomal proteins would also be conducted to determine if structural homology exist between different annotated ribosomal protein. Finally, a multi-locus sequence typing approach would be used to determine if analysis of concatenated ribosomal protein amino acid sequence could explain the phylogeny of microbial species with annotated ribosomal protein mass peaks.

## Materials and Methods

### Mass spectra, proteome and 16S rRNA information

Mass spectra and peak lists of microbial species were downloaded from SpectraBank (https://www.usc.es/gl/investigacion/grupos/lhica/spectrabank/Database.html) and constitute the basis of this work. Proteomes of the microbes catalogued in the SpectraBank database were downloaded from UniProt and used in the calculation of molecular mass of the ribosomal proteins profiled from the respective proteome. Molecular weight calculations were performed with the online calculator: “Compute pI/Mw Tool” (https://web.expasy.org/compute_pi/). 16S rRNA gene sequences of the microbial species catalogued in the SpectraBank database were downloaded from the SILVA database (https://www.arb-silva.de/).

### Annotation of ribosomal protein mass peaks

Molecular masses of the ribosomal proteins of the large and small ribosome subunits were used in the annotation of ribosomal protein mass peaks in the peak list of the respective microbial species and strains. Given that differences and deviation in calibration of the mass spectrometer could result in slight differences to the detected molecular weight of the respective proteins by matrix-assisted laser desorption/ionization time-of-flight mass spectrometer (MALDI-TOF MS), a tolerance of 10 Da was applied in annotating mass peaks that could be accounted for by ribosomal proteins. Briefly, if the molecular weight of the mass peak did not differ by more than 10 Da from the calculated molecular mass of the ribosomal protein, the mass peak could be annotated as a ribosomal protein mass peak.

### Reconstruction of phylogenetic tree based on ribosomal proteins and 16S rRNA sequence

Protein sequence of ribosomal proteins of bacterial species with annotated ribosomal protein mass peaks were used in reconstruction of maximum likelihood phylogenetic tree (based on default parameters) after the amino acid sequences were aligned with ClustalW algorithm in MEGA X software^16^ (Molecular Evolutionary Genetic Analysis software, https://www.megasoftware.net/) using default parameters. The phylogenetic tree based on ribosomal proteins were compared with the maximum likelihood phylogenetic tree of 16S rRNA of the same species. Given the computational demands of aligning long sequences of 16S rRNA of large number of bacterial species (n > 22) on a budget laptop, the phylogenetic trees of 16S rRNA of bacterial species with an annotated ribosomal protein of L36, L29 and L34 were not able to be reconstructed. Analysis of the concordance between 16S rRNA and ribosomal protein phylogenetic trees was based on the presence/absence of specific phylogenetic cluster.

Besides the reconstruction of ribosomal protein phylogenetic tree for bacterial species with a corresponding annotated ribosomal protein, phylogenetic tree for the same ribosomal protein was also reconstructed for a common set of bacterial species belonging to different genera represented in the SpectraBank database. Doing so helped identify whether ribosomal protein could confer phylogenetic significance to bacterial species with and without a corresponding annotated ribosomal protein. Specifically, it tested the hypothesis concerning the biological meaning of an annotated ribosomal protein mass peak in the MALDI-TOF mass spectrum of a bacterial species. The set of bacterial species utilized for this analysis is available in Table S112 of Supplementary materials.

### Multi-locus sequence typing analysis

Given that multiple ribosomal protein could better describe the phylogeny of bacterial species, a multi-locus sequence typing approach was used in concatenating the amino acid sequence of multiple ribosomal proteins in the full set of bacterial species in Table S112 for phylogenetic tree reconstruction analysis. The type and sequence for the concatenation was N-terminus to C-terminus, L29-S16-S20-S17-L27-L35. Phylogenetic tree reconstruction was carried out after ClustalW alignment of the amino acid sequence of different bacterial species and maximum likelihood phylogenetic tree was selected.

### Structural analysis of annotated ribosomal proteins

Given the likely co-evolution between ribosomal proteins and between ribosomal proteins and ribonucleic acids in both the large and small ribosome subunits, structural analysis is necessary to detect possible conservation of structure between different ribosomal proteins. To this end, the structure of all annotated ribosomal proteins (L27, L28, L29, L30, L31, L31 Type B, L32, L33, L34, L35, L36, S16, S17, S18, S20, S21) were modelled with the Phyre^2^ server (http://www.sbg.bio.ic.ac.uk/phyre2/html/page.cgi?id=index) and visualized with Chimera 1.13 software (https://www.rbvi.ucsf.edu/chimera/). Amino acid sequence of the respective ribosomal proteins was obtained from the proteome of *Bacillus subtilis* strain 168 as model organism.

## Results

### Annotation of ribosomal protein mass peak

Annotation of ribosomal protein mass peaks was carried out by comparing the calculated molecular masses of ribosomal proteins with molecular weight less than 10000 Da with the mass/charge (*m/z*) ratio of mass peaks in the MALDI-TOF mass spectrum of bacterial species catalogued in the SpectraBank database. The charge of the mass peaks was assumed to be +1 given that most MALDI-TOF mass spectrometers were operated in the positive linear ion mode that conferred a +1 electrical charge to the ionized biomolecules. A ribosomal protein mass peak was annotated when there was less than a 10 Da mass difference between the calculated molecular weight of the ribosomal protein and the mass/charge ratio of the pertinent mass peak. A total of 110 microbial species and strains were catalogued in the database and the results of the analysis were presented as Table S1 to S110 in the Supplementary material of this manuscript. The list of bacterial species represented in the SpectraBank database is available as Table S111 of the Supplementary materials.

Overall, between 1 and 6 ribosomal protein mass peaks could be annotated in the mass spectrum of different bacterial species and strains, with a few species and strains with no annotated ribosomal protein mass peaks. This result differed from that of the literature where it was reported that ribosomal proteins account for a substantial proportion of the total mass peaks profiled in a MALDI-TOF mass spectrum.^3,13^ Table 1 shows the ribosomal proteins with annotated mass peaks as well as the total number of annotations achieved for each annotated ribosomal protein.

**Table 1:** Ribosomal proteins with annotated mass peaks.

| **Ribosomal protein** | **No. of annotations** |
| --- | --- |
| L36 | 82 |
| L29 | 78 |
| L34 | 53 |
| L33 | 34 |
| L31 | 16 |
| L30 | 9 |
| L32 | 9 |
| S18 | 5 |
| S16 | 4 |
| S20 | 2 |
| S21 | 2 |
| L28 | 2 |
| S17 | 2 |
| L31 Type B | 2 |
| L27 | 2 |
| L35 | 2 |

Specifically, the data revealed that only 16 ribosomal proteins from the large and small ribosome subunit could find corresponding mass peaks in the MALDI-TOF mass spectrum of the profiled microbial species and strains. Annotated ribosomal proteins from the large ribosome subunit were L36, L29, L34, L33, L31, L30, L32, L28, L31 Type B, L27 and L35, while annotated ribosomal proteins from the small ribosome subunit were S18, S16, S20, S21 and S17. From the molecular weight perspective, all annotated ribosomal proteins have molecular mass less than 10000 Da, which was lower than the typical mass range of between 10000 Da to 20000 Da for ribosomal proteins. More importantly, annotated ribosomal proteins came from the cluster from L27 to L36 for the large ribosome subunit and from S17 to S21 for the small ribosome subunit. In addition, the data revealed that none of the annotated ribosomal protein from the small ribosome subunit had more than 5 annotations in mass spectra of the microbial species and strains, which suggested that ribosomal proteins from the small subunit might be less important to cellular physiology and thus had a lower relative abundance beyond that required to constitute the ribosome. On the other hand, ribosomal proteins from the large ribosome subunit accounted for a larger fraction of annotations, which suggested that they might be more important physiologically and thus were expressed at higher relative abundance beyond that required to constitute the ribosome.

Overall, 7 large ribosome subunit ribosomal proteins had the most number of mass peak annotations. They are ribosomal protein L36, L29, L34, L33, L31, L30 and L32. In particular, ribosomal protein L36 and L29 had the most number of mass peak annotations. Ribosomal protein L36 is known to be important for the structural stability of the large ribosome subunit, but questions remain on why particular sets of ribosomal protein such as those described above dominate the annotation of ribosomal protein mass peaks. It must be noted that for the MALDI-TOF mass spectra catalogued in the SpectraBank database, the molecular weight cut-off for the mass spectrometry analysis of bacteria appeared to be 10000 *m/z*, which corresponded to protein of molecular weight 10000 Da assuming a charge of +1. Thus, only relatively low molecular weight ribosomal proteins were annotated, and this may constitute a bias in the analysis.

Given that ribosomal proteins generated during expression of the corresponding ribosomal protein genes should be roughly balanced between the different proteins that constituted the ribosome, the preferential detection of particular ribosomal proteins by MALDI-TOF MS during mass spectrum analysis of whole cell bacteria opens up several questions concerning the reasons underlying the selective overabundance of specific ribosomal proteins such as L36 and L29 as well as possible bias in the profiling of the cellular proteome by MALDI-TOF MS.

One reason that could account for the selective overabundance of ribosomal protein L36 and L29 that led to their repeated annotation in the mass spectrum of bacterial cells could be the presence of as-yet unknown physiological function of the ribosomal protein that led to their preferential over-expression. Another factor could be the preferential ionization of particular ribosomal protein during the MALDI-TOF MS ionization process.

### Phylogenetic analysis of annotated ribosomal proteins

The study next turned to the analysis of the phylogenetic significance of the annotated ribosomal proteins. In particular, the key question of interest is whether the phylogenetic tree based on the annotated ribosomal protein share similarity with that based on 16S rRNA of the same species. To this end, amino acid sequences of the respective annotated ribosomal protein were collected from the corresponding proteome of bacterial species with the annotated ribosomal protein, aligned with ClustalW algorithm, and maximum likelihood phylogenetic tree generated with MEGA X software. Proteomes of the bacterial species were downloaded from UniProt.

Given that phylogenetic tree could be generated for any gene or protein, and are not 100% similar to each other, high level of concordance between phylogenetic tree was defined as the presence/absence of specific phylogenetic cluster group of bacterial species. From another perspective, the phylogenetic analysis sought to uncover if there are similar branches of phylogeny (or phylogenetic cluster group) present in the phylogenetic tree of 16S rRNA and the candidate ribosomal protein. Thus, differences are likely to exist between phylogenetic trees with high level of concordance in phylogenetic branches or phylogenetic cluster group.

Due to the computational demand from aligning long nucleotide sequences of 16S rRNA of many bacterial species with an annotated ribosomal protein L36, the phylogenetic trees of 16S rRNA for checking the phylogeny of ribosomal protein L36, L29 and L34 could not be computed. Thus, the analysis presented below will compare the phylogenetic trees of ribosomal proteins L30, L31, L32 and L33 with the corresponding 16S rRNA phylogenetic tree of the same set of bacterial species. The phylogenetic trees of ribosomal protein L36, L29 and L34 would be presented independently.

The phylogenetic tree based on ribosomal protein L36 is as shown in Figure 1. Species of the same genus such as *Bacillus, Staphylococcus, Listeria, Vibrio, Shewanella, Proteus* and *Pseudomonas* tend to cluster close to each other, which revealed that the maximum likelihood method was capable of detecting close phylogeny between species of the same genus. The analysis results also revealed that there was high level of sequence conservation of amino acid sequences of ribosomal proteins from bacterial species belonging to the same genus. Thus, ribosomal protein L36 does endow phylogenetic significance which could be used to inform species provenance and relatedness with other microbial species.

**Figure 1:**
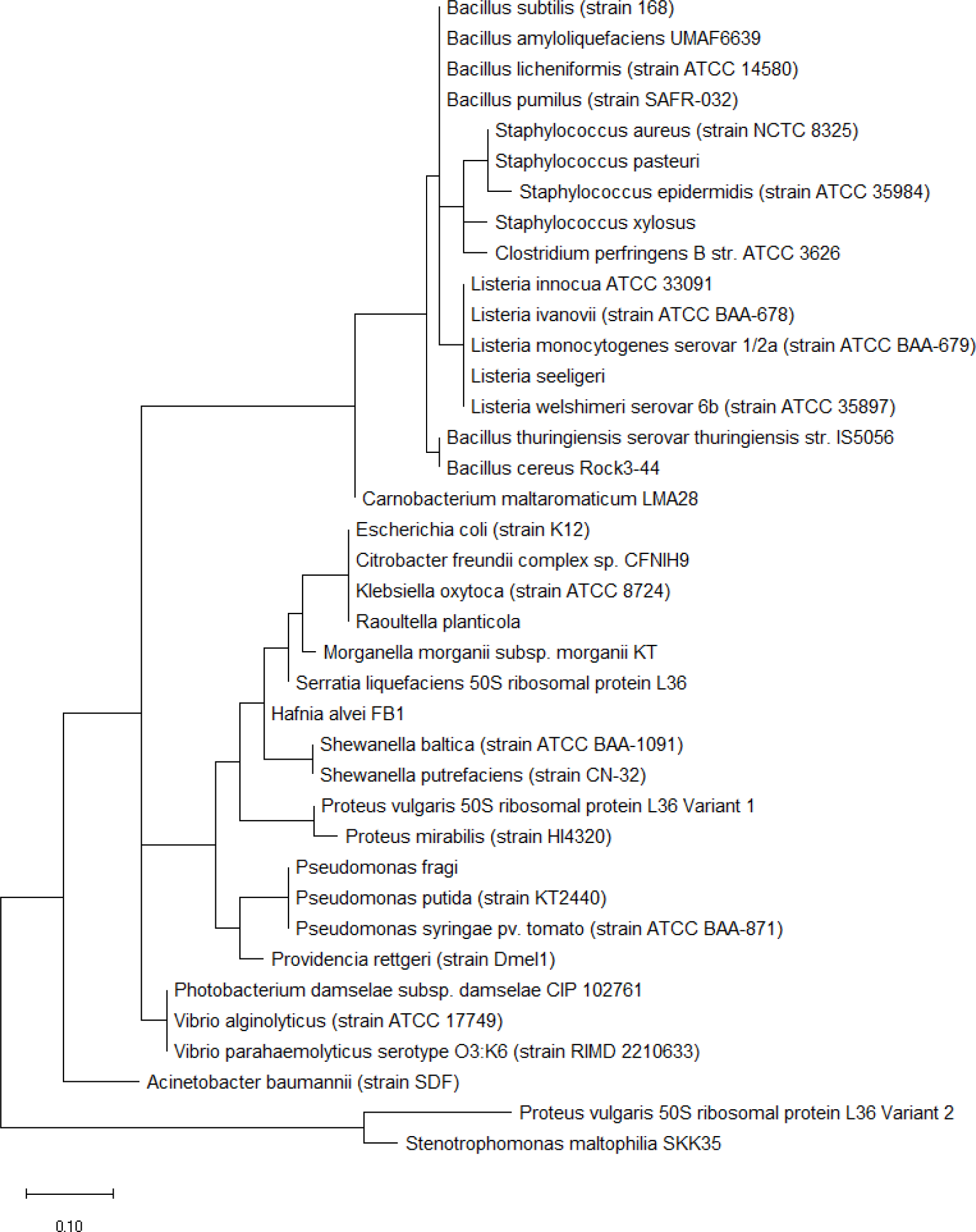
Maximum likelihood phylogenetic tree of ribosomal protein L36 in bacterial species whose MALDI-TOF mass spectrum contains a corresponding ribosomal protein L36 mass peak. Note that variant 1 and 2 refers to different variants of the same protein.

Maximum likelihood phylogenetic tree based on ribosomal protein L29 is shown in Figure 2 and revealed that bacterial species of the same genus such as *Bacillus, Listeria, Clostridium, Shewanella, Vibrio, Photobacterium, Pseudomonas, Serratia, Enterobacter, Providencia, Proteus*, and *Klebsiella* tend to cluster together. This indicated that ribosomal protein L29 could explain the provenance of the different species and holds phylogenetic significance in informing the phylogenetic relatedness between different bacterial species. High level of conservation in amino acid sequence of ribosomal protein L29 between different species of the same genus likely explains the clustering effect observed.

**Figure 2:**
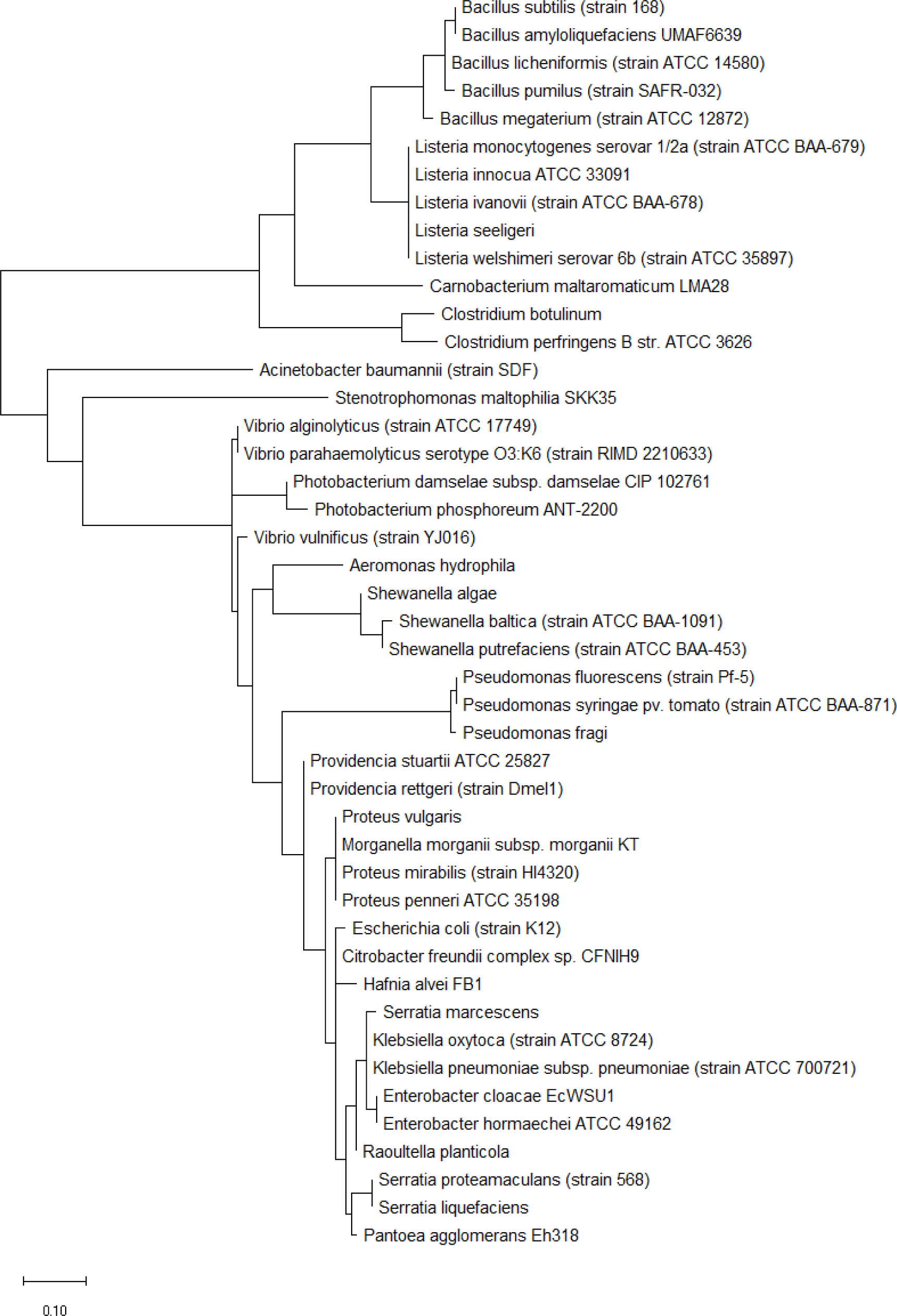
Maximum likelihood phylogenetic tree based on ribosomal protein L29 of bacterial species whose MALDI-TOF mass spectrum contain a ribosomal protein L29 mass peak.

Figure 3 shows the maximum likelihood phylogenetic tree based on ribosomal protein L34 of different bacterial species. While the phylogenetic tree could explain the close evolutionary relationship between different species of the same genus of genera such as *Vibrio, Clostridium, Pseudomonas*, and *Proteus*, it nevertheless cluster together species of different genera such as *Escherichia, Enterobacter, Morganella, Providencia*, and *Klebsiella*. This revealed that ribosomal protein L34 might not be suitable for informing the phylogenetic relationship between different species on the basis of 16S rRNA classification which did not place species of genera such as *Escherichia, Enterobacter, Morganella, Providencia*, and *Klebsiella* in the same cluster group. However, from another perspective, the data highlighted that the ribosomal protein L34 of species in the genera of *Escherichia, Enterobacter, Morganella, Providencia*, and *Klebsiella* are closely related, which pointed to possible horizontal exchange of the ribosomal protein in the distant past.

**Figure 3:**
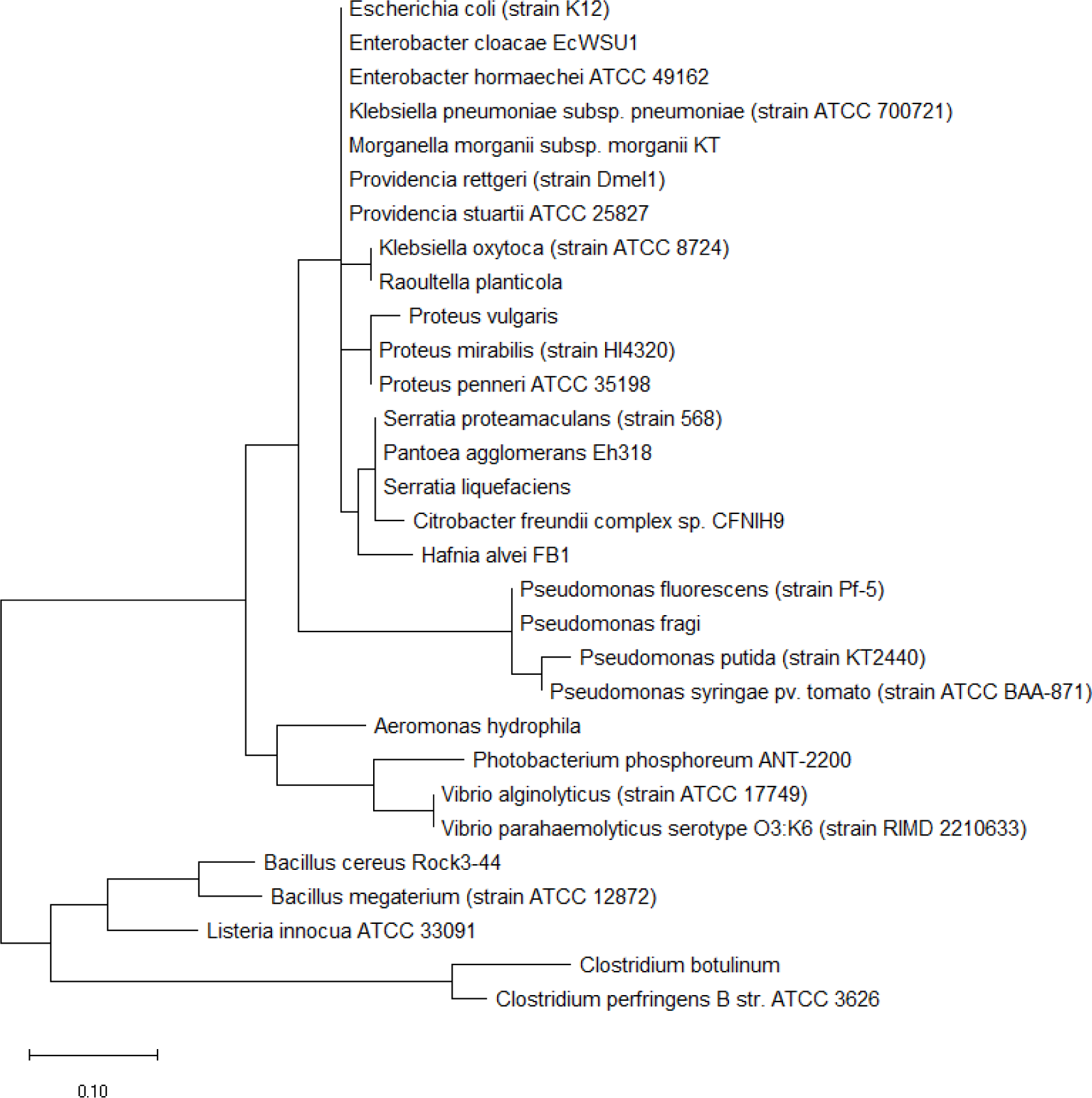
Maximum likelihood phylogenetic tree based on ribosomal protein L34 of bacterial species whose MALDI-TOF mass spectrum contains a ribosomal protein L34 mass peak.

Similar to the case for ribosomal protein L36, L29 and L34, phylogenetic tree based on the respective ribosomal protein with high number of mass peak annotations (i.e., at least 9 annotations) were reconstructed and compared with the corresponding 16S rRNA maximum likelihood phylogenetic tree for the same set of bacterial species. Comparing the maximum likelihood phylogenetic tree of ribosomal protein L30 with that of 16S rRNA of the same species, Figure 4a and 4b revealed that ribosomal protein L30 could explain the phylogeny between the species to a large extent and thus hold phylogenetic significance. Small difference in the placement of *Morganella morganii* could be due to microevolution at the amino acid level.

**Figure 4a:**
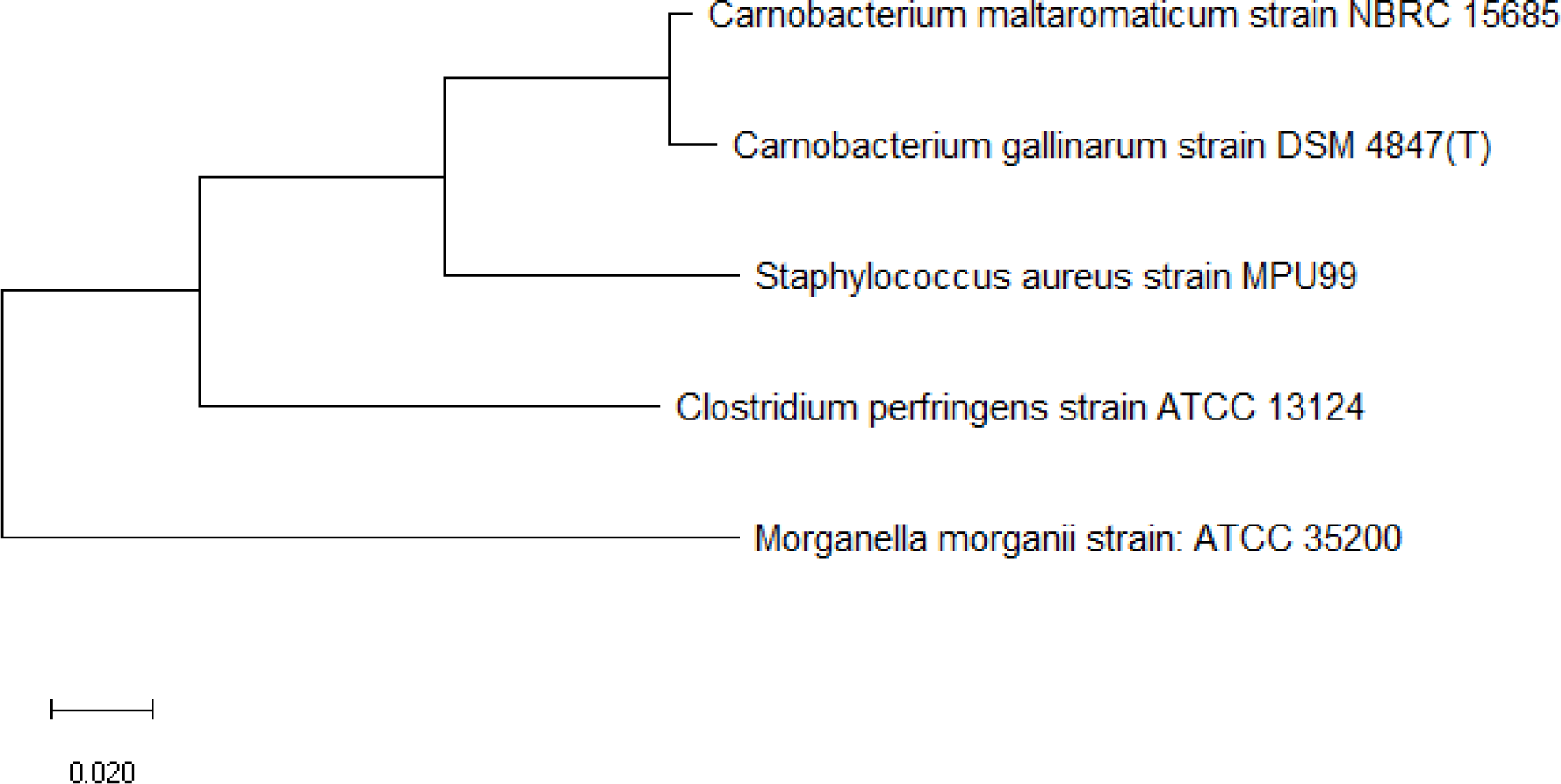
Maximum likelihood phylogenetic tree based on 16S rRNA gene sequences of bacterial species with a ribosomal protein L30 mass peak in their MALDI-TOF mass spectrum.

**Figure 4b:**
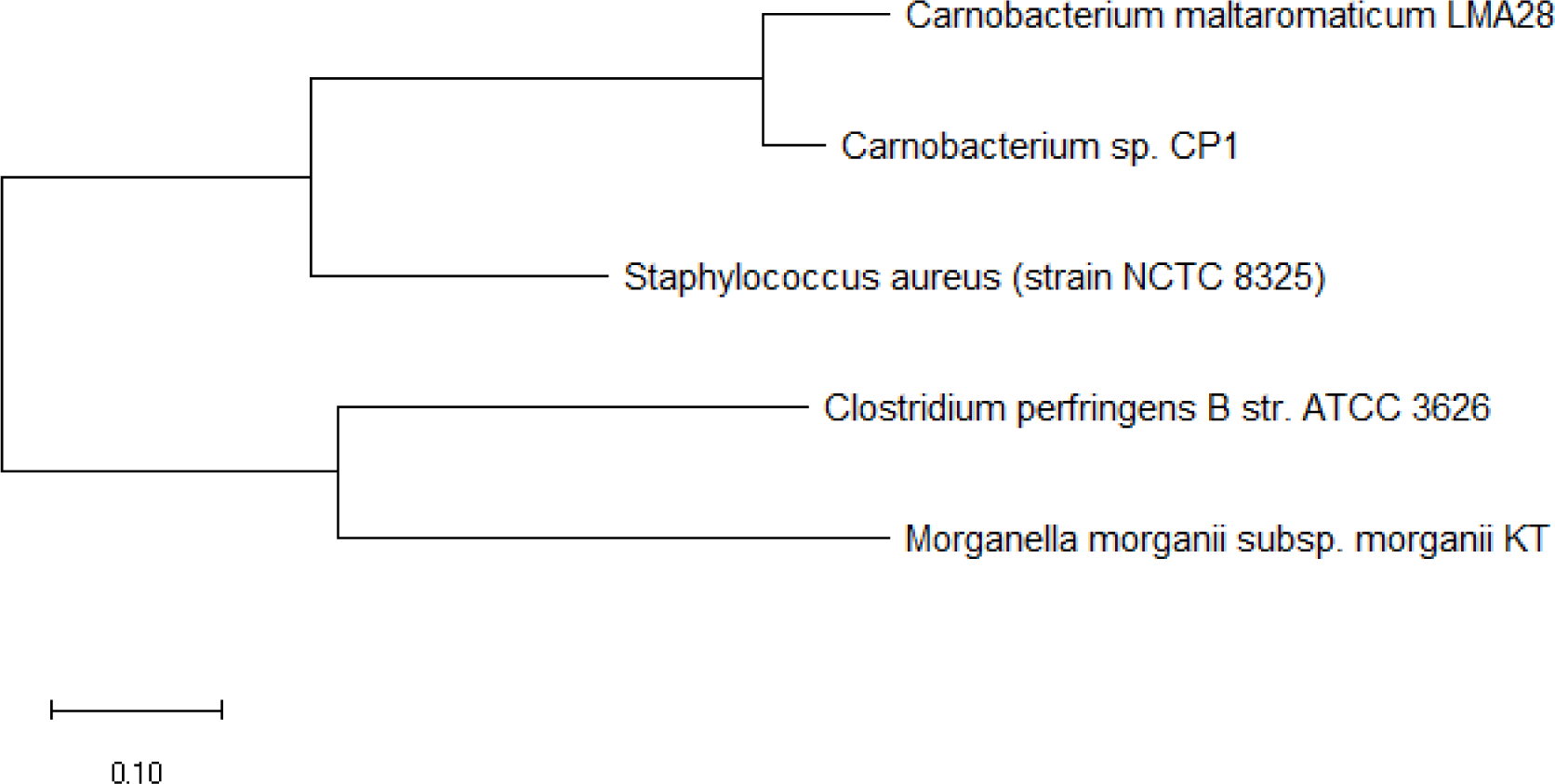
Maximum likelihood phylogenetic tree based on ribosomal protein L30 of bacterial species whose MALDI-TOF mass spectrum contains a ribosomal protein L30 mass peak.

Figure 5a and 5b shows the phylogenetic tree based on 16S rRNA and ribosomal protein L31 of a set of bacterial species with an annotated ribosomal protein L31 mass peak in their MALDI-TOF mass spectrum. The data revealed that ribosomal protein L31 could replicate phylogenetic cluster 1 and 2 in its maximum likelihood phylogenetic tree, which indicated that the ribosomal protein could explain the evolutionary relationship between bacterial species in the phylogenetic cluster groups and their relative arrangement in the phylogenetic tree.

**Figure 5a:**
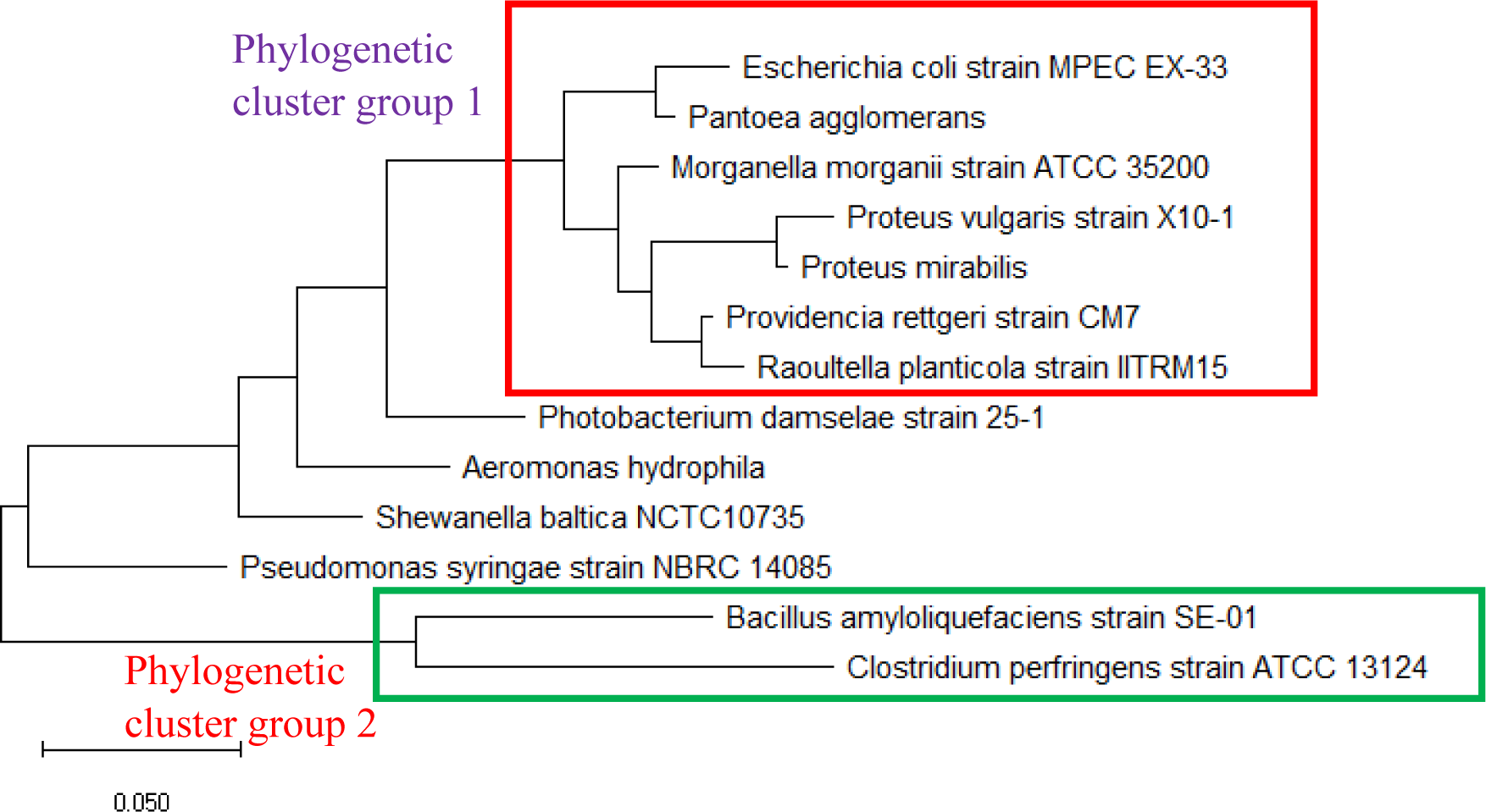
Maximum likelihood phylogenetic tree based on 16S rRNA gene sequence of bacterial species with a ribosomal protein L31 mass peak in their MALDI-TOF mass spectrum.

**Figure 5b:**
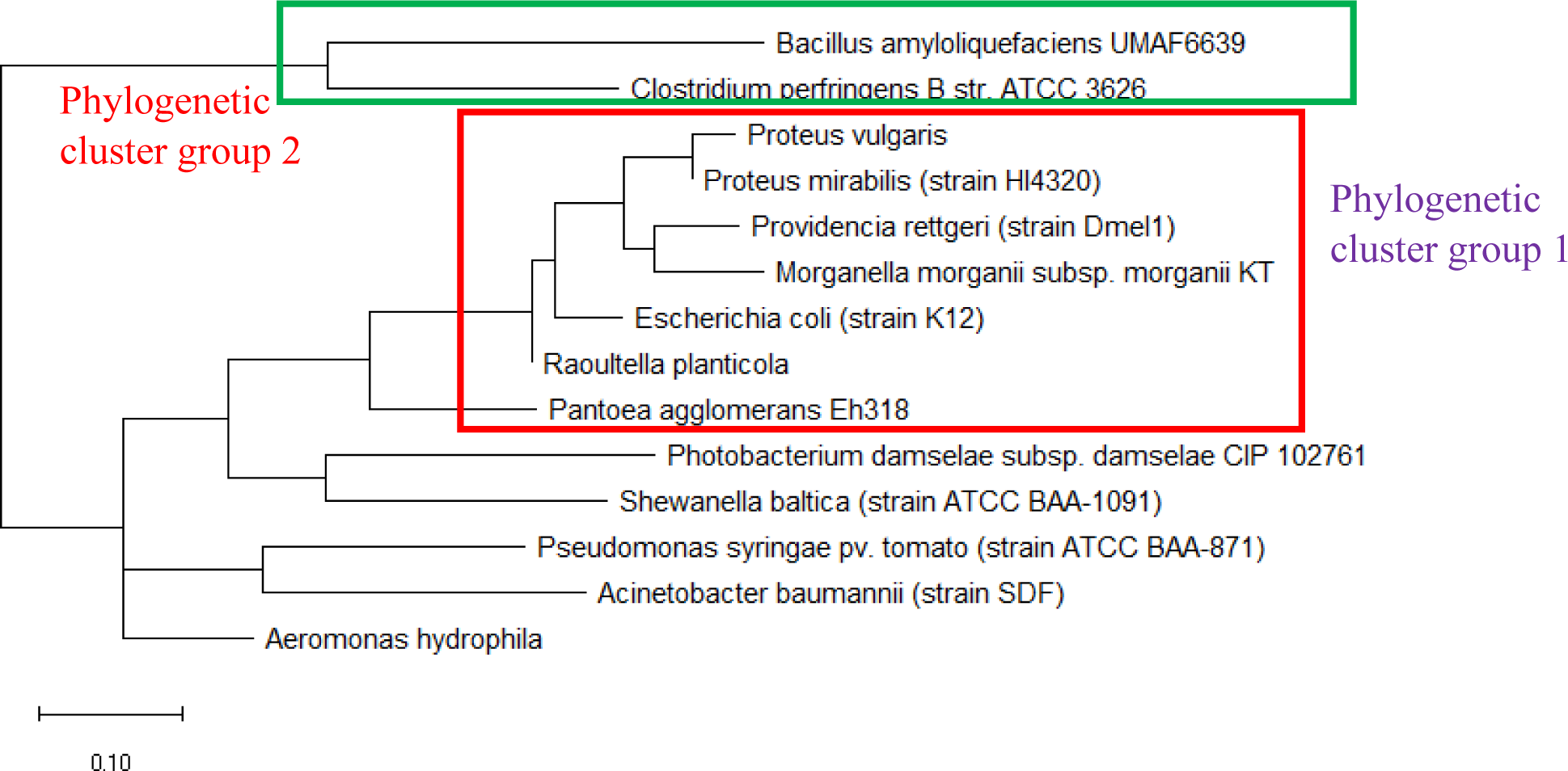
Maximum likelihood phylogenetic tree based on ribosomal protein L31 of bacterial species whose MALDI-TOF mass spectrum contains a ribosomal protein L31 mass peak.

Maximum likelihood phylogenetic tree based on 16S rRNA and ribosomal protein L32 of the same set of bacterial species with an annotated ribosomal protein L32 mass peak in their MALDI-TOF mass spectrum were shown in Figure 6a and 6b. Overall, ribosomal protein L32 could explain the phylogenetic relationships between the species as depicted in the phylogenetic tree based on 16S rRNA of the same species. For example, ribosomal protein L32 placed the three *Listeria* species in the same phylogenetic cluster similar to that endowed by the 16S rRNA phylogenetic tree probably due to the high level of sequence conservation between ribosomal protein L32 of the three *Listeria* species. This revealed that ribosomal protein L32 holds phylogenetic significance in explaining the evolutionary relationships between species.

**Figure 6a:**
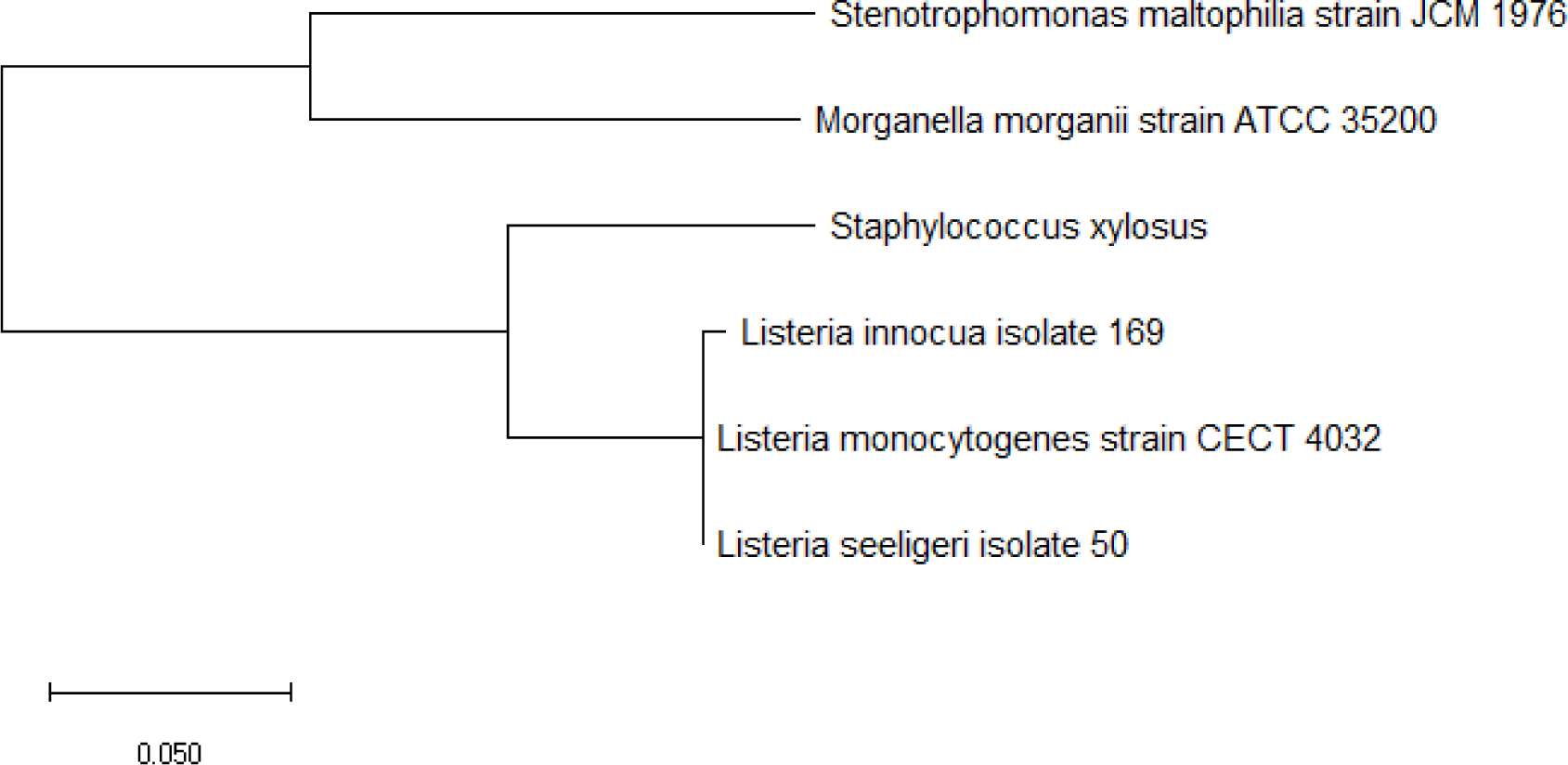
Maximum likelihood phylogenetic tree based on 16S rRNA gene sequence of bacterial species with a ribosomal protein L32 mass peak in their MALDI-TOF mass spectrum.

**Figure 6b:**
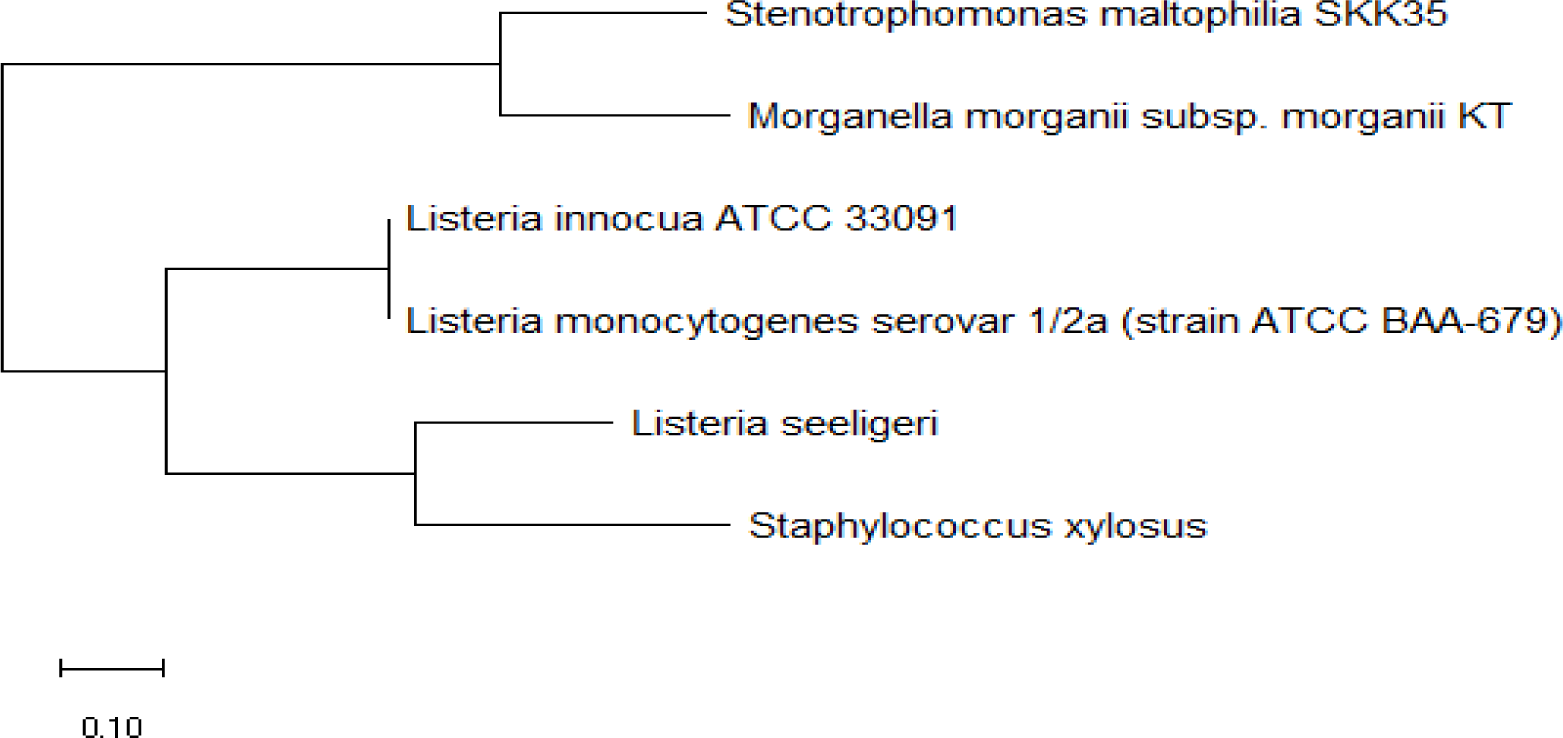
Maximum likelihood phylogenetic tree based on ribosomal protein L32 of bacterial species whose MALDI-TOF mass spectrum contains a ribosomal protein L32 mass peak.

Figure 7 shows the phylogenetic tree based on ribosomal protein L33 and 16S rRNA of the same bacterial species. The data revealed that ribosomal protein L33 holds phylogenetic significance given that phylogenetic tree based on the protein could correctly cluster bacterial species from the same genus such as *Bacillus, Pseudomonas, Providencia*, and *Listeria*. More importantly, the phylogenetic tree based on ribosomal protein L33 also correctly placed species into respective phylogenetic cluster group 1 and 2 in comparison with that based on the 16S rRNA gene of the same species.

**Figure 7a:**
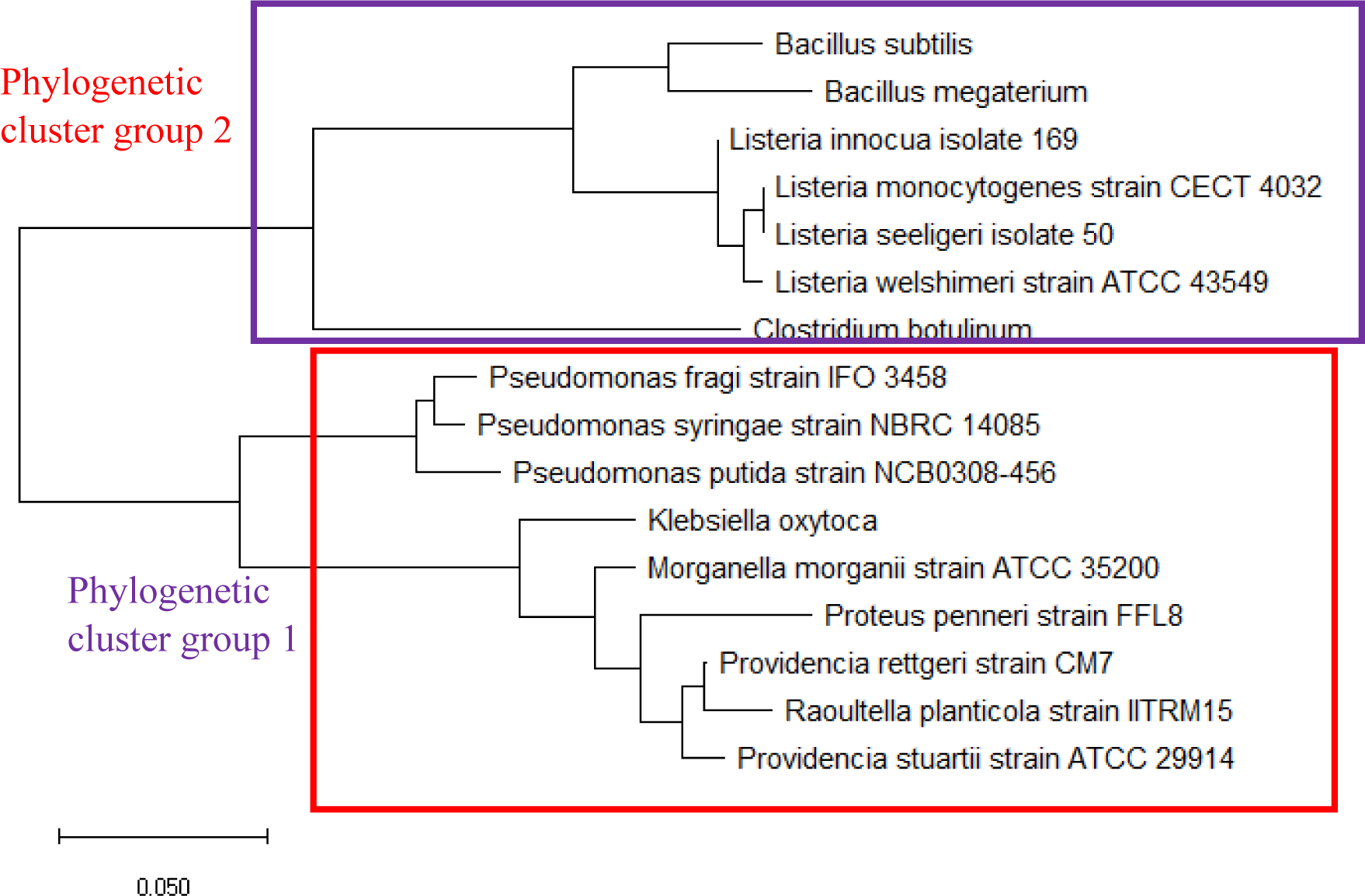
Maximum likelihood phylogenetic tree based on 16S rRNA gene sequences of bacterial species with a ribosomal protein L33 mass peak in their MALDI-TOF mass spectrum.

**Figure 7b:**
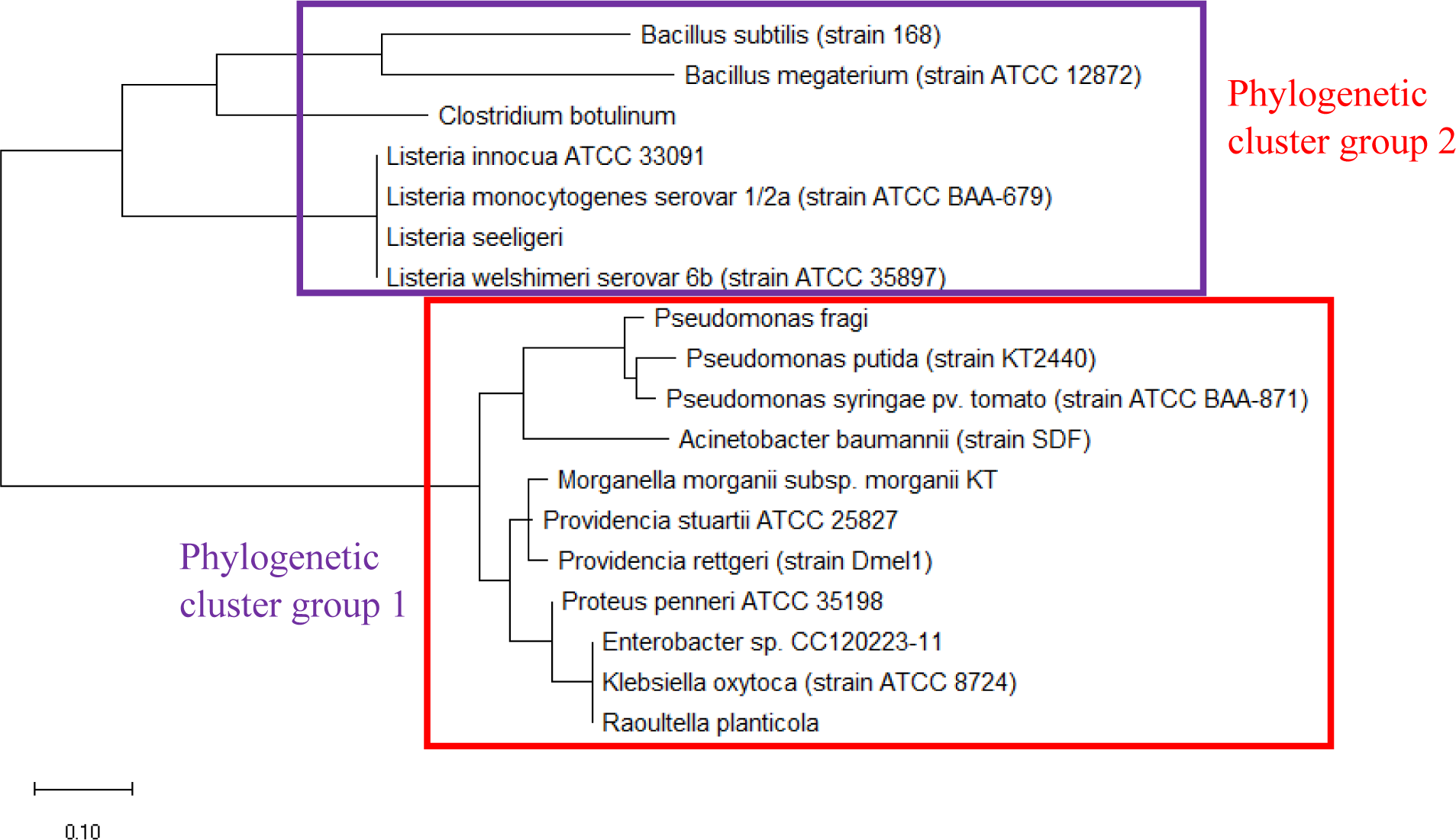
Maximum likelihood phylogenetic tree based on ribosomal protein L33 of bacterial species whose MALDI-TOF mass spectrum contains a ribosomal protein L33 mass peak.

Overall, the above phylogenetic analysis revealed that except for ribosomal protein L34, all the other ribosomal proteins with large number of mass peak annotations hold phylogenetic significance. However, the phylogenetic analysis of the respective ribosomal protein was conducted for bacterial species with an annotated mass peak of the same ribosomal protein. This naturally leads to the question of whether the phylogenetic significance exhibited by the respective ribosomal protein could be replicated for bacterial species without an annotated mass peak of the same ribosomal protein. Thus, phylogenetic analysis was conducted for all annotated ribosomal proteins using a set of bacterial species where a single representative was chosen from each genus. The list of 22 bacterial species chosen for the analysis is as described in Table S112.

As a reference for comparing the phylogenetic tree based on different ribosomal protein, the maximum likelihood phylogenetic tree based on 16S rRNA gene of the set of 22 bacterial species was reconstructed after ClustalW alignment of the 16S rRNA nucleotide sequence. Figure 8 revealed that *Carnobacterium maltaromaticum, Bacillus subtilis, Listeria monocytogenes, Staphylococcus aureus*, and *Clostridium botulinum* all fall into phylogenetic cluster group 1. On the other hand, *Hafnia alvei, Morganella morganii, Proteus vulgaris, Providencia rettgeri*, and *Raoultella planticola* are grouped into phylogenetic cluster group 2.

**Figure 8:**
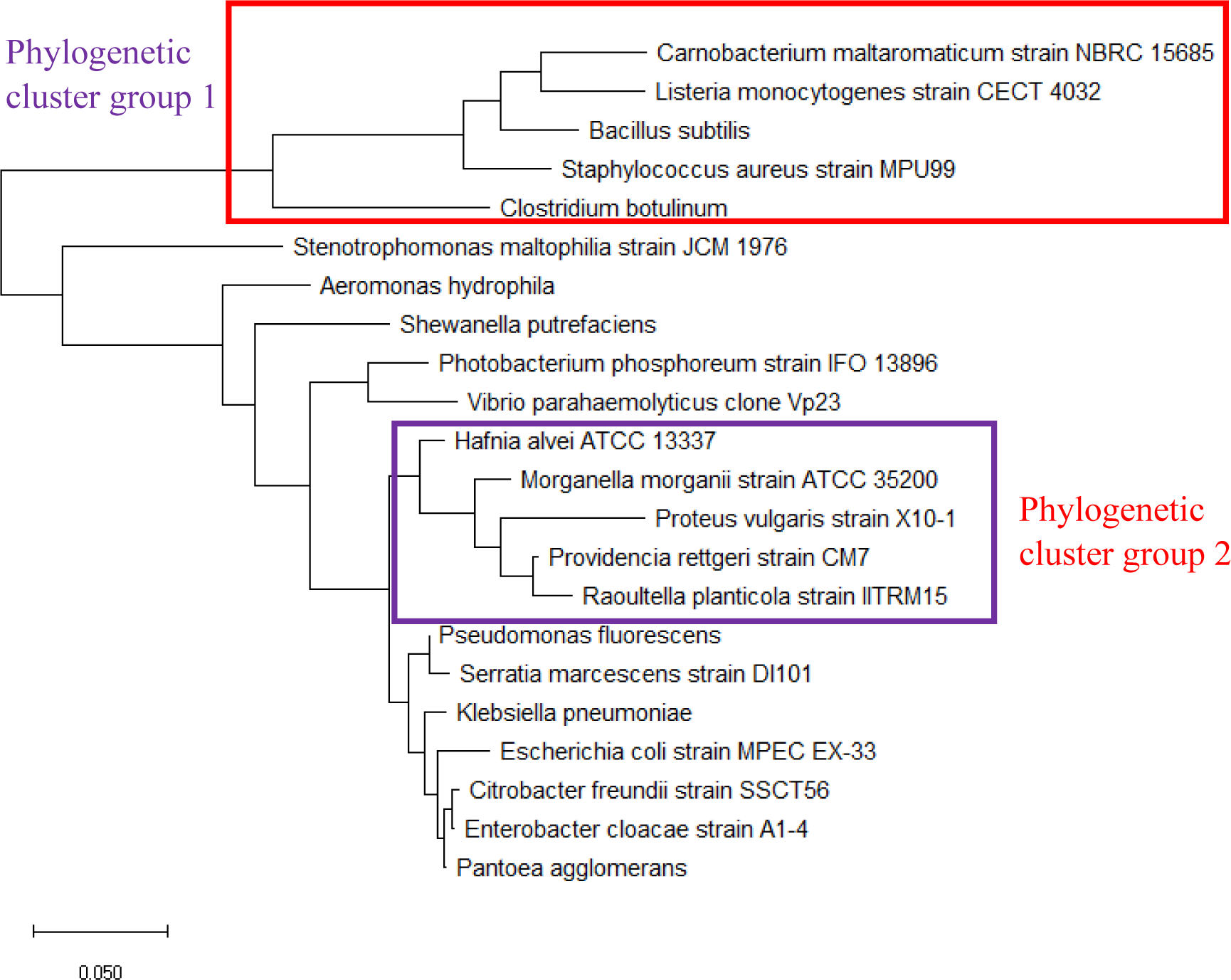
Maximum likelihood phylogenetic tree based on 16S rRNA gene sequence of selected set of bacterial species.

Figure 9 shows the maximum likelihood phylogenetic tree based on ribosomal protein L27 for the selected group of bacteria listed in Table S112. Except for phylogenetic cluster group 1, which was in common with that depicted in the phylogenetic tree based on 16S rRNA of the same group of bacterial species, there were substantial differences to the phylogenetic tree based on ribosomal protein L27 and 16S rRNA gene. This indicated that the evolutionary trajectories of ribosomal protein L27 and 16S rRNA was dissimilar and that they did not have a strong co-evolutionary relationship. However, ribosomal protein L27 still holds phylogenetic significance as it could replicate the correct placing of phylogenetic cluster group 1.

**Figure 9:**
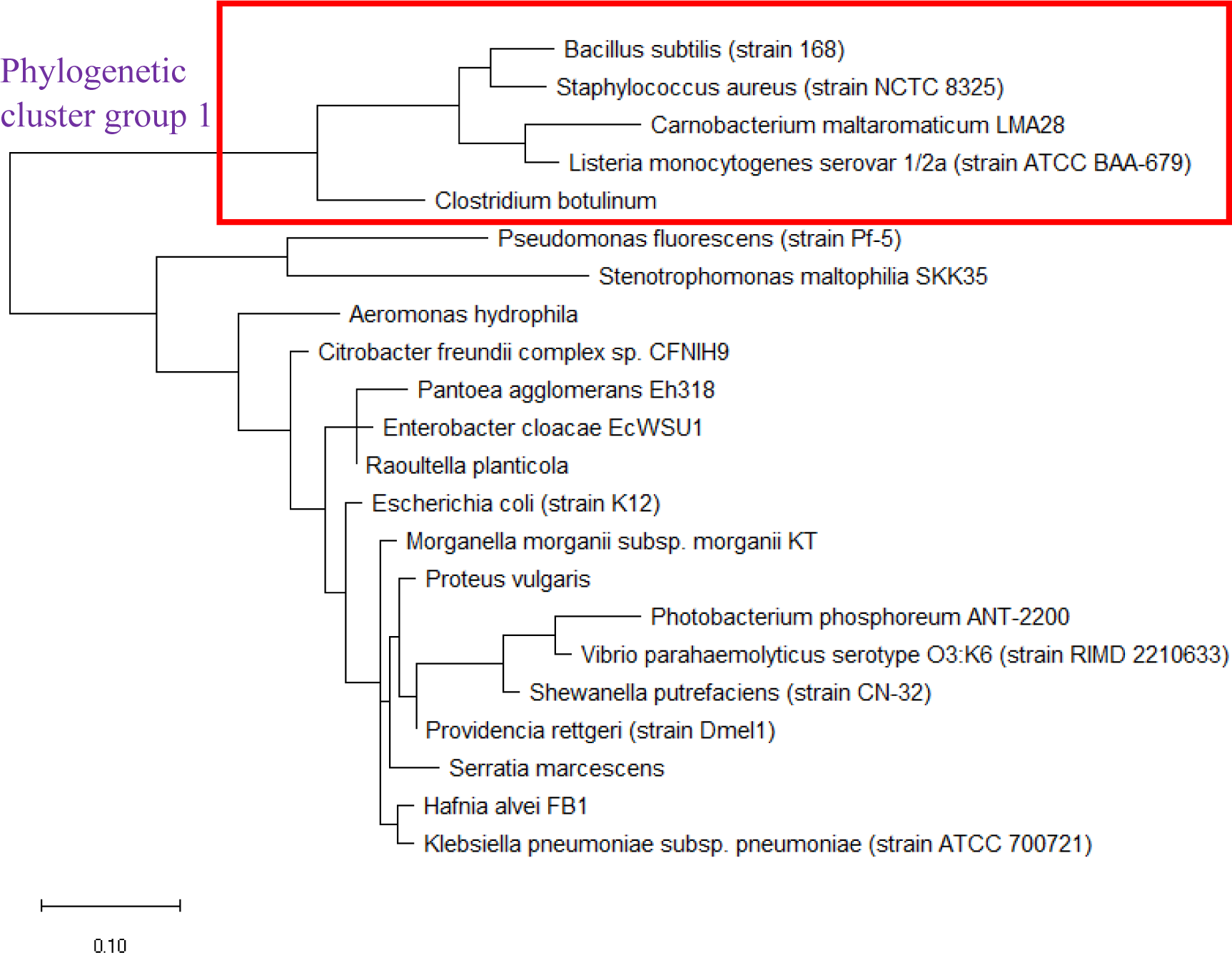
Maximum likelihood phylogenetic tree based on ribosomal protein L27 of selected set of bacterial species.

The maximum likelihood phylogenetic tree based on ribosomal protein L28 of selected set of bacterial species was shown in Figure 10, and it could be seen that the ribosomal protein L28 could replicate two phylogenetic cluster groups found in the phylogenetic tree of 16S rRNA. Thus, ribosomal protein L28’s evolutionary trajectory might have overlapped that of 16S rRNA even though differences could be observed in the two phylogenetic trees. A point to note was that the phylogenetic tree of ribosomal protein L28 could not discern differences in the phylogeny of *Citrobacter freundii, Enterobacter cloacae, Escherichia coli, Klebsiella pneuumoniae and Raoultella planticola*, which suggested that the ribosomal protein L28 of the species were closely-related and might be transferred between species by horizontal gene transfer. Overall, ribosomal protein L28 holds phylogenetic significance.

**Figure 10:**
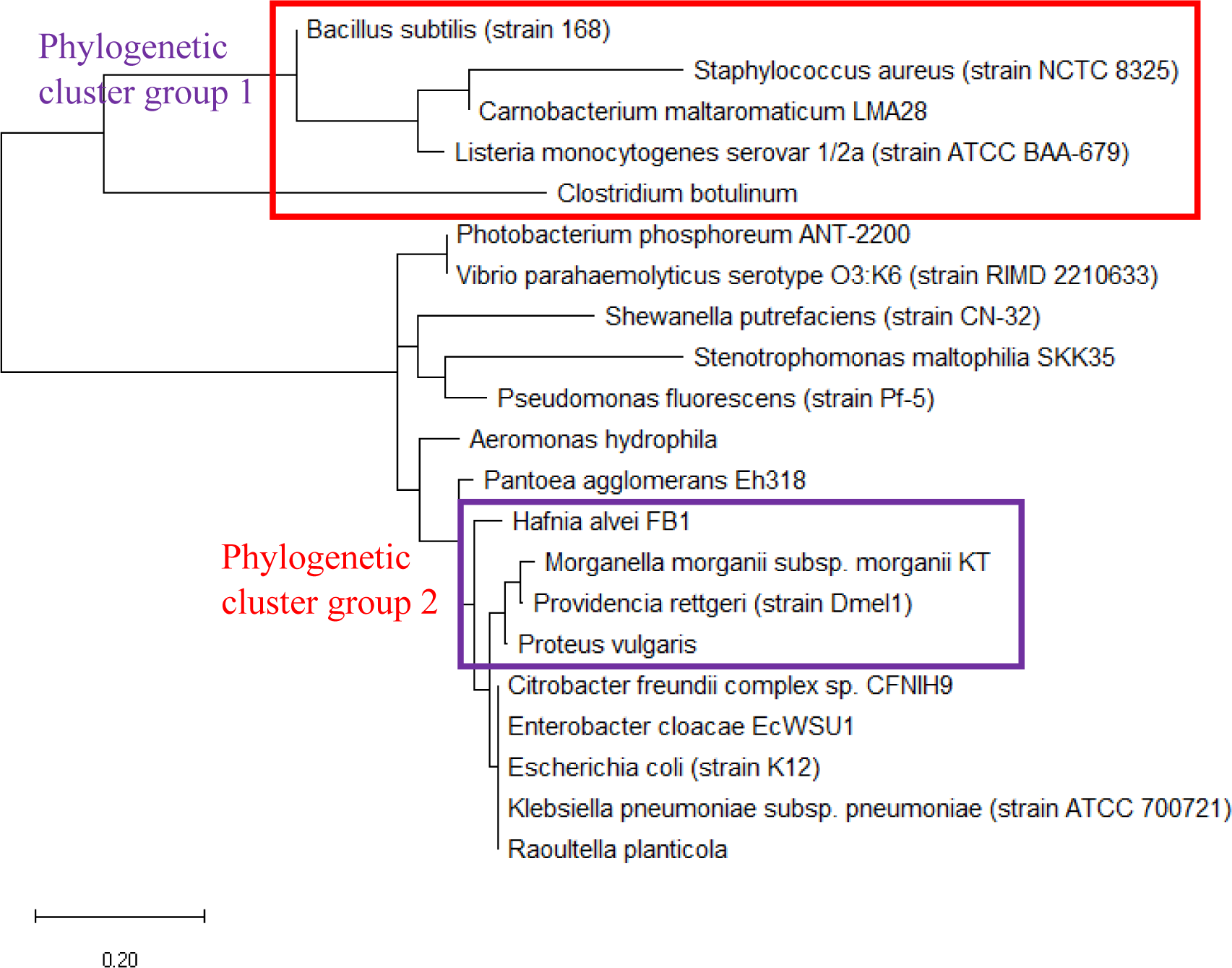
Maximum likelihood phylogenetic tree based on ribosomal protein L28 of selected bacterial species.

Figure 11 shows the maximum likelihood phylogenetic tree of ribosomal protein L29 of a selected set of bacterial species. The data revealed that the phylogeny of ribosomal protein L29 could recapitulate phylogenetic cluster group 1 and 2 of the phylogenetic tree of 16S rRNA. This indicated that significant overlap existed between the phylogenetic trajectories of 16S rRNA and ribosomal protein L29. Thus, ribosomal protein L29 holds phylogenetic significance for the classification of bacterial species but the phylogeny obtained would be different from that of 16S rRNA given the evolutionary divergence between the two biomolecules.

**Figure 11:**
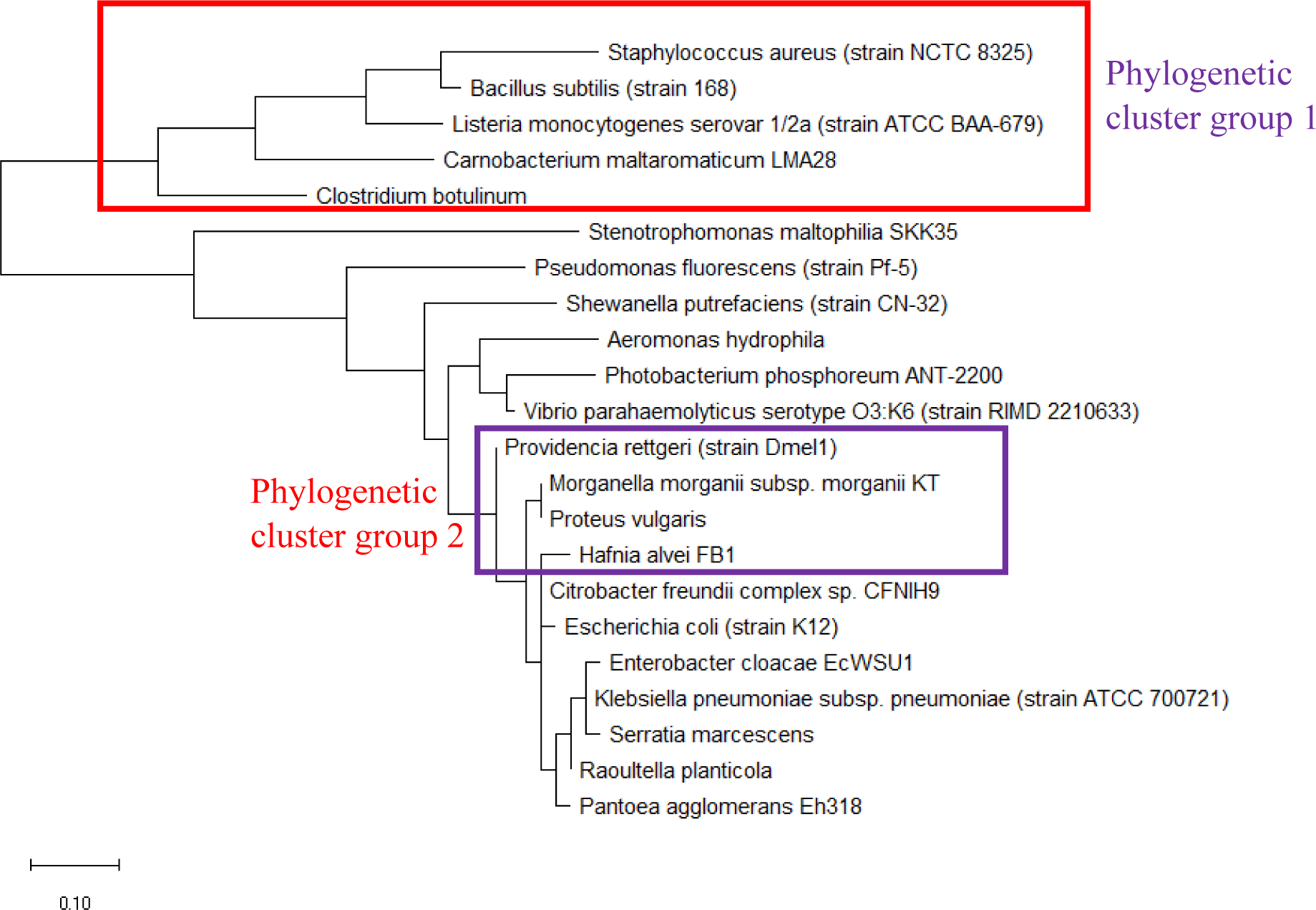
Maximum likelihood phylogenetic tree based on ribosomal protein L29 of selected group of bacterial species.

Figure 12 shows the maximum likelihood phylogenetic tree based on ribosomal protein L30 of selected bacterial species. The data revealed that phylogeny inferred from ribosomal protein L30 sequence recapitulated phylogenetic cluster group 1 and 2 present in the phylogenetic tree of 16S rRNA. Lack of evolutionary distance between the ribosomal protein L30 of *Escherichia coli, Enterobacter cloacae, Citrobacter freundii, Klebsiella pneumoniae*, and *Raoultella planticola* revealed that the ribosomal protein in this species may share a common ancestry protein. This indicated that ribosomal protein L30 holds phylogenetic significance for understanding the evolutionary history of microbial species profiled by MALDI-TOF mass spectrometry.

**Figure 12:**
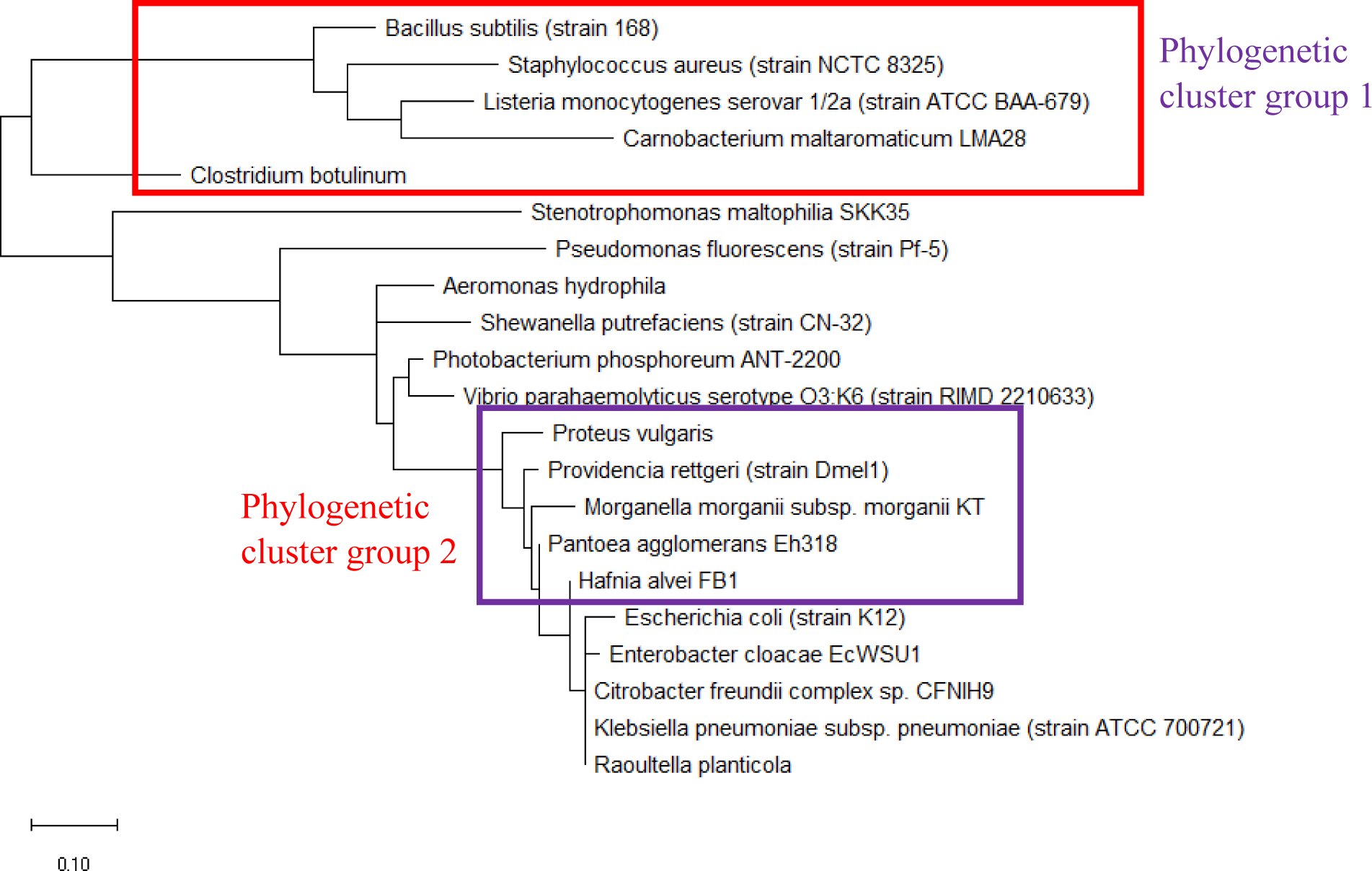
Maximum likelihood phylogenetic tree based on ribosomal protein L30 of selected set of bacterial species.

The maximum likelihood phylogenetic tree based on ribosomal protein L31 of selected set of bacterial species was depicted in Figure 13. The data revealed that the phylogenetic tree of ribosomal protein L31 could not replicate phylogenetic cluster group 1 and 2 of 16S rRNA. This highlighted that ribosomal protein L31 does not hold phylogenetic significance for understanding the evolutionary history of microbial species.

**Figure 13:**
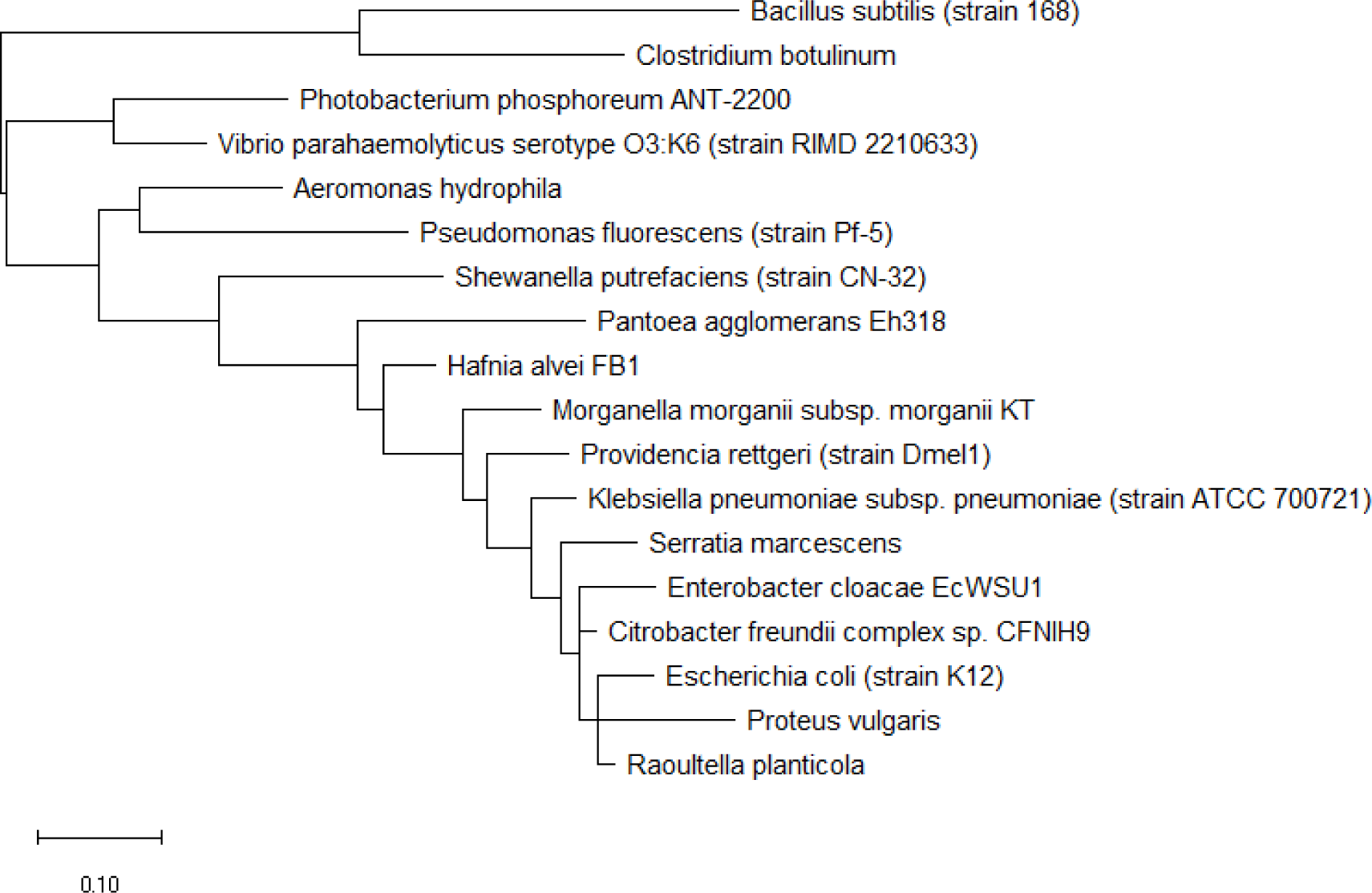
Maximum likelihood phylogenetic tree based on ribosomal protein L31 of selected set of bacterial species.

The maximum likelihood phylogenetic tree based on ribosomal protein L31 Type B of a selected set of bacterial species is presented in Figure 14. The data showed that the phylogeny of ribosomal protein L31 Type B could replicate phylogenetic cluster group 1 of 16S rRNA’s phylogeny. However, phylogenetic cluster group 2 of 16S rRNA’s phylogeny could not be discerned in the phylogenetic tree of ribosomal protein L31 Type B. Overall, ribosomal protein L31 Type B holds phylogenetic significance for understanding the broad evolutionary history of the bacterial species.

**Figure 14:**
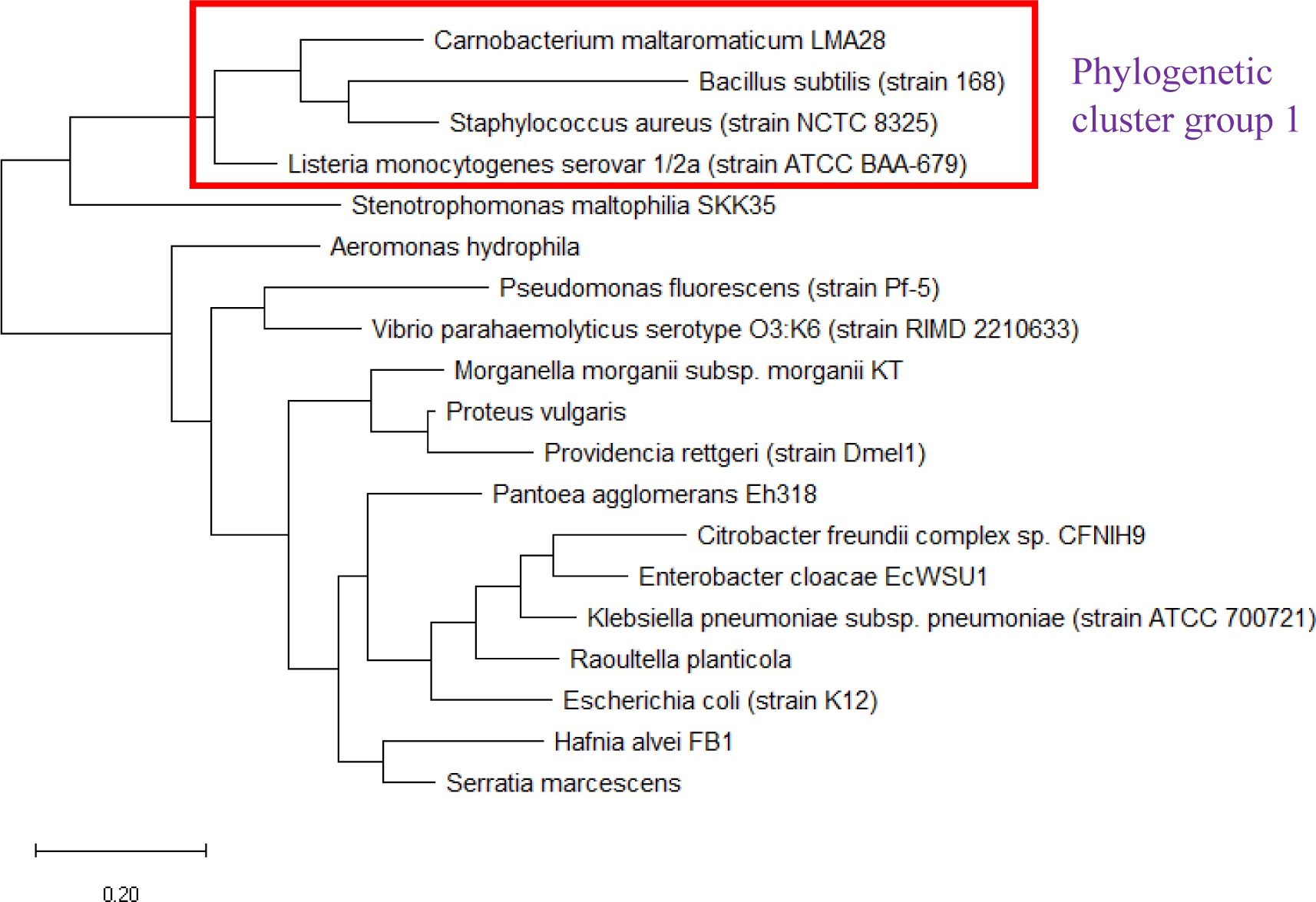
Maximum likelihood phylogenetic tree based on ribosomal protein L31 Type B of selected set of bacterial species.

Figure 15 shows the maximum likelihood phylogenetic tree based on ribosomal protein L32 of selected bacterial species. The date revealed that phylogenetic cluster group 1 of 16S rRNA’s phylogeny could be replicated in the phylogeny of ribosomal protein L32. However, phylogenetic cluster group 2 of 16S rRNA phylogenetic tree was absent in that of ribosomal protein L32. Overall, ribosomal protein L32 holds phylogenetic significance for understanding the evolutionary history of bacterial species.

**Figure 15:**
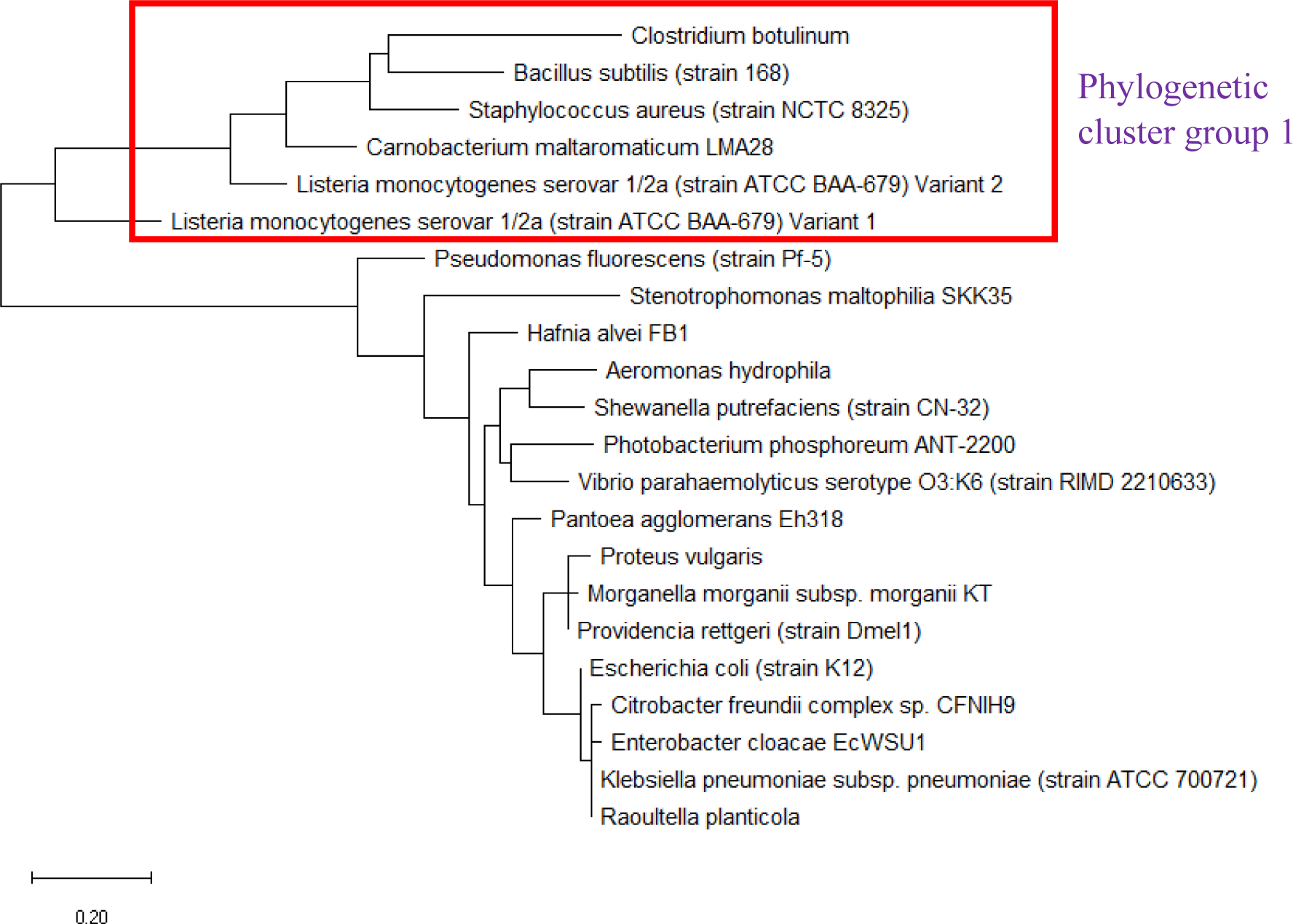
Maximum likelihood phylogenetic tree based on ribosomal protein L32 of selected set of bacterial species.

Maximum likelihood phylogenetic tree based on ribosomal protein L33 of selected set of bacterial species is shown in Figure 16. The data revealed that the phylogeny of ribosomal protein L33 could replicate the phylogenetic cluster group 1 of 16S rRNA’s phylogeny; however, phylogenetic cluster group 2 could not be replicated. Additionally, the phylogenetic tree revealed that the ribosomal protein L33 of *Raoultella planticola, Citrobacter freundii, Enterobacter cloacae*, and *Pantoea agglomerans* were closely-related and highly conserved, suggesting that it might have descended from a common ancestral protein.

**Figure 16:**
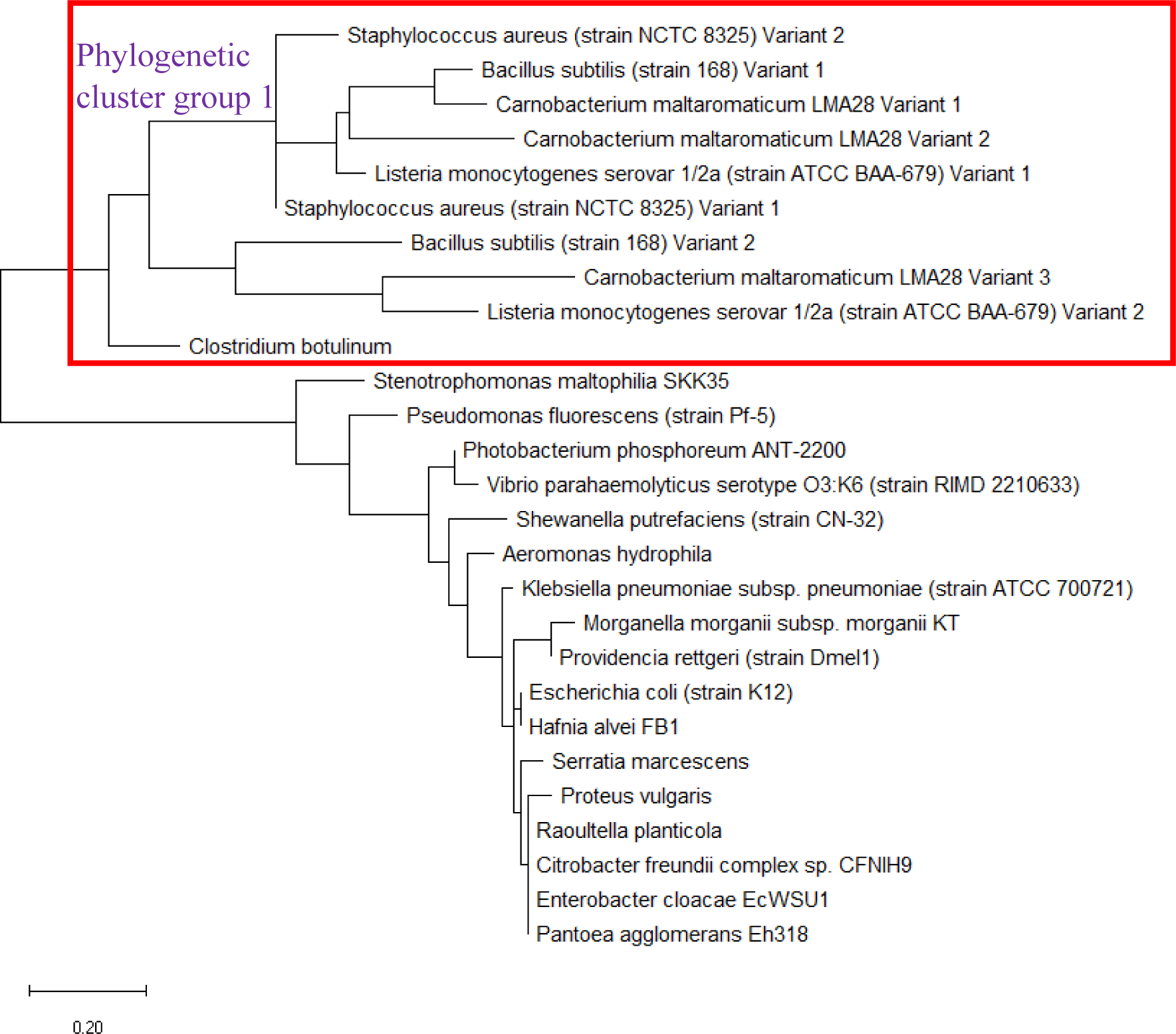
Maximum likelihood phylogenetic tree based on ribosomal protein L33 of selected set of bacterial species.

Figure 17 shows the maximum likelihood phylogenetic tree based on ribosomal protein L34 of selected set of bacterial species. Specifically, the data revealed that phylogenetic cluster group 1 of 16S rRNA’s phylogeny could be replicated in that of ribosomal protein L34. However, phylogenetic cluster group 2 of 16S rRNA’s phylogeny could not be replicated. Additionally, the phylogenetic tree presented revealed that the ribosomal protein L34 of *Enterobacter cloacae, Escherichia coli, Klebsiella pneumoniae, Morganella morganii*, and *Providencia rettgeri* were closely related and likely descended from a common ancestral protein. Taken together, ribosomal protein L34 holds phylogenetic significance.

**Figure 17:**
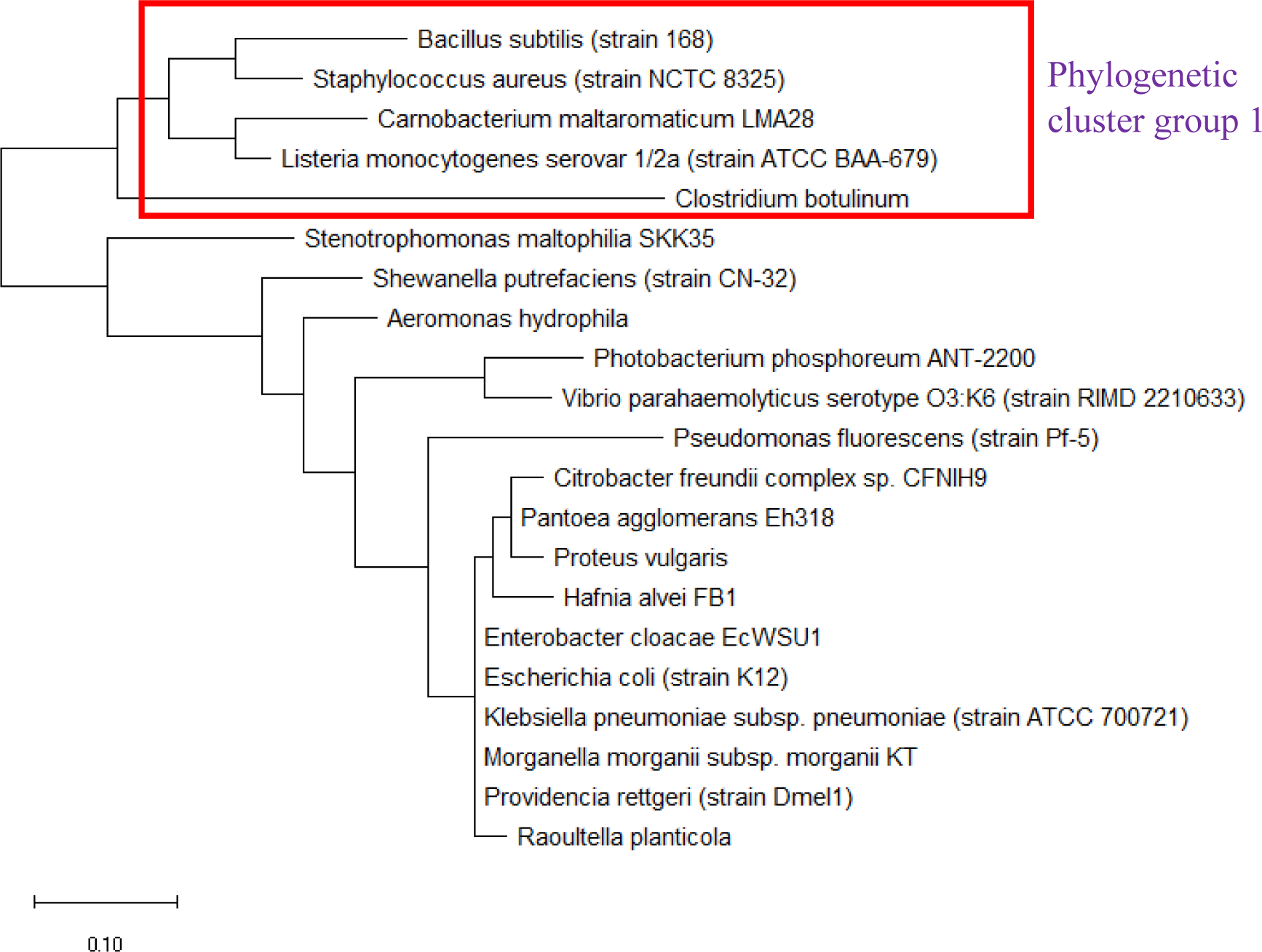
Maximum likelihood phylogenetic tree based on ribosomal protein L34 of selected set of bacterial species.

The maximum likelihood phylogenetic tree based on ribosomal protein L35 of selected set of bacterial species is shown in Figure 18. The data revealed that the phylogenetic tree of ribosomal protein L35 could replicate phylogenetic cluster group 1 and 2 of 16S rRNA’s phylogeny, which indicated that there might be substantial overlap in the evolutionary histories of ribosomal protein L35 and 16S rRNA. Additionally, the phylogenetic tree also revealed that the ribosomal protein L35 of *Escherichia coli, Klebsiella pneumoniae, and Raoultella planticola* are closely-related and highly conserved. The same also applied to ribosomal protein L35 of *Citrobacter freundii*, and *Enterobacter cloacae*, which suggested a common ancestral protein for ribosomal protein L35 in the bacterial species. Overall, ribosomal protein L35 holds phylogenetic significance for understanding the evolutionary trajectories of bacterial species.

**Figure 18:**
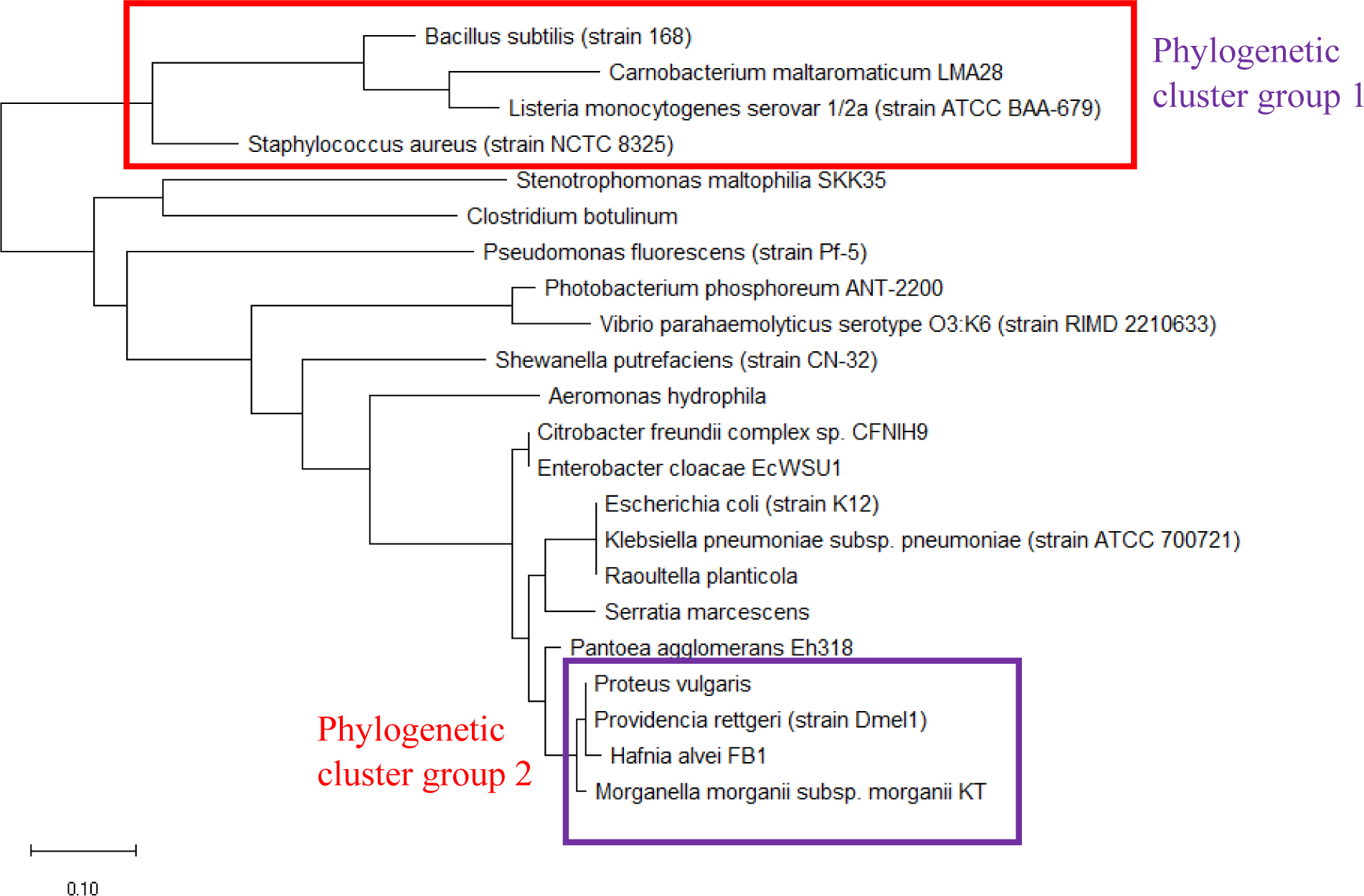
Maximum likelihood phylogenetic tree based on ribosomal protein L35 of selected bacterial species.

Figure 19 shows the maximum likelihood phylogenetic tree based on ribosomal protein L36 of selected set of bacterial species. The data revealed that the phylogeny of ribosomal protein L36 could not reproduce phylogenetic cluster group 2 of 16S rRNA’s phylogeny even though phylogenetic cluster group 1 was reproduced. However, additional species were placed on the branch of the phylogenetic tree on the side of phylogenetic cluster group 1. These were not present in the phylogenetic tree of 16S rRNA. Thus, ribosomal protein L36 does not hold phylogenetic significance for understanding the evolutionary trajectories of bacterial species. This is in contrast to the central role that ribosomal protein L36 plays in maintaining the structural stability of the ribosome large subunit, which suggested that its structure should be highly conserved. However, conservation of protein structure does not require strict conservation of protein amino acid sequence. Hence, divergence in amino acid sequence of ribosomal protein L36 of different bacterial species occurred and was captured by the maximum likelihood phylogenetic tree of ribosomal protein L36.

**Figure 19:**
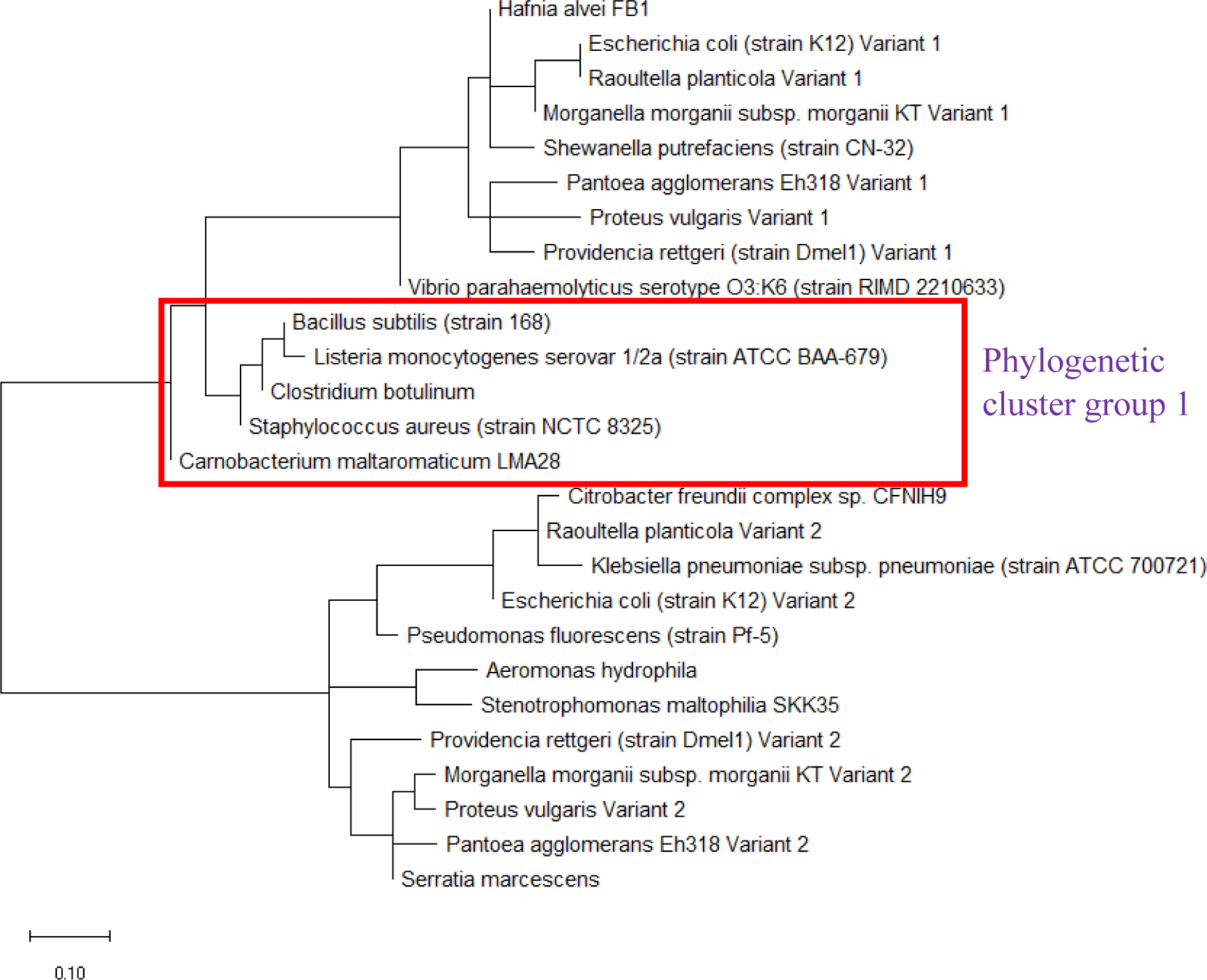
Maximum likelihood phylogenetic tree based on ribosomal protein L36 of a selected set of bacterial species.

Maximum likelihood phylogenetic tree based on ribosomal protein S16 of selected set of bacterial species is shown in Figure 20. The data revealed that the phylogeny of ribosomal protein S16 could replicate phylogenetic cluster group 1 of 16S rRNA’s phylogeny. However, phylogenetic cluster group 2 of 16S rRNA’s phylogeny could not be replicated in the phylogenetic tree of ribosomal protein S16. Overall, ribosomal protein S16 holds phylogenetic significance for understanding the evolutionary history of bacterial species.

**Figure 20:**
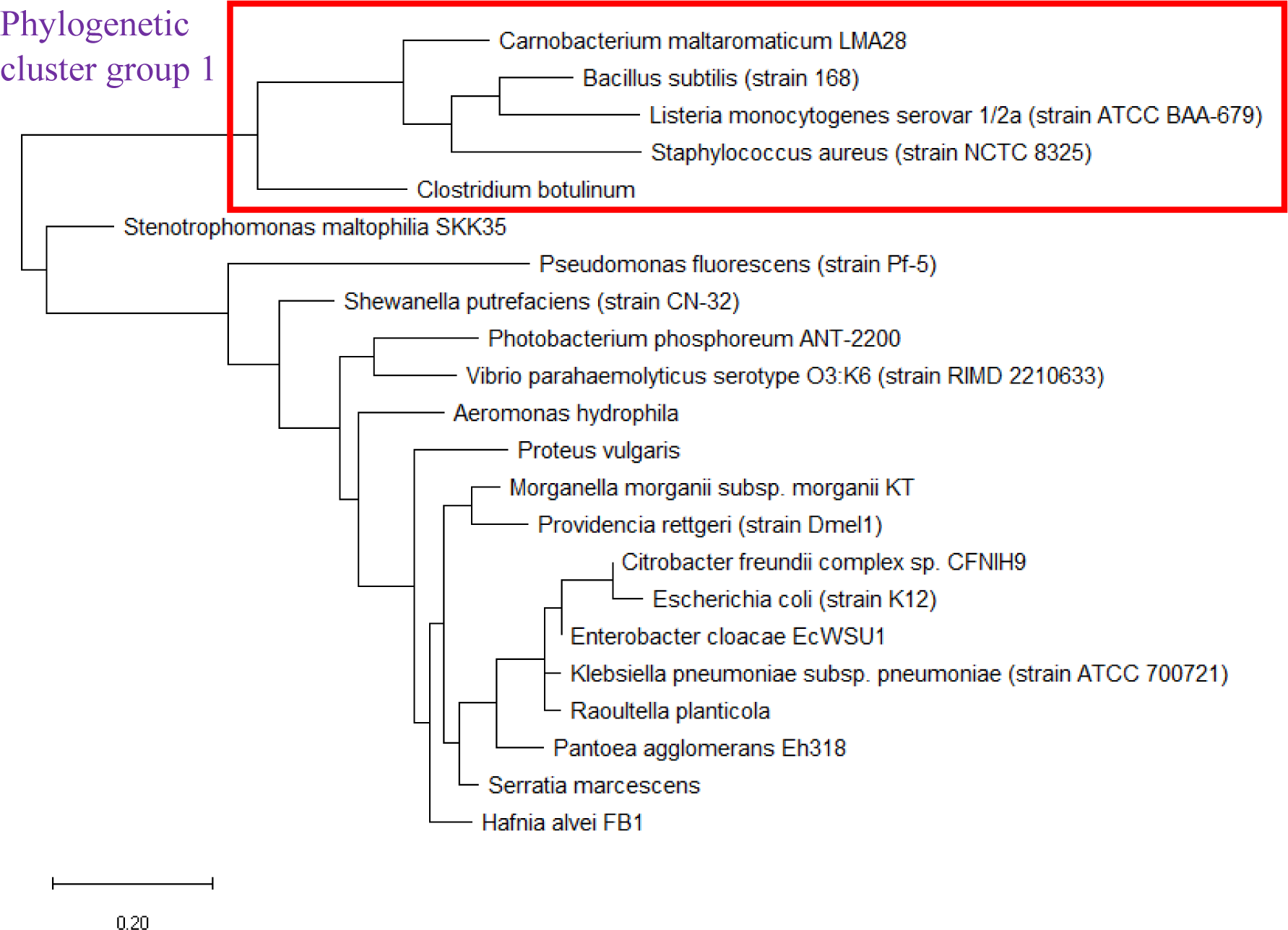
Maximum likelihood phylogenetic tree based on ribosomal protein S16 of selected set of bacterial species.

Figure 21 shows the maximum likelihood phylogenetic tree based on ribosomal protein S17 of selected set of bacterial species. The data revealed that the phylogeny of ribosomal protein S17 could replicate both phylogenetic cluster group 1 and 2 of 16S rRNA’s phylogeny. Overall, ribosomal protein S17 holds phylogenetic significance for understanding the evolutionary trajectories taken by different bacterial species.

**Figure 21:**
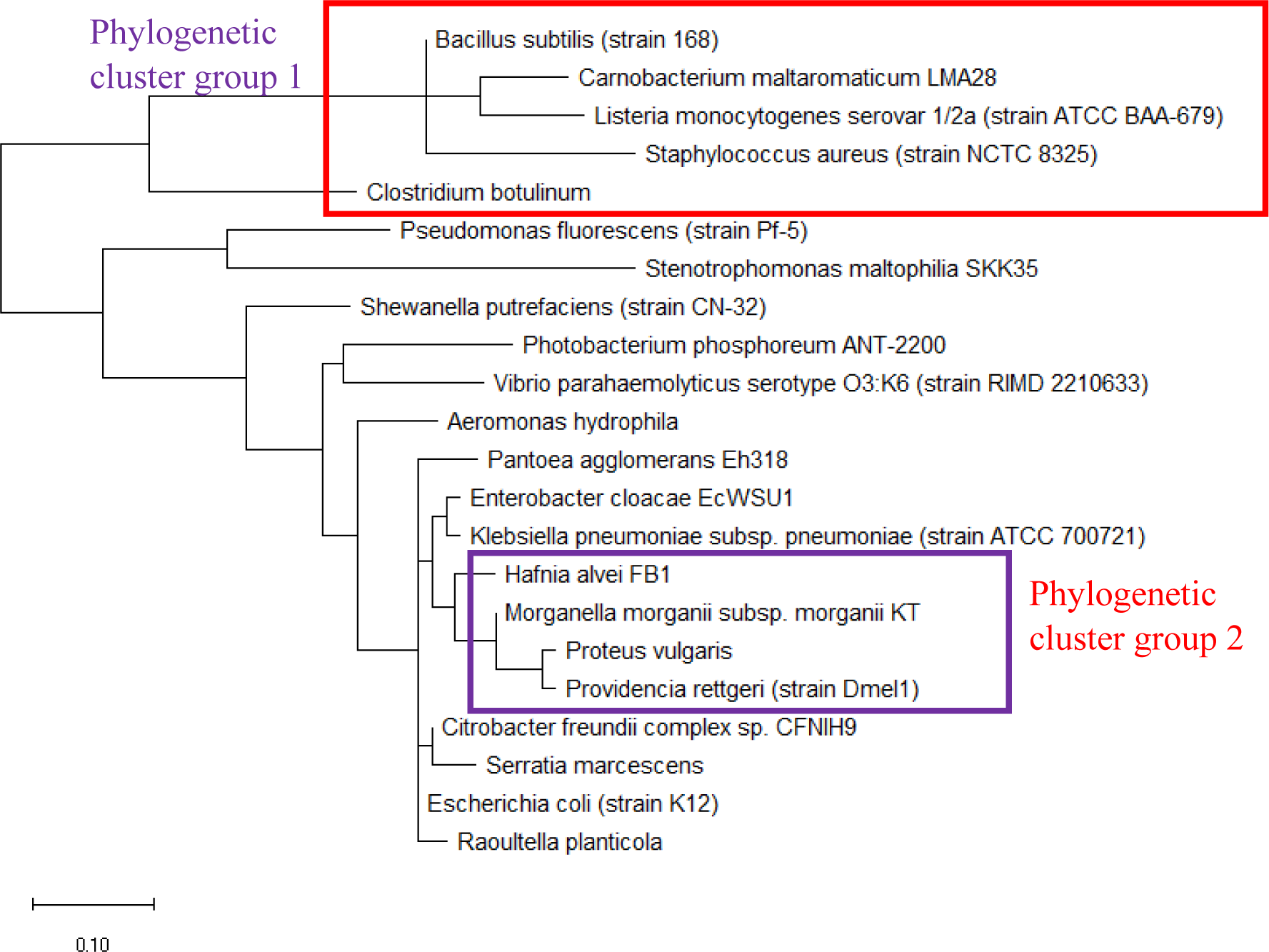
Maximum likelihood phylogenetic tree based on ribosomal protein S17 of selected set of bacterial species.

Maximum likelihood phylogenetic tree based on ribosomal protein S18 of selected set of bacterial species is shown in Figure 22. The data revealed that the phylogenetic tree of ribosomal protein S18 recapitulated the phylogenetic cluster group 1 of 16S rRNA’s phylogeny. However, the ribosomal protein does not hold phylogenetic significance as the phylogenetic tree revealed a set of bacterial species with high conserved amino acid sequence of ribosomal protein S18. These bacterial species were *Serratia marcescens, Citrobacter freundii, Enterobacter cloacae, Escherichia coli, Hafnia alvei, Klebsiella pneumoniae, Proteus vulgaris, Providencia rettgeri*, and *Raoultella planticola.* For a protein to be useful for classifying different species of bacteria, it must retain sufficient sequence diversity to allow the evolutionary relationships between different species to be captured in amino acid alterations. Thus, a protein with a highly conserved amino acid sequence lacks the sequence diversity necessary to endow it with sufficient phylogenetic power to probe the phylogenetic relationships between different bacterial species.

**Figure 22:**
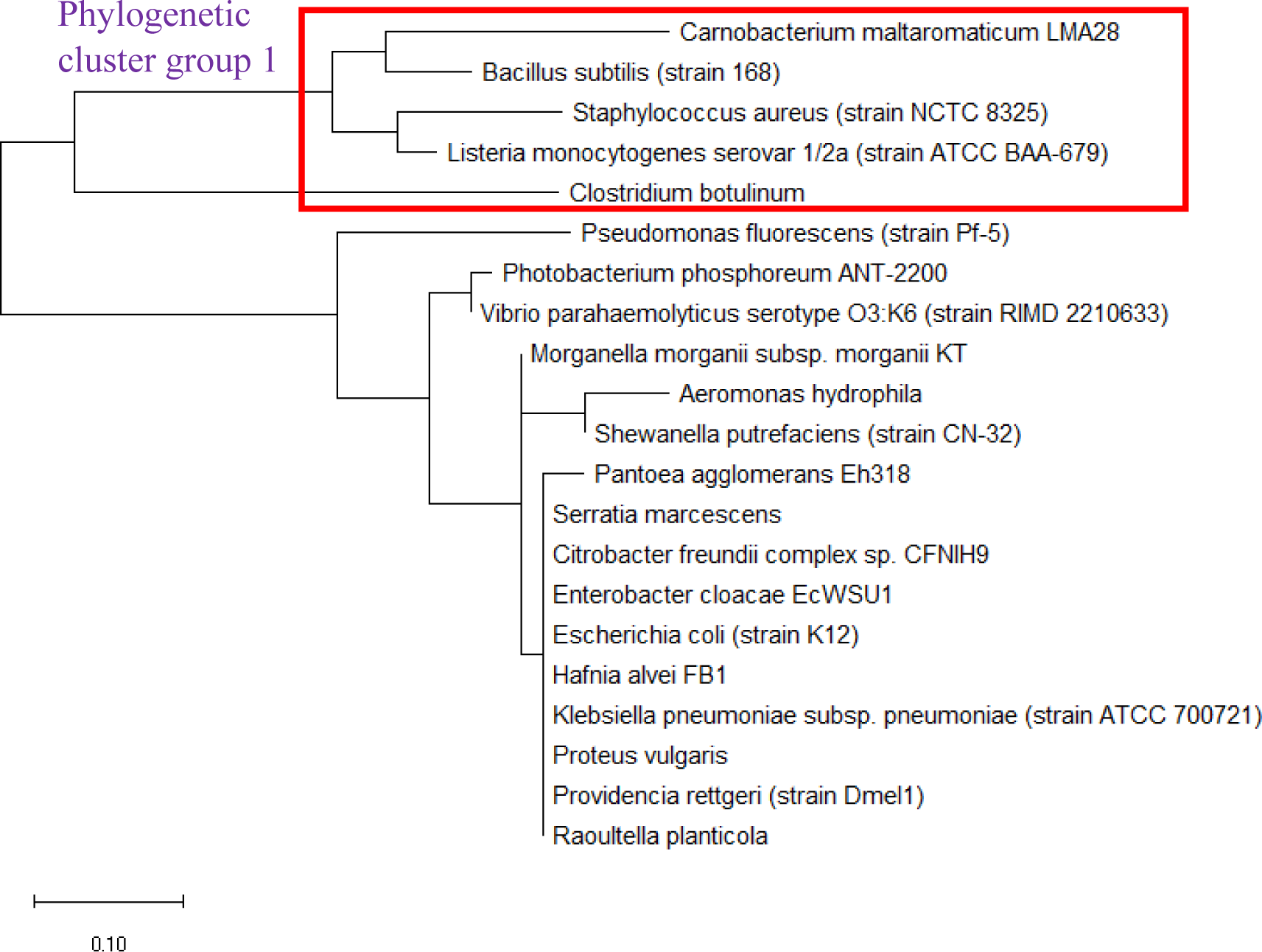
Maximum likelihood phylogenetic tree based on ribosomal protein S18 of selected bacterial species.

Figure 23 shows the maximum likelihood phylogenetic tree based on ribosomal protein S20 of selected set of bacterial species. The data revealed that the phylogeny of ribosomal protein S20 could replicate phylogenetic cluster group 1 of 16S rRNA’s phylogeny. However, phylogenetic cluster group 2 of 16S rRNA’s phylogeny could not be recapitulated in the phylogenetic tree of ribosomal protein S20. Overall, ribosomal protein S20 holds phylogenetic significance for explaining the evolutionary trajectories of different bacterial species.

**Figure 23:**
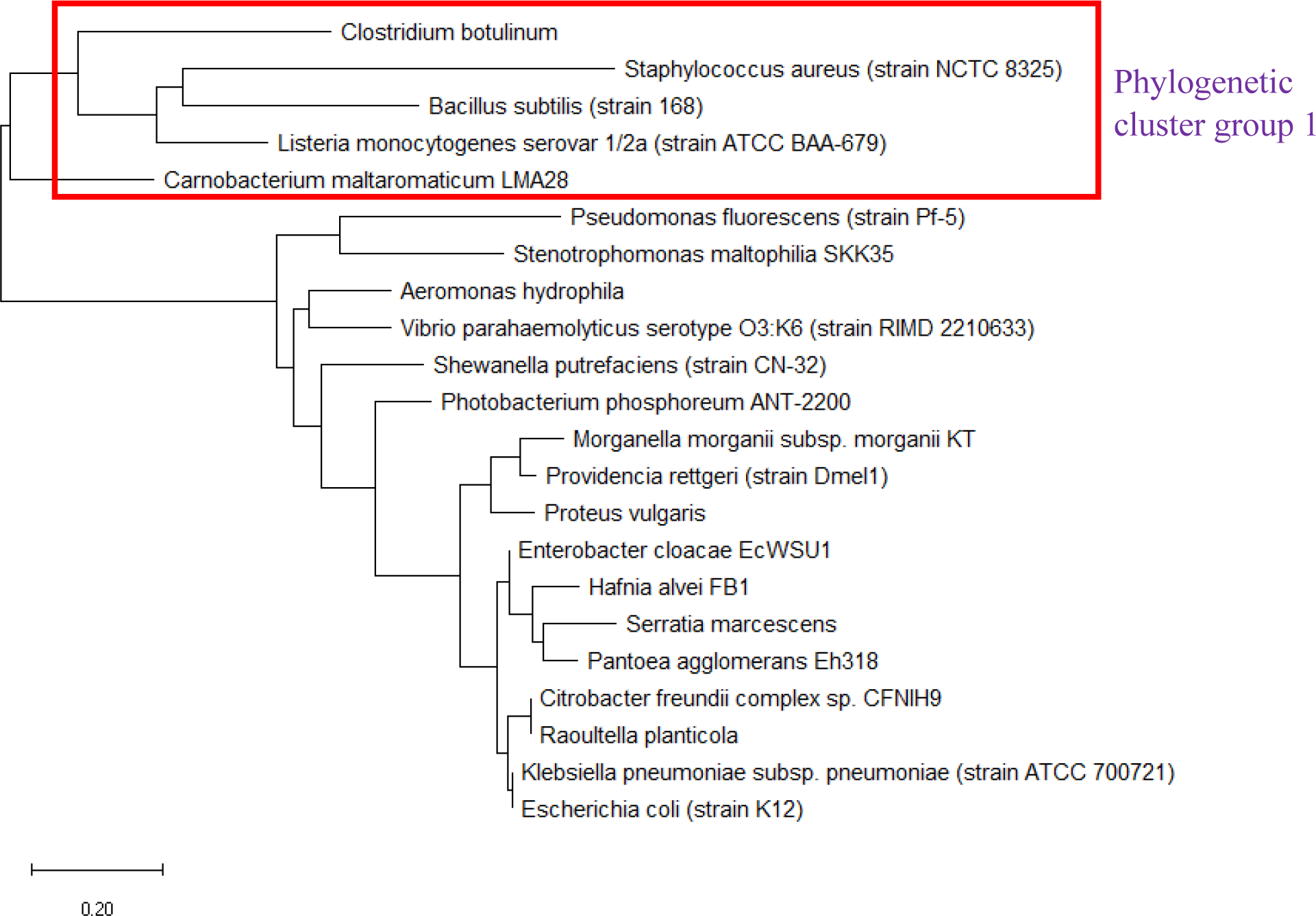
Maximum likelihood phylogenetic tree based on ribosomal protein S20 of selected set of bacterial species.

Maximum likelihood phylogenetic tree based on ribosomal protein S21 of a selected set of bacterial species is shown in Figure 24. The data revealed that the phylogenetic tree of ribosomal protein S21 could replicate phylogenetic cluster group 1 of 16S rRNA’s phylogeny. However, the ribosomal protein does not hold phylogenetic significance as its phylogenetic tree showed a set of bacterial species with closely-related amino acid sequence. Lack of sequence diversity meant that the ribosomal protein could not tracked the evolutionary trajectories of different bacterial species to an extent sufficient to inform classification decisions. The set of bacterial species with closely-related ribosomal protein S21 was *Raoultella planticola, Citrobacter freundii, Enterobacter cloacae, Escherichia coli, Hafnia alvei, Klebsiella pneumoniae, Pantoea agglomerans, Proteus vulgaris*, and *Providencia rettgeri.*

**Figure 24:**
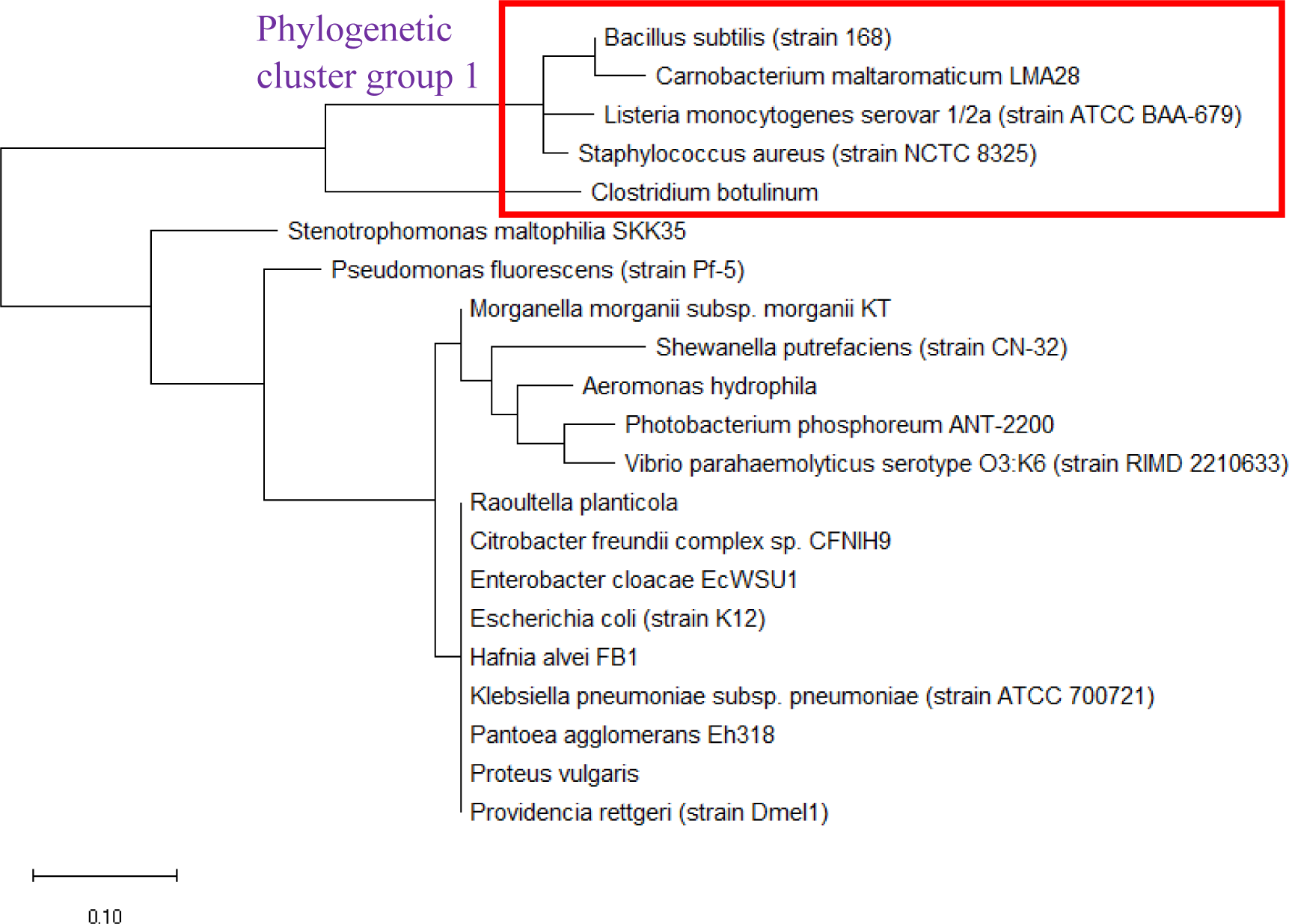
Maximum likelihood phylogenetic tree based on ribosomal protein S21 of selected bacterial species.

### Multi-locus sequence typing

Given that individual ribosomal protein depicts a different evolutionary history between species in the same set of bacterial species, the evolutionary trajectories traversed by individual species could not be accurately depicted by a single ribosomal protein. Thus, the concept of using multiple genes or proteins for understanding the phylogenetic relationships between different species was born.^17^ Specifically, known as multi-locus sequence typing, the approach concatenates the nucleotide or amino acid sequences of multiple genes and proteins to inform on the phylogenetic relationships between species. Used successfully in many phylogenetic studies of bacterial species from different genera, the approach was used in this study to assess if concatenating different ribosomal protein amino acid sequence together would help improve the reconstruction of phylogenetic tree compared to that of 16S rRNA. To this end, amino acid sequence of 6 ribosomal proteins L29, S16, S20, S17, L27 and L35 was concatenated randomly in the order L29-S16-S20-S17-L27-L35 (Figure 25).

**Figure 25:**
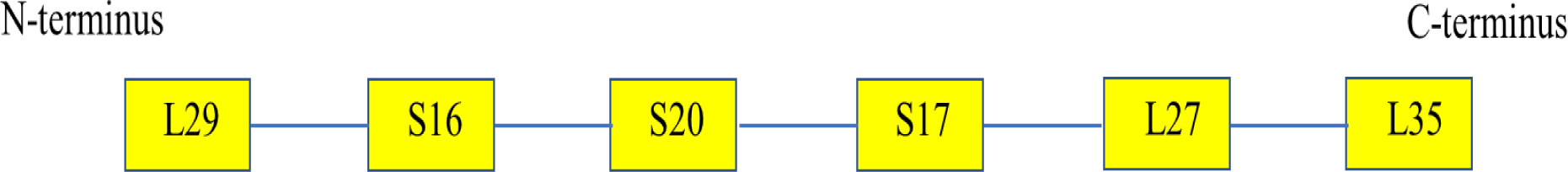
Concatenation of amino acid sequence of different ribosomal proteins for multi-locus sequence typing.

Figure 26 shows the maximum likelihood phylogenetic tree based on the concatenated amino acid sequence of different ribosomal protein used in multi-locus sequence typing of selected set of bacterial species. The data revealed that phylogenetic cluster group 1 and 2 of 16S rRNA’s phylogeny could be reproduced by the phylogeny of the concatenated ribosomal protein amino acid sequence. However, differences in phylogenetic tree structure and placement of individual bacterial species in the tree exist between the phylogenetic tree of 16S rRNA and the concatenated ribosomal protein amino acid sequence. This is to be expected given that different evolutionary histories were chronicled by individual ribosomal protein compared to 16S rRNA gene that in aggregate could not be smoothed over by the effects of amino acid sequence concatenation. Overall, concatenation of ribosomal protein amino acid sequence could explain, broadly, the phylogeny of different bacterial species as compared to the classification engendered from the phylogenetic tree based on 16S rRNA gene sequence of the same species.

**Figure 26:**
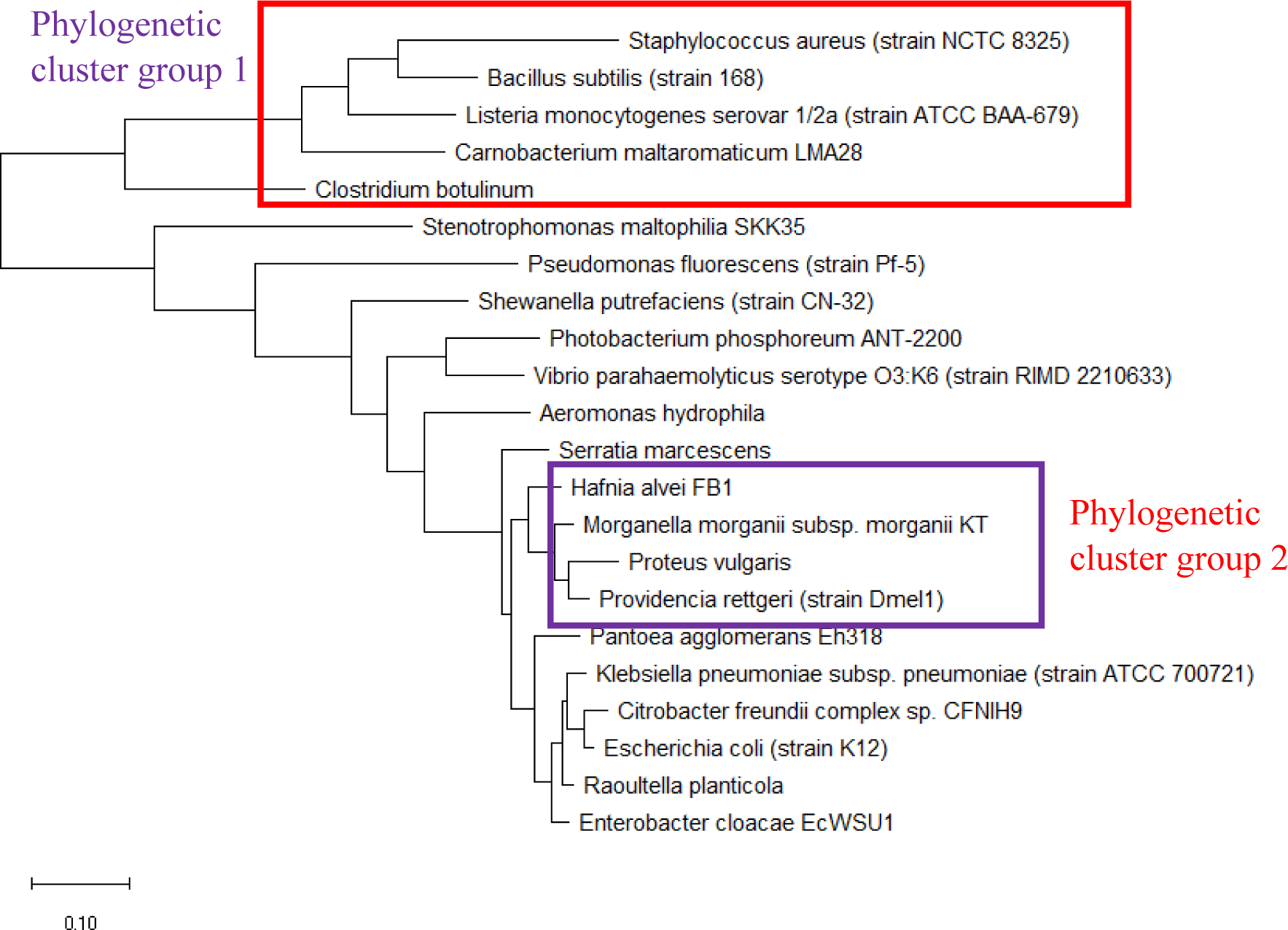
Maximum likelihood phylogenetic tree based on multi-locus sequence typing of concatenated ribosomal protein L29-S16-S20-S17-L27-L35 of selected set of bacterial species.

### Structural analysis of annotated ribosomal protein

Molecular structure of proteins provides a different layer of biological information compared to phylogenetic analysis that help complement the search for phylogenetic significance of individual ribosomal proteins. Specifically, given that only a selected set of ribosomal proteins were annotated in the MALDI-TOF mass spectra of the bacterial species catalogued in the SpectraBank database, an important question revolves around the reasons why a specific group of ribosomal proteins were profiled by the MALDI-TOF mass spectrometer compared to other ribosomal proteins. Could it due to the unique functions of individual ribosomal proteins that confer them a higher relative abundance that facilitated their detection by MALDI-TOF mass spectrometry? Or, was it due to a higher gene dosage of specific ribosomal protein that resulted in higher relative abundance of the ribosomal protein available for mass spectrometric detection? Finally, could molecular structure of the ribosomal proteins inform us of the possible additional functions played by the annotated ribosomal proteins in cell physiology?

To help elucidate the above questions, molecular structure of the annotated ribosomal proteins were obtained with the Phyre^2^ server. Briefly, through a combination of hidden Marknov and homology modelling, the server was able to provide a molecular structure based on a given amino acid sequence. The results of the structural modelling are as shown in Figure 27.

**Figure 27:**
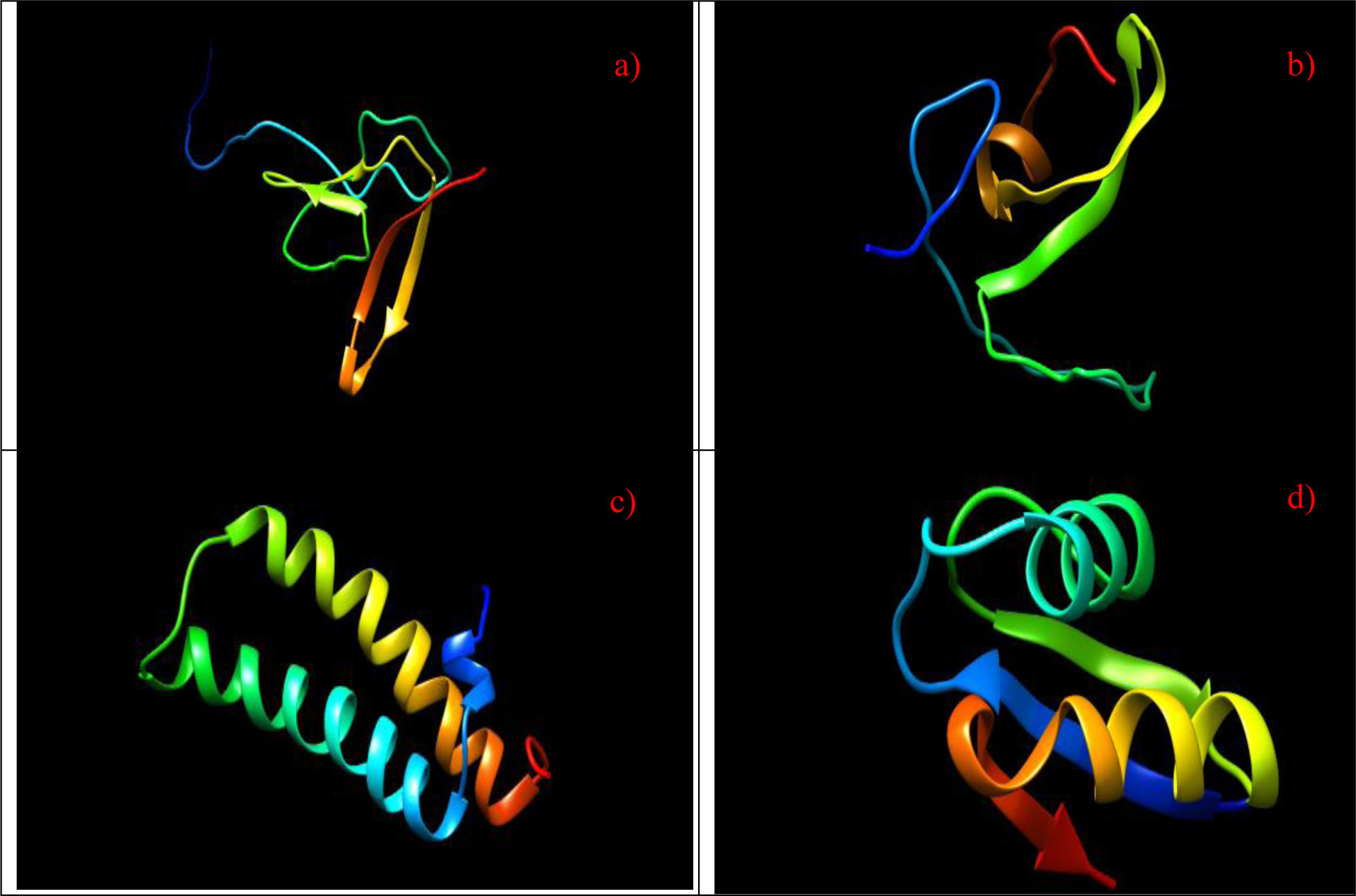
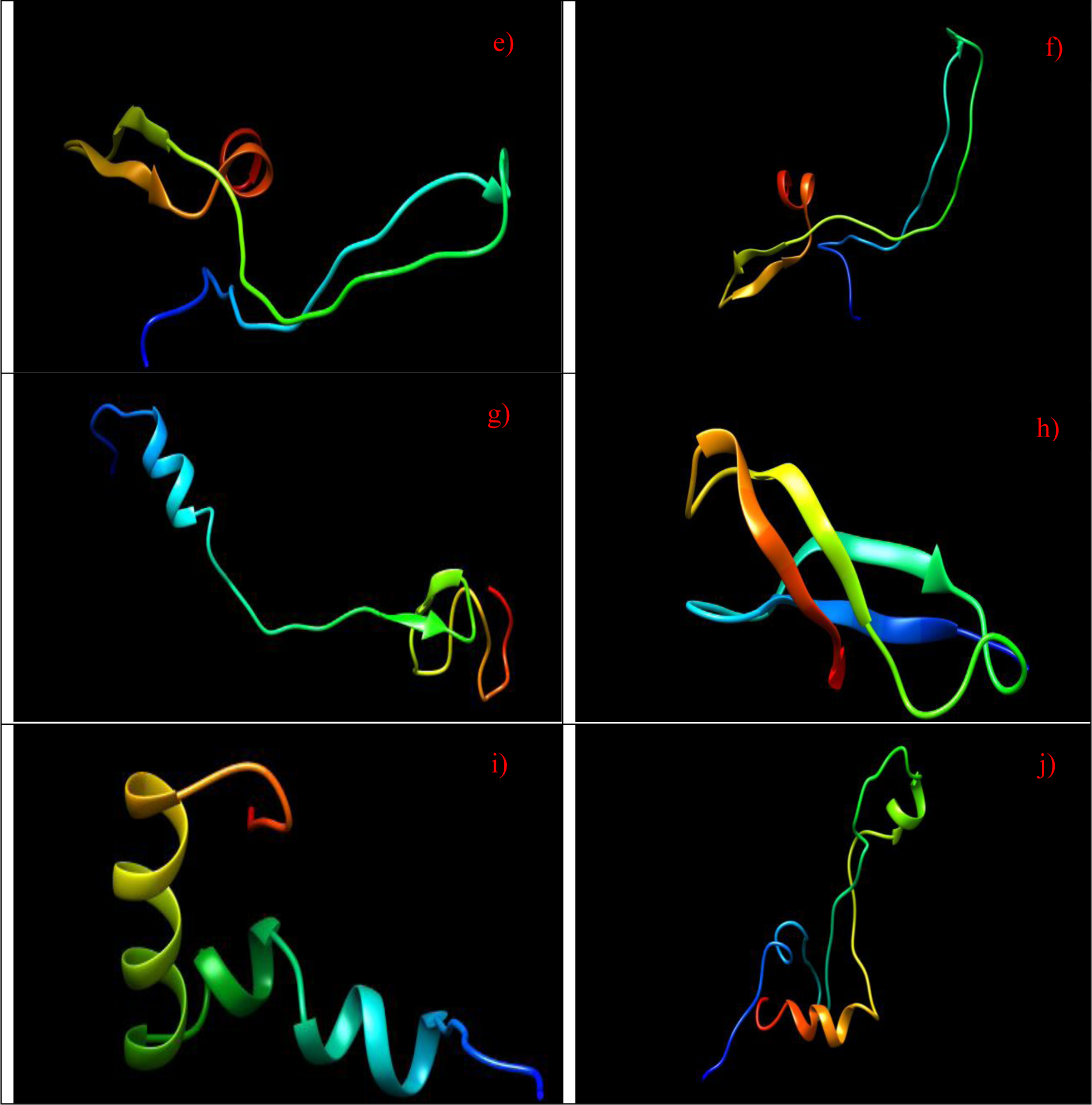
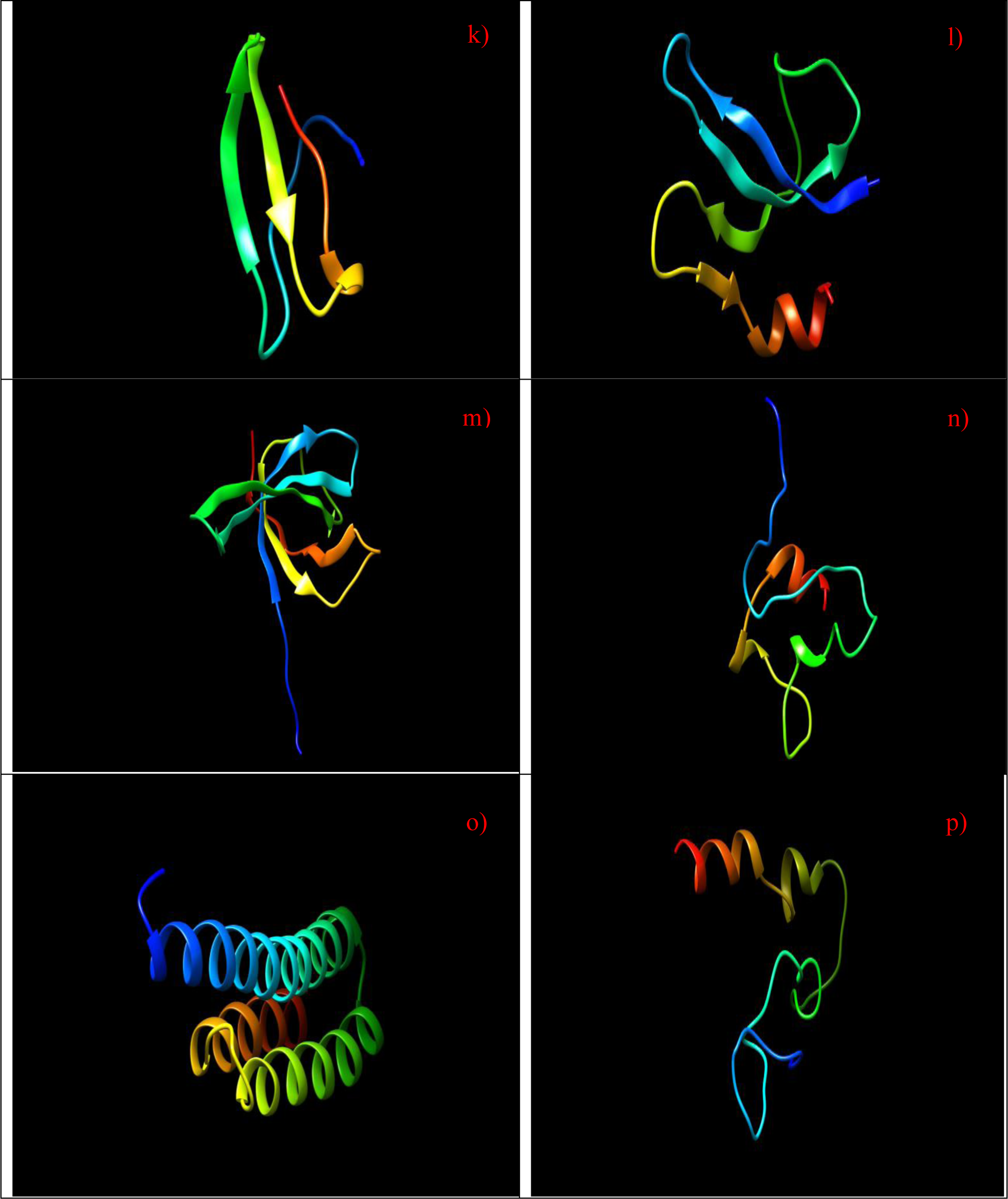
Molecular structure of ribosomal proteins, a) ribosomal protein L27, b) ribosomal protein L28, c) ribosomal protein L29, d) ribosomal protein L30, e) ribosomal protein L31, f) ribosomal protein L31 Type B, g) ribosomal protein L32, h) ribosomal protein L33, i) ribosomal protein L34, j) ribosomal protein L35, k) ribosomal protein L36, l) ribosomal protein S16, m) ribosomal protein S17, n) ribosomal protein S18, o) ribosomal protein S20, p) ribosomal protein S21.

Figure 27 shows the molecular structure of annotated ribosomal proteins. The data revealed that each ribosomal protein had a unique structure except for ribosomal protein L29 and S20 where they shared a similar structure of helixes. However, ribosomal protein L29 had two α-helixes in its structure compared to three α-helixes in ribosomal protein S20. Additionally, the molecular structure of ribosomal protein L31 and L31 Type B were similar probably due to the fact that they are closely-related in sequence. Overall, the annotated ribosomal proteins did not share similar molecular structure, which implied that they did not have similar functions in cellular physiology given that different ribosomal protein played different roles in the ribosome macromolecular complex. This raised the question of why particular ribosomal proteins were more relatively abundant compared to others such that they could be profiled by the MALDI-TOF mass spectrometer.

## Discussion

The question of whether there exists biological basis in the use of MALDI-TOF mass spectrometry for identifying bacterial species was answered in this study through the annotation of ribosomal protein mass peaks in the mass spectra of bacteria. Specifically, many small ribosomal proteins with molecular weight of less than 10000 Da were found to be present in the mass peaks profiled from various bacterial species. Annotated ribosomal proteins were S16, S17, S18, S20, and S21 from the small ribosome subunit, and L27, L28, L29, L30, L31, L31 Type B, L32, L33, L34, L35, and L36 from the large ribosome subunit. However, contrary to previous reports implying that ribosomal proteins could account for many of the mass peaks in MALDI-TOF mass spectra of bacterial species, the current study revealed that only between 1 and 6 mass peaks in each mass spectrum could be annotated by ribosomal proteins. Thus, other classes of proteins such as housekeeping proteins must be investigated for annotating mass peaks in MALDI-TOF mass spectra of bacterial species. Overall, ribosomal protein L36 and L29 accounted for the most number of ribosomal protein mass peak annotation. Other ribosomal proteins with large number of mass peak annotations were L34, L33 and L31. The underlying reasons accounting for the observed annotation frequency of ribosomal proteins remain unknown.

Since ribosomal proteins are highly conserved given their important functional and structural roles in the ribosome macromolecular complex, possible phylogenetic significance of the annotated ribosomal proteins were sought in comparison with that based on 16S rRNA gene. Using maximum likelihood phylogenetic tree as readout, the data revealed that except for ribosomal protein L34, L31, L36 and S18, all other ribosomal proteins showed phylogenetic potential. Specifically, the measurement yardstick was based on whether the maximum likelihood phylogenetic tree based on the ribosomal protein could replicate phylogenetic cluster groups 1 and 2 of 16S rRNA’s phylogeny of the same set of bacterial species. In thinking about phylogenetic significance, it is important to remember that the phylogenetic tree based on different genes and proteins are likely to be different. Thus, what is of importance to conferment of phylogenetic significance to particular ribosomal protein lies in the observation of whether they are able to place different phylogenetic cluster groups correctly in the phylogenetic tree as well as the existence of bacterial species with closely-related ribosomal protein amino acid sequence that defied classification.

To test whether the combined phylogenetic potential of different ribosomal proteins could replicate the phylogeny depicted by 16S rRNA gene, the approach of multi-locus sequence typing was used to concatenate the amino acid sequence of different ribosomal protein together for understanding how the different evolutionary trajectories of the ribosomal proteins could answer the phylogenetic question in a combinatorial manner. Ribosomal protein used in this analysis were L29, S16, S20, S17, L27 and L35. The results revealed that the multi-locus sequence typing approach could replicate phylogenetic cluster group 1 and 2 of 16S rRNA’s phylogeny.

Finally, to understand whether the annotated ribosomal proteins share similar molecular structure and thus functions that could account for why they were in high relative abundance in the cell, the molecular structure of the various annotated ribosomal proteins were modelled using the Phyre^2^ server. Except for close similarity between the molecular structure of ribosomal protein L31 and L31 Type B as well as that between L29 and S20, no close structural similarity was observed for the annotated ribosomal proteins which suggested that they did not share functions beyond their designated roles in the ribosome macromolecular complex. Thus, it remains a mystery why particular ribosomal proteins were in high relative abundance that enable them to be profiled by the MALDI-TOF mass spectrometer.

## Conclusions

Understanding the biological basis inherent in MALDI-TOF mass spectrometry-based microbial identification requires the annotation of mass peaks profiled. This study provided confirmatory evidence that ribosomal proteins could be annotated in the MALDI-TOF mass spectra of 110 bacterial species and strains catalogued in the SpectraBank database. Specifically, annotated ribosomal proteins were small, low molecular weight ribosomal proteins with molecular mass of <10000 Da. Annotated ribosomal proteins were S16, S17, S18, S20, S21 of the small ribosome subunit, and L27, L28, L29, L30, L31, L31 Type B, L32, L33, L34, L35 and L36 of the large ribosome subunit. However, number of annotated ribosomal protein mass peaks were between 1 and 6 per mass spectrum, which was significantly lower than that implied by previous studies linking ribosomal proteins importance to MALDI-TOF MS microbial identification.

To understand the phylogenetic significance of the annotated ribosomal proteins, maximum likelihood phylogenetic trees were reconstructed and compared to that of 16S rRNA. Given that phylogenetic trees based on individual gene or protein would differ depending on the evolutionary trajectory chronicled by the specific biomolecule, reproduction of phylogenetic cluster groups in the phylogenetic tree was taken as readout for phylogenetic significance of the ribosomal protein. Results obtained revealed that except for ribosomal protein L34, L31, L36 and S18, all other annotated ribosomal proteins hold phylogenetic significance. More importantly, concatenation of different ribosomal proteins’ amino acid sequence (L29, S16, S20, S17, L27 and L35) in a multi-locus sequence typing approach also led to the reconstruction of a phylogenetic tree that reproduced major phylogenetic cluster groups of 16S rRNA’s phylogeny.

Finally, structural analysis of the annotated ribosomal proteins did not identify conserved molecular structure of the ribosomal proteins, which implied that they did not share functional similarity. This is understandable given the specific roles and functions played by individual annotated ribosomal proteins in the ribosome macromolecular complex. However, the question of why only specific subset of ribosomal proteins were annotated remain unanswered.

Overall, this study confirmed that small, low molecular weight ribosomal proteins could annotate mass peaks in MALDI-TOF mass spectra of significant number of bacterial species across major bacterial genera. This helps provide the biological basis for MALDI-TOF MS microbial identification. Furthermore, detection of phylogenetic significance of the annotated ribosomal proteins lend further credence to the use of MALDI-TOF mass spectrometry for identification and classification of different bacterial species. Future work should seek to verify if other classes of housekeeping proteins could annotate mass peaks in MALDI-TOF mass spectra of bacterial species.

## Supplementary materials

Annotation of ribosomal protein mass peaks in different species and strains are provided in the accompanying supplementary materials file of this manuscript.

