## Supplementary file for "Annotation of ribosomal protein mass peaks in MALDI-TOF mass spectra of bacterial species and their phylogenetic significance"

Wenfa Ng

Department of Chemical and Biomolecular Engineering, National University of Singapore,

### Annotation of ribosomal protein mass peaks in mass spectra of bacterial species

| Table S1: *Bacillus subtilis* Peak list of ATCC 6051 | Ribosomal Protein |
|---|---|
| 2110.46 | |
| 2181.26 | |
| 2452.48 | |
| 2747.22 | |
| 2786.25 | |
| 3048.53 | |
| 3116.26 | |
| 3342.89 | |
| 3385.04 | |
| 3728.02 | |
| 3860.63 | |
| 3880.77 | |
| 3890.46 | |
| 3902.52 | |
| 3992.31 | |
| 4114.31 | |
| 4245.41 | |
| 4307.68 | L36 |
| 4571.65 | |
| 4945.85 | |
| 5005.3 | |
| 5033 | |
| 6462.61 | |
| 6508.64 | |
| 6602.84 | |
| 6678.29 | |
| 6942.02 | |
| 7715.52 | L29 |
| 9137.74 | |
| 9887.74 | |

Table S1 shows the annotation of ribosomal protein mass peaks in mass spectrum of *Bacillus subtilis* ATCC 6051. Specifically, mass peak 4307.68 Da could be mapped to ribosomal protein L36 and mass peak 7715.52 Da could be mapped to ribosomal protein L29.

| **Table S2: *Bacillus subtilis* Peak list of ATCC 9524** | **Ribosomal Protein** |
|---|---|
| 2106.9 | |
| 2177.85 | |
| 2447.67 | |
| 2743.91 | |
| 2782.81 | |
| 3045.36 | |
| 3113.51 | |
| 3339.17 | |
| 3382.6 | |
| 3725.43 | |
| 3858.19 | |
| 3878.23 | |
| 3989.81 | |
| 4111.57 | |
| 4242.63 | |
| 4305.27 | L36 |
| 4570.29 | |
| 4944.12 | |
| 5005.48 | |
| 5031.26 | |
| 6461.69 | |
| 6507.43 | |
| 6599.98 | |
| 6678.27 | |
| 6942.26 | |
| 7713.99 | L29 |
| 9137.77 | |
| 9887.12 | |

Table S2 shows the mass peaks in *B. subtilis* ATCC 9524 that could be mapped to ribosomal proteins. Specifically, mass peak 4305.27 Da could be mapped to ribosomal protein L36 and that at 7713.99 Da could be mapped to ribosomal protein L29.

| **Table S3:** ***Bacillus subtilis*** **Peak list of ATCC 11774** | **Ribosomal Protein** |
|---|---|
| 2184.81 | |
| 2448.2 | |
| 2744.77 | |
| 3045.96 | |
| 3339.56 | |
| 3645.39 | |
| 3725.69 | |
| 3858.81 | |
| 3878.63 | |
| 4304.95 | L36 |
| 4584.24 | |
| 4944.62 | |
| 5004.96 | |
| 5030.93 | |
| 5313.15 | |
| 6505.67 | |
| 6601.36 | |
| 6674.42 | |
| 6939.97 | |
| 7485.47 | |
| 7712.4 | L29 |
| 9161.75 | |
| 9886.38 | |

Table S3 shows the annotation of ribosomal protein mass peaks in the MALDI-TOF MS mass spectrum of *B. subtilis* ATCC 11774. Ribosomal protein L36 could be mapped to mass peak 4304.95 Da, and ribosomal protein L29 could be mapped to mass peak 7712.4 Da.

| **Table S4:** ***Bacillus subtilis*** **Peak list of Proc6a** | **Ribosomal Protein** |
|---|---|
| 2056.26 | |
| 2178.62 | |
| 2536 | |
| 2744.86 | |
| 2783.85 | |
| 3017.88 | |
| 3045.68 | |
| 3584.46 | |
| 3725.12 | |
| 3857.79 | |
| 3887.61 | |
| 4042.07 | |
| 4111.42 | |
| 4305.23 | L36 |
| 4422.18 | |
| 4568.91 | |
| 4943.22 | |
| 5002.23 | |
| 5030.71 | |
| 5193.56 | |
| 5312.27 | |
| 5572.32 | |
| 5900.65 | L33 |
| 6506.64 | |
| 6599.97 | |
| 6675.74 | |
| 7192.01 | |
| 7713.64 | L29 |
| 9137.45 | |
| 9887.04 | |

Table S4 shows the annotated mass peaks of *B. subtilis* Proc6a. Specifically, ribosomal protein L36 mapped to mass peak 4305.23 Da, ribosomal protein L33 mapped to mass peak 5900.65 Da, and ribosomal protein L29 mapped to mass peak 7713.64 Da.

| **Table S5:** ***Bacillus subtilis*** **Peak list of Proc6b** | **Ribosomal Protein** |
|---|---|
| 2107.65 | |
| 2148.8 | |
| 2186.56 | |
| 2286.02 | |
| 2414.96 | |
| 2505.13 | |
| 2545.96 | |
| 2617.39 | |
| 2744.97 | |
| 2975.33 | |
| 3013.58 | |
| 3045.83 | |
| 3221.95 | |
| 3340.82 | |
| 3353.36 | |
| 3387.4 | |
| 3411.71 | |
| 3483.27 | |
| 3585.21 | |
| 3888.25 | |
| 4082.25 | |
| 4304.96 | L36 |
| 4736.76 | |
| 4944.11 | |
| 5003.95 | |
| 5223.29 | |
| 5572.83 | |
| 5900.92 | L33 |
| 6507.17 | |
| 6598.24 | |
| 6677.98 | |
| 7717.92 | L29 |
| 9467.35 | |
| 9889.34 | |

Table S5 shows the ribosomal protein mass peaks of *B. subtilis* Proc6b. Specifically, ribosomal protein L36, L33 and L29 could be mapped to mass peak 4304.96 Da, 5900.92 Da, and 7717.92 Da, respectively.

| **Table S6:** ***Bacillus subtilis*** **Peak list of Proc6c** | **Ribosomal Protein** |
|---|---|
| 2055.72 | |
| 2184.49 | |
| 2458.36 | |
| 2515.39 | |
| 2784.04 | |
| 3017.45 | |
| 3045.57 | |
| 3230.24 | |
| 3253.43 | |
| 3725.33 | |
| 3857.6 | |
| 4039.8 | |
| 4304.81 | L36 |
| 4569.42 | |
| 5002.52 | |
| 5031.55 | |
| 5194.43 | |
| 5313.06 | |
| 6461.93 | |
| 6507.76 | |
| 7717.3 | L29 |
| 9140.06 | |
| 9600.82 | S20 |
| 9893.69 | |

Table S6 shows the ribosomal protein mass peaks of *B. subtilis* Proc6c. Specifically, ribosomal protein L36, L29 and S20 could be mapped to mass peaks 4304.81 Da, 7717.3 Da, and 9600.82 Da, respectively.

| ***Table S7: Bacillus subtilis*** **Peak list of Proc721** | **Ribosomal Protein** |
|---|---|
| 2056.44 | |
| 2152.39 | |
| 2178.28 | |
| 2183.8 | |
| 2536.5 | |
| 2545.22 | |
| 2744.84 | |
| 2783.4 | |
| 2950.14 | |
| 3045.5 | |
| 3253.73 | |
| 3298.83 | |
| 3339.31 | |
| 3584.9 | |
| 3724.56 | |
| 3857.61 | |
| 4111.5 | |
| 4305.07 | L36 |
| 4421.11 | |
| 4734.85 | |
| 4944.11 | |
| 5004.97 | |
| 5223.97 | |
| 5573.08 | |
| 5901.3 | L33 |
| 6507.39 | |
| 6599.23 | |
| 6678.02 | |
| 7193.28 | |
| 7714.95 | L29 |

Table S7 shows the ribosomal protein mass peaks of *B. subtilis* Proc 721. Specifically, ribosomal protein L36, L33 and L29 could be mapped to mass peaks 4305.07 Da, 5901.3 Da, and 7714.95 Da, respectively.

| **Table S8:** ***Bacillus subtilis*** **Peak list of ProcB6c** | **Ribosomal Protein** |
|---|---|
| 2056.95 | |
| 2185.51 | |
| 2515.19 | |
| 2744.55 | |
| 2783.73 | |
| 3016.81 | |
| 3044.88 | |
| 3337.15 | |
| 3723.22 | |
| 3855.05 | |
| 3886.02 | |
| 4036.76 | |
| 4302.22 | L36 |
| 4565.05 | |
| 4996.02 | |
| 5026.62 | |
| 5188.94 | |
| 5307.83 | |
| 5567.76 | |
| 5895.12 | L33 |
| 6452.27 | |
| 6499.38 | |
| 6593.83 | |
| 6670.21 | |
| 6932.63 | |
| 7704.19 | L29 |
| 9125.31 | |

Table S8 shows the ribosomal protein mass peaks of *B. subtilis* ProcB6c. Specifically, ribosomal protein L36, L33 and L29 could be mapped to mass peaks 4302.22 Da, 5895.12 Da, and 7704.19 Da, respectively.

| **Table S9:** ***Bacillus subtilis*** **Peak list of ATCC 6633** | **Ribosomal Protein** |
|---|---|
| 2176.21 | |
| 2746.54 | |
| 3322.05 | |
| 3340.07 | |
| 3359.74 | |
| 3421.85 | |
| 3867.27 | |
| 3888.64 | |
| 4019.1 | |
| 4304.53 | L36 |
| 4719.08 | |
| 4942.21 | |
| 5002.02 | |
| 5223.14 | |
| 5897.24 | L33 |
| 6504.36 | |
| 6594.67 | |
| 6675.49 | |
| 7725.36 | L29 |
| 9430.65 | |
| 9879.24 | |

Table S9 shows the ribosomal protein mass peaks of *B. subtilis* ATCC 6633. Specifically, ribosomal protein L36, L33 and L29 could be mapped to mass peaks 4304.53 Da, 5897.24 Da, and 7725.36 Da, respectively.

| **Table S10: *Carnobacterium maltaromaticum* Peak list of ATCC 27865** | **Ribosomal protein** |
|---|---|
| 2172.97 | |
| 2901.46 | |
| 3237.71 | |
| 3349.16 | |
| 3435.99 | |
| 4101.31 | |
| 4345.93 | L36 |
| 5659.39 | |
| 5803.91 | |
| 6209.95 | |
| 6347.27 | |
| 6474.56 | L30 |
| 6697.89 | |
| 6871.44 | |
| 6927.45 | |
| 7441.11 | L29 |
| 9083.43 | |

Table S10 shows the annotated ribosomal protein mass peaks of *Carnobacterium maltaromaticum* ATCC 27865. Specifically, ribosomal protein L36, L30, and L29 mapped to mass peaks 4345.93 Da, 6474.56 Da, and 7441.11 Da, respectively.

| **Table S11:** ***Carnobacterium maltaromaticum*** **Peak list of ATCC 35586** | **Ribosomal protein** |
|---|---|
| 2173.74 | |
| 2902.14 | |
| 3239.13 | |
| 3350.58 | |
| 3436.62 | |
| 4114.74 | |
| 4347.5 | L36 |
| 4784 | |
| 5805.47 | |
| 6210.88 | |
| 6347.05 | |
| 6476.6 | L30 |
| 6699.68 | |
| 6873.19 | |
| 7534.46 | |

Table S11 shows the annotated ribosomal protein mass peaks of *C. maltaromaticum* ATCC 35586. Specifically, ribosomal protein L36 and L30 mapped to mass peaks 4347.5 Da, and 6476.6 Da, respectively.

| **Table S12:** ***Carnobacterium maltaromaticum*** **Peak list of Proc3T4** | **Ribosomal protein** |
|---|---|
| 2174.5 | |
| 2902.83 | |
| 3238.89 | |
| 3350.47 | |
| 3436.94 | |
| 4102.52 | |
| 4347.76 | L36 |
| 4783.76 | |
| 5803.21 | |
| 6347.3 | |
| 6475.08 | L30 |
| 6698.47 | |
| 6871.2 | |

Table S12 shows the annotated ribosomal protein mass peaks of *C. maltaromaticum* Proc3T4. Specifically, ribosomal protein L36 and L30 mapped to mass peaks 4347.76 Da, and 6475.08 Da, respectively.

| **Table S13:** ***Carnobacterium maltaromaticum*** **Peak list of Proc4T4** | **Ribosomal protein** |
|---|---|
| 2173.64 | |
| 2902.03 | |
| 3173.96 | |
| 3237.7 | |
| 3349.37 | |
| 3435.8 | |
| 3721.31 | |
| 4101.19 | |
| 4346.09 | L36 |
| 5803.12 | |
| 6346.54 | |
| 6474.53 | L30 |
| 6870.38 | |

Table S13 shows the annotated ribosomal protein mass peaks of *C. maltaromaticum* Proc4T4. Specifically, ribosomal protein L36 and L30 mapped to mass peaks 4346.09 Da and 6474.53 Da, respectively.

| **Table S14:** ***Bacillus thuringiensis*** **Peak list of ATCC 10792** | **Ribosomal protein** |
|---|---|
| 2167.76 | |
| 2240.53 | |
| 2277.64 | |
| 2586.25 | |
| 2788.8 | |
| 2827.62 | |
| 3089.6 | |
| 3116.68 | |
| 3650.28 | |
| 3706.38 | |
| 3744.36 | |
| 4331.48 | L36 |
| 4548.42 | |
| 4992.08 | |
| 5474.69 | |

Table S14 shows the annotated ribosomal protein mass peak of *Bacillus thuringiensis* ATCC 10792. Specifically, ribosomal protein L36 could be mapped to mass peak 4331.48 Da.

| **Table S15:** ***Bacillus thuringiensis*** **Peak list of ATCC 33679** | **Ribosomal protein** |
|---|---|
| 2167.44 | |
| 2532.95 | |
| 2661.97 | |
| 2809.3 | |
| 3088.89 | |
| 3116.82 | |
| 3561.55 | |
| 3651.46 | |
| 3707.88 | |
| 3745.31 | |
| 4030.13 | |
| 4333.35 | L36 |
| 4551.44 | |
| 4992.08 | |
| 5052.78 | |
| 5291.61 | |
| 5473.64 | |
| 6714.85 | |

Table S15 shows the annotated ribosomal protein mass peak of *B. thuringiensis* ATCC 33679. Specifically, ribosomal protein L36 could be mapped to mass peak 4333.35 Da.

| **Table S16:** ***Bacillus thuringiensis*** **Peak list of ATCC 35866** | **Ribosomal protein** |
|---|---|
| 2169.68 | |
| 2535.49 | |
| 2664.71 | |
| 2812.35 | |
| 3091.31 | |
| 3119.45 | |
| 3563.62 | |
| 3653.49 | |
| 3709.91 | |
| 3747.7 | |
| 4335.52 | L36 |
| 4552.8 | |
| 4994.56 | |

| |
|---|
| 5293.88 |
| 5474.72 |
| 6714.01 |

Table S16 shows the annotated ribosomal protein mass peak of *B. thuringiensis* ATCC 35866. Specifically, ribosomal protein L36 could be mapped to mass peak 4335.52 Da.

| **Table S17: *Escherichia coli* Peak list of NCTC 50271** | **Ribosomal protein** |
|---|---|
| 2183.9 | |
| 2549.6 | |
| 2691.85 | |
| 2835.87 | |
| 3128.84 | |
| 3159.23 | |
| 3580.14 | |
| 3638.35 | |
| 3674.36 | |
| 3935.81 | |
| 4185.91 | |
| 4365.37 | L36 |
| 4769.21 | |
| 4777.77 | |
| 4871.61 | |
| 5097 | |
| 5151.29 | |
| 5381.55 | L34 |
| 5613.38 | |
| 6255.75 | |
| 6316.69 | |
| 6411.59 | |
| 7157.99 | |
| 7274.48 | L29 |
| 7868.46 | L31 |
| 8328.41 | |
| 8370.72 | |
| 8433.22 | |
| 8881.04 | |
| 8997.4 | S18 |
| 9066.28 | |
| 9226.12 | |
| 9543.35 | |

| |
|---|
| 9742.91 |

Table S17 shows the annotated ribosomal protein mass peaks of *Escherichia coli* NCTC 50271. Specifically, ribosomal protein L36, L34, L29, L31 and S18 mapped to mass peaks 4365.37 Da, 5381.55 Da, 7274.48 Da, 7868.46 Da, and 8997.4 Da, respectively.

| **Table S18: *Escherichia coli* Peak list of NCTC 50365** | **Ribosomal protein** |
|---|---|
| 2183.62 | |
| 2691.42 | |
| 2835.51 | |
| 3128.33 | |
| 3158.63 | |
| 3206.34 | |
| 3579.8 | |
| 3637.69 | |
| 3674.05 | |
| 3936.26 | |
| 4185.37 | |
| 4364.77 | L36 |
| 4438.9 | |
| 4497.54 | |
| 4768.43 | |
| 4777.23 | |
| 5096.3 | |
| 5150.45 | |
| 5380.66 | L34 |
| 6254.58 | |
| 6315.62 | |
| 6410.82 | |
| 6507.21 | |
| 7157.63 | |
| 7273.59 | L29 |
| 7870.4 | L31 |
| 8119 | |
| 8369.32 | |
| 8992.76 | S18 |
| 9225.82 | |
| 9543.13 | |

Table S18 shows the annotated ribosomal protein mass peaks of *E. coli* NCTC 50365. Specifically, ribosomal protein L36, L34, L29, L31 and S18 mapped to mass peaks 4364.77 Da, 5380.66 Da, 7273.59 Da, 7870.4 Da, and 8992.76 Da, respectively.

| **Table S19:** ***Proteus vulgaris*** **Peak list of ATCC 9484** | **Ribosomal protein** |
|---|---|
| 2243.02 | |
| 2756.12 | |
| 2826.13 | |
| 3137.47 | |
| 3554.64 | |
| 3637.69 | |
| 3908.88 | |
| 3982.9 | |
| 4186.21 | |
| 4484.75 | L36 |
| 4737.77 | L36 |
| 4770.95 | |
| 4796.96 | |
| 5130.52 | |
| 5510.96 | L34 |
| 6271.2 | |
| 6504.92 | |
| 7107.96 | |
| 7273.86 | L29 |
| 7816.65 | L31 |
| 7965.29 | |
| 8370.57 | |
| 9476.56 | |

Table S19 shows the annotated ribosomal protein mass peak of *Proteus vulgaris* ATCC 9484. Specifically, ribosomal protein L36, L36, L34, L29 and L31 mapped to mass peaks 4484.75 Da, 4737.77 Da, 5510.96 Da, 7273.86 Da, and 7816.65 Da, respectively.

| **Table S20:** ***Proteus vulgaris*** **Peak list of Sard1** | **Ribosomal protein** |
|---|---|
| 2242.85 | |
| 2748.12 | |
| 2825.39 | |
| 3138.03 | |
| 3252.09 | |
| 3554.41 | |
| 3637.85 | |
| 3907.44 | |
| 3954.56 | |
| 4185.05 | |
| 4484.54 | L36 |
| 4738.26 | L36 |
| 4770.76 | |
| 4804.11 | |
| 5130.76 | |
| 5395.04 | |
| 5496.02 | |
| 6274.28 | |
| 7108.03 | |
| 7274.2 | L29 |
| 7908.74 | |
| 9480.71 | |

Table S20 shows the annotated ribosomal protein mass peaks of *P. vulgaris* Sard1. Specifically, ribosomal protein L36, L36 and L29 mapped to mass peaks 4484.54 Da, 4738.26 Da and 7274.2 Da, respectively.

| **Table S21:** ***Proteus vulgaris*** **Peak list of Sard2** | **Ribosomal protein** |
|---|---|
| 2242.79 | |
| 2748.23 | |
| 2825.82 | |
| 3137.89 | |
| 3554.49 | |
| 3637.52 | |
| 3954.24 | |
| 4185.52 | |
| 4484.6 | L36 |
| 4737.85 | L36 |
| 4768.4 | |
| 4802.54 | |
| 5130.18 | |
| 5495.92 | |
| 6274.87 | |
| 6504.05 | |
| 7107.18 | |
| 7273.65 | L29 |
| 7908.69 | |
| 9476.8 | |

Table S21 shows the annotated ribosomal protein mass peaks of *P. vulgaris* Sard2. Specifically, ribosomal protein L36, L36 and L29 mapped to mass peaks 4484.6 Da, 4737.85 Da, and 7273.65 Da, respectively.

| **Table S22:** ***Proteus vulgaris*** **Peak list of Sard3** | **Ribosomal protein** |
|---|---|
| 2242.1 | |
| 2747.53 | |
| 2755.45 | |
| 2825.17 | |
| 3137.33 | |
| 3145.7 | |
| 3554.27 | |
| 3636.79 | |
| 3644.07 | |
| 3953.37 | |
| 4184.02 | |
| 4483.36 | L36 |
| 4737.11 | L36 |
| 4769.38 | |
| 4804.59 | |
| 5018.14 | |
| 5130.49 | |
| 5495.18 | |
| 6274.46 | |
| 6501.19 | |
| 7272.4 | L29 |
| 7906.89 | |
| 9475.6 | |

Table S22 shows the annotated ribosomal protein mass peaks of *P. vulgaris* Sard3. Specifically, ribosomal protein L36, L36 and L29 mapped to mass peaks 4483.36 Da, 4737.11 Da, and 7272.4 Da, respectively.

| **Table S23:** ***Proteus vulgaris*** **Peak list of Sard4** | **Ribosomal protein** |
|---|---|
| 2242.79 | |
| 2747.9 | |
| 2825.64 | |
| 3137.75 | |
| 3554.23 | |
| 3637.41 | |
| 3954.11 | |
| 4185.36 | |
| 4484.38 | L36 |
| 4737.76 | L36 |
| 4769.85 | |
| 4803.13 | |
| 5130.75 | |
| 5393.68 | |
| 5495.39 | |
| 6104.41 | |
| 6273.98 | |
| 6503.08 | |
| 7107.91 | |
| 7273.73 | L29 |
| 7817.02 | L31 |
| 7908.06 | |
| 9475.01 | |
| 9542.6 | |

Table S23 shows the annotated ribosomal protein mass peaks of *P. vulgaris* Sard4. Specifically, ribosomal protein L36, L36, L29 and L31 mapped to mass peaks 4484.38 Da, 4737.76 Da, 7273.73 Da, and 7817.02 Da, respectively.

| **Table S24:** ***Pseudomonas fluorescens*** **Peak list of ATCC 13525** | **Ribosomal protein** |
|---|---|
| 2218.13 | |
| 2534.37 | |
| 3041.18 | |
| 3313.31 | |
| 3410.12 | |
| 3439.65 | |
| 3585.49 | |
| 3618.21 | |
| 4127.14 | |
| 4172.41 | |
| 4432.9 | |
| 4978.92 | |
| 5065.27 | L34 |
| 5671.68 | |
| 5802.48 | |
| 6078.9 | |
| 6264.15 | |
| 6391.72 | |
| 6621.13 | |
| 7169.58 | L29 |
| 7593.99 | |
| 8250.18 | |
| 8341.15 | |
| 8817.33 | |
| 9059.07 | |
| 9549.66 | |
| 9809.05 | |
| 9950.38 | |

Table S24 shows the annotated ribosomal protein mass peaks of *Pseudomonas fluorescens* ATCC 13525. Specifically, ribosomal protein L34 and L29 mapped to mass peaks 5065.27 Da and 7169.58 Da, respectively.

| **Table S25: *Pseudomonas fluorescens* Peak list of ATCC 17397** | **Ribosomal protein** |
|---|---|
| 2218.46 | |
| 2535.19 | |
| 3042.39 | |
| 3092.5 | |
| 3320.97 | |
| 3421.51 | |
| 3587.57 | |
| 4129.03 | |
| 4434.32 | |
| 4776.92 | |
| 4905.49 | |
| 4978.51 | |
| 5067.16 | L34 |
| 5662.46 | |
| 6081.58 | |
| 6182.15 | |
| 6394.48 | |
| 6638.75 | |
| 7173.86 | L29 |
| 8253.78 | |
| 8389.35 | S21 |
| 8507.78 | |
| 9547.99 | |
| 9802.82 | |

Table S25 shows the annotated ribosomal protein mass peaks of *P. fluorescens* ATCC 17397. Specifically, ribosomal protein L34, L29 and S21 mapped to mass peaks 5067.16 Da, 7173.86 Da, and 8389.35 Da, respectively.

| **Table S26:** ***Pseudomonas fluorescens*** **Peak list of Turb28** | **Ribosomal protein** |
|---|---|
| 2146.68 | |
| 2217.55 | |
| 2264.03 | |
| 2533.88 | |
| 2615.63 | |
| 2863.63 | |
| 3040.75 | |
| 3197.45 | |
| 3279.44 | |
| 3305.38 | |
| 3406.58 | |
| 3586.31 | |
| 3618.18 | |
| 3769.28 | |
| 4127.6 | |
| 4164.35 | |
| 4433.1 | |
| 4531.64 | |
| 4910.69 | |
| 4980.32 | |
| 5023.32 | |
| 5066.2 | L34 |
| 5101.41 | |
| 5570.12 | |
| 5649.85 | |
| 6080.2 | |
| 6393.94 | |
| 6610.19 | |
| 7171.94 | L29 |
| 7234.88 | |
| 8328.98 | |
| 8403.79 | |
| 9061.55 | |
| 9820.9 | |

Table S26 shows the annotated ribosomal protein mass peaks of *P. fluorescens* Turb28. Specifically, ribosomal protein L34 and L29 mapped to mass peaks 5066.2 Da, and 7171.94 Da, respectively.

| **Table S27:** ***Pseudomonas fluorescens*** **Peak list of Turb46** | **Ribosomal protein** |
|---|---|
| 2217.39 | |
| 2533.6 | |
| 3040.61 | |
| 3196.99 | |
| 3305.29 | |
| 3406.03 | |
| 3586.04 | |
| 3768.33 | |
| 4128.32 | |
| 4164.29 | |
| 4201.95 | |
| 4432.73 | |
| 4531.45 | |
| 4910.58 | |
| 4979.35 | |
| 5024.03 | |
| 5066.18 | L34 |
| 5571.02 | |
| 5649.83 | |
| 6080.24 | |
| 6392.7 | |
| 6611.17 | |
| 7171.35 | L29 |
| 8330.48 | |
| 8404.81 | |
| 9063.62 | |
| 9820.2 | |

Table S27 shows the annotated ribosomal protein mass peaks of *P. fluorescens* Turb46. Specifically, ribosomal protein L34 and L29 mapped to mass peaks 5066.18 Da and 7171.35 Da, respectively.

| **Table S28:** ***Pseudomonas fluorescens*** **Peak list of Turb52** | **Ribosomal protein** |
|---|---|
| 2217.67 | |
| 2534.1 | |
| 3040.61 | |
| 3197.43 | |
| 3306.02 | |
| 3406.53 | |
| 3586.59 | |
| 4128.24 | |
| 4165 | |
| 4433.73 | |
| 4980.5 | |
| 5066.93 | L34 |
| 5570.51 | |
| 5650.1 | |
| 6079.88 | |
| 6393.33 | |
| 6610.97 | |
| 7172.54 | L29 |
| 8329.14 | |
| 8404.17 | |

Table S28 shows the annotated ribosomal protein mass peaks of *P. fluorescens* Turb52. Specifically, ribosomal protein L34 and L29 mapped to mass peaks 5066.93 Da and 7172.54 Da, respectively.

| **Table S29:** ***Pseudomonas fluorescens*** **Peak list of Turb64** | **Ribosomal protein** |
|---|---|
| 2217.64 | |
| 2534.05 | |
| 3041.11 | |
| 3197.58 | |
| 3306.34 | |
| 3406.16 | |
| 3586.53 | |
| 4128.41 | |
| 4164.71 | |
| 4433.81 | |
| 4531.23 | |
| 4981.46 | |
| 5025.13 | |
| 5066.99 | L34 |
| 5570.9 | |
| 5650.72 | |
| 6081.48 | |
| 6393.62 | |
| 6611.33 | |
| 7172.21 | L29 |
| 8330.87 | |
| 8403.79 | |

Table S29 shows the annotated ribosomal protein mass peaks of *P. fluorescens* Turb64. Specifically, ribosomal protein L34 and L29 mapped to mass peaks 5066.99 Da and 7172.21 Da, respectively.

| **Table S30: *Pseudomonas fragi* Peak list of ATCC 4973** | **Ribosomal protein** |
|---|---|
| 2218.25 | |
| 2534.5 | |
| 3023.51 | |
| 3306.55 | |
| 3594.14 | |
| 3619.02 | |
| 3802.55 | |
| 4128.54 | |
| 4434.25 | L36 |
| 4466.35 | |
| 4951.58 | |
| 5067.2 | L34 |
| 5660.24 | |
| 6045.46 | L33 |
| 6379.59 | |
| 6611.34 | |
| 7186.2 | L29 |
| 7236.31 | |
| 7603.59 | |
| 8258.66 | |
| 8343.2 | |
| 8932.11 | S18 or L28 |
| 9130.84 | |
| 9560.8 | |
| 9896.74 | |

Table S30 shows the annotated ribosomal protein mass peaks of *Pseudomonas fragi* ATCC 4973. Specifically, ribosomal protein L36, L34, L33 and L29 mapped to mass peaks 4434.25 Da, 5067.2 Da, 6045.46 Da, and 7186.2 Da, respectively. Finally, ribosomal protein S18 or L28 could be mapped to mass peak 8932.11 Da.

| **Table S31:** ***Pseudomonas fragi*** **Peak list of Seab03** | **Ribosomal protein** |
|---|---|
| 2217.66 | |
| 2534.14 | |
| 3022.73 | |
| 3195.58 | |
| 3305.54 | |
| 3592.59 | |
| 3793.56 | |
| 4126.58 | |
| 4432.32 | L36 |
| 4945.07 | |
| 5064.76 | L34 |
| 5674.63 | |
| 6006.17 | |
| 6041.56 | L33 |
| 6387.67 | |
| 6607.58 | |
| 7180.51 | L29 |
| 7478.33 | |
| 7584.42 | |
| 8248.91 | |
| 8898.8 | |
| 9887.49 | |

Table S31 shows the annotated ribosomal protein mass peaks of *P. fragi* Seab03. Specifically, ribosomal protein L36, L34, L33 and L29 mapped to mass peaks 4432.32 Da, 5064.76 Da, 6041.56 Da, and 7180.51 Da, respectively.

| **Table S32:** ***Pseudomonas fragi*** **Peak list of Seab22** | **Ribosomal protein** |
|---|---|
| 2217.55 | |
| 2533.73 | |
| 3022.57 | |
| 3195.99 | |
| 3304.63 | |
| 3592.68 | |
| 4126.73 | |
| 4432.22 | L36 |
| 4559.82 | |
| 4937.92 | |
| 5065.07 | L34 |
| 5347.24 | |
| 5676.11 | |
| 6042.2 | L33 |
| 6392.44 | |
| 6607.74 | |
| 7186.26 | L29 |
| 8251.55 | |
| 8374.17 | S21 |
| 9124.91 | |
| 9873.79 | |

Table S32 shows the annotated ribosomal protein mass peaks of *P. fragi* Seab22. Specifically, ribosomal protein L36, L34, L33, L29 and S21 mapped to mass peaks 4432.22 Da, 5065.07 Da, 6042.2 Da, 7186.26 Da, and 8374.17 Da, respectively.

| **Table S33:** ***Pseudomonas fragi*** **Peak list of Seab23** | **Ribosomal protein** |
|---|---|
| 2217.71 | |
| 2533.96 | |
| 2832.65 | |
| 3023.02 | |
| 3306.05 | |
| 3597.88 | |
| 3617.4 | |
| 3801.01 | |
| 4126.71 | |
| 4432.58 | L36 |
| 4943.55 | |
| 5065.44 | L34 |
| 5662.64 | |
| 6042.89 | L33 |
| 6376.04 | |
| 6608.14 | |
| 7192.64 | L29 |
| 7448.67 | |
| 7599.18 | |
| 8250.47 | |
| 9015.19 | |
| 9889.83 | |

Table S33 shows the annotated ribosomal protein mass peaks of *P. fragi* Seab23. Specifically, ribosomal protein L36, L34, L33 and L29 mapped to mass peaks 4432.58 Da, 5065.44 Da, 6042.89 Da, and 7192.64 Da, respectively.

| **Table S34:** ***Pseudomonas fragi*** **Peak list of Turb32** | **Ribosomal protein** |
|---|---|
| 2218.02 | |
| 2396.24 | |
| 2534.34 | |
| 2840.56 | |
| 3023.3 | |
| 3306.42 | |
| 3593.4 | |
| 3741.36 | |
| 4128.35 | |
| 4434.69 | L36 |
| 4949.76 | |
| 5024.89 | |
| 5067.56 | L34 |
| 5679.6 | |
| 6008.8 | |
| 6045.86 | L33 |
| 6394.05 | |
| 6611.98 | |
| 7185.56 | L29 |
| 7482.47 | |
| 7588.81 | |
| 7743.63 | |
| 8257.25 | |
| 8906.24 | |

Table S34 shows the annotated ribosomal protein mass peaks of *P. fragi* Turb32. Specifically, ribosomal protein L36, L34, L33 and L29 mapped to mass peaks 4434.69 Da, 5067.56 Da, 6045.86 Da and 7185.56 Da, respectively.

| **Table S35:** ***Pseudomonas fragi*** **Peak list of Turb43** | **Ribosomal protein** |
|---|---|
| 2217.79 | |
| 2534.22 | |
| 2675.91 | |
| 3023.3 | |
| 3198.17 | |
| 3306.4 | |
| 3419.82 | |
| 3594.07 | |
| 3618.78 | |
| 4128.11 | |
| 4187.58 | |
| 4402.08 | |
| 4434.3 | L36 |
| 4569.68 | |
| 4941.35 | |
| 5025.02 | |
| 5067.4 | L34 |
| 5350.84 | |
| 5679.14 | |
| 6046.1 | L33 |
| 6396.74 | |
| 6611.42 | |
| 7188.59 | L29 |
| 8257.04 | |
| 9135.63 | |

Table S35 shows the annotated ribosomal protein mass peaks of *P. fragi* Turb43. Specifically, ribosomal protein L36, L34, L33 and L29 mapped to mass peaks 4434.3 Da, 5067.4 Da, 6046.1 Da, and 7188.59 Da, respectively.

| **Table S36:** ***Pseudomonas fragi*** **Peak list of Turb47** | **Ribosomal protein** |
|---|---|
| 2217.82 | |
| 2534.19 | |
| 2840.26 | |
| 3004.94 | |
| 3023.19 | |
| 3306.1 | |
| 3566.2 | |
| 3592.08 | |
| 3741.1 | |
| 3794.54 | |
| 3871.25 | |
| 4127.74 | |
| 4400.9 | |
| 4434.02 | L36 |
| 4452.05 | |
| 4950.27 | |
| 5066.89 | L34 |
| 5524.12 | |
| 5678.83 | |
| 6008.74 | |
| 6044.13 | L33 |
| 6610.9 | |
| 7185.28 | L29 |
| 7481.68 | |
| 7588.16 | |
| 7741.7 | |
| 8905.79 | |

Table S36 shows the annotated ribosomal protein mass peaks of *P. fragi* Turb47. Specifically, ribosomal protein L36, L34, L33 and L29 mapped to mass peaks 4434.02 Da, 5066.89 Da, 6044.13 Da, and 7185.28 Da, respectively.

| **Table S37:** ***Pseudomonas putida*** **Peak list of ATCC 12633** | **Ribosomal protein** |
|---|---|
| 2217.99 | |
| 2369.8 | |
| 2570.18 | |
| 3334.4 | |
| 3587.68 | |
| 3811.35 | |
| 4082.54 | |
| 4121.92 | |
| 4337.7 | |
| 4435.01 | L36 |
| 4738.73 | |
| 5138.92 | L34 |
| 5355.65 | |
| 5631.79 | |
| 5991.17 | L33 |
| 6134.23 | |
| 6333.91 | |
| 6665.06 | |
| 7173.76 | |
| 7450.7 | |
| 7620.85 | |
| 8081.53 | |
| 8166.51 | |
| 8242.79 | |
| 8672.35 | |
| 9132.55 | |
| 9232.21 | |
| 9545.16 | |

Table S37 shows the annotated ribosomal protein mass peaks of *Pseudomonas putida* ATCC 12633. Specifically, ribosomal protein L36, L34 and L33 mapped to mass peaks 4435.01 Da, 5138.92 Da, and 5991.17 Da, respectively.

| **Table S38:** ***Pseudomonas putida*** **Peak list of ATCC 17453** | **Ribosomal protein** |
|---|---|
| 2569.61 | |
| 3009.75 | |
| 3023.33 | |
| 3586.3 | |
| 4119.03 | |
| 4126.38 | |
| 4434.13 | L36 |
| 5137.76 | L34 |
| 6018.8 | |
| 6045.35 | |
| 6320.07 | |
| 6637.74 | |
| 7173.52 | |
| 7395.33 | |
| 7598.87 | |
| 7811.86 | |
| 8060.09 | |
| 8164.4 | |
| 8237.58 | |
| 8731.56 | |
| 9541.24 | |

Table S38 shows the annotated ribosomal protein mass peaks of *P. putida* ATCC 17453. Specifically, ribosomal protein L36 and L34 mapped to mass peaks 4434.13 Da, and 5137.76 Da, respectively.

| **Table S39:** ***Pseudomonas putida*** **Peak list of Seab04** | **Ribosomal protein** |
|---|---|
| 2890.61 | |
| 3070.45 | |
| 4432.42 | L36 |
| 4738.43 | |
| 5136.12 | L34 |
| 5628.34 | |
| 5777.67 | |
| 5988.14 | L33 |
| 6137.9 | |
| 6317.58 | |
| 6606.99 | |
| 7166.21 | |
| 7461.82 | |
| 7599.67 | |
| 8054.28 | |
| 8146.73 | |
| 8228.4 | |
| 8724.94 | |

Table S39 shows the annotated ribosomal protein mass peaks of *P. putida* Seab04. Specifically, ribosomal protein L36, L34 and L33 mapped to mass peaks 4432.42 Da, 5136.12 Da, and 5988.14 Da, respectively.

| **Table S40: *Pseudomonas putida* Peak list of Seab11** | **Ribosomal protein** |
|---|---|
| 2937.11 | |
| 3023.63 | |
| 4120.04 | |
| 4432.63 | L36 |
| 4737.1 | |
| 5136.19 | L34 |
| 5629.18 | |
| 5872.29 | |
| 5988.12 | L33 |
| 6044.71 | |
| 6662.18 | |
| 7169.29 | |
| 7270.03 | |
| 7331.24 | |
| 7417.99 | |
| 7636.59 | |
| 8078.17 | |
| 8238.29 | |
| 9536.36 | |

Table S40 shows the annotated ribosomal protein mass peaks of *P. putida* Seab11. Specifically, ribosomal protein L36, L34 and L33 mapped to mass peaks 4432.63 Da, 5136.19 Da, and 5988.12 Da, respectively.

| **Table S41: *Pseudomonas syringae* Peak list of ATCC 19304** | **Ribosomal protein** |
|---|---|
| 2218.98 | |
| 2563.6 | |
| 2988.21 | |
| 3329.3 | |
| 3586.92 | |
| 3618.49 | |
| 3781.27 | |
| 4127.81 | |
| 4434.01 | L36 |
| 4835.21 | |
| 4910.66 | |
| 4937.76 | |
| 5123.34 | L34 |
| 5645.29 | |
| 5671.26 | |
| 5972.74 | L33 |
| 6340.97 | |
| 6419.84 | |
| 6651.03 | |
| 7050.78 | |
| 7169.92 | L29 |
| 7234.24 | |
| 7441.24 | |
| 7558.46 | |
| 7608.87 | |
| 7877.56 | |
| 8251.49 | |
| 8429.51 | |
| 8797.11 | |
| 9104.68 | |
| 9663.31 | |
| 9874.55 | |

Table S41 shows the annotated ribosomal protein mass peaks of *Pseudomonas syringae* ATCC 19304. Specifically, ribosomal protein L36, L34, L33, and L29 mapped to mass peaks 4434.01 Da, 5123.34 Da, 5972.74 Da, and 7169.92 Da, respectively.

| **Table S42:** ***Pseudomonas syringae*** **Peak list of ATCC 19310** | **Ribosomal protein** |
|---|---|
| 2218.81 | |
| 2563.62 | |
| 2988.39 | |
| 3329.73 | |
| 3587.44 | |
| 3619.73 | |
| 3635.03 | |
| 3790.01 | |
| 4128.61 | |
| 4434.79 | L36 |
| 4553.87 | |
| 4836.74 | |
| 4899.87 | |
| 4938.4 | |
| 5124.62 | L34 |
| 5655.32 | |
| 5670.58 | |
| 5806.93 | |
| 5974.41 | L33 |
| 6356.53 | |
| 6421.91 | |
| 6653.71 | |
| 7053.86 | |
| 7172.86 | L29 |
| 7577.82 | |
| 8070.77 | |
| 8254.53 | |
| 9104.3 | |
| 9670.45 | |
| 9797.76 | |
| 9875.56 | |

Table S42 shows the annotated ribosomal protein mass peaks of *P. syringae* ATCC 19310. Specifically, ribosomal protein L36, L34, L33 and L29 mapped to mass peaks 4434.79 Da, 5124.62 Da, 5974.41 Da and 7172.86 Da, respectively.

| ***Table S43: Pseudomonas syringae*** **Peak list of Seab02** | **Ribosomal protein** |
|---|---|
| 2217.71 | |
| 2563.36 | |
| 3029.91 | |
| 3283.04 | |
| 3305.42 | |
| 3587.12 | |
| 3784.88 | |
| 3803.28 | |
| 3971.19 | |
| 4071.21 | |
| 4128.84 | |
| 4433.9 | L36 |
| 4559.07 | |
| 4824.11 | |
| 4883.48 | |
| 5124.01 | L34 |
| 5542.99 | |
| 5611.99 | |
| 5677.86 | |
| 5924.31 | |
| 5986.91 | L33 |
| 6059.67 | |
| 6320.54 | |
| 6610.79 | |
| 7173.39 | L29 |
| 7809.89 | |
| 7941.5 | |
| 8142.46 | L31 |
| 8208.77 | |
| 9116.83 | |
| 9646.28 | |
| 9761.81 | |

Table S43 shows the annotated ribosomal protein mass peaks of *P. syringae* Seab02. Specifically, ribosomal protein L36, L34, L33, L29 and L31 mapped to mass peaks 4433.9 Da, 5124.01, 5986.91 Da, 7173.39 Da, and 8142.46 Da, respectively.

| **Table S44:** ***Serratia marcescens*** **Peak list of ATCC 274** | **Ribosomal protein** |
|---|---|
| 2690.88 | |
| 2825.81 | |
| 3961.64 | |
| 4184.37 | |
| 4348.98 | |
| 4605.56 | |
| 4767.21 | |
| 4780.4 | |
| 5222.31 | |
| 5337.59 | |
| 5379.88 | |
| 6115.85 | |
| 6225.69 | |
| 6441.74 | |
| 6544.85 | |
| 7207.43 | |
| 7271.83 | L29 |
| 7923.18 | |
| 8367.62 | |
| 9209.87 | |
| 9537.01 | |
| 9930.37 | |

Table S44 shows the annotated ribosomal protein mass peak of *Serratia marcescens* ATCC 274. Specifically, ribosomal protein L29 mapped to mass peak 7271.83 Da.

| **Table S45:** ***Serratia marcescens*** **Peak list of Proc7T6** | **Ribosomal protein** |
|---|---|
| 2175.42 | |
| 2691.11 | |
| 2825.41 | |
| 3963 | |
| 4185.97 | |
| 4348.88 | |
| 4606.44 | |
| 4769.38 | |
| 4983.26 | |
| 5380.37 | |
| 6115.6 | |
| 6226.09 | |
| 7925.55 | |
| 9537.52 | |

Table S45 revealed that no ribosomal protein mass peak could be annotated for *Serratia marcescens* Proc7T6.

| **Table S46:** ***Serratia proteamaculans*** **Peak list of Proc5T6** | **Ribosomal protein** |
|---|---|
| 2175.37 | |
| 2697.53 | |
| 2825.68 | |
| 3009.08 | |
| 3947.97 | |
| 4184.58 | |
| 4349.02 | |
| 4782.23 | |
| 5396.01 | L34 |
| 6014.73 | |
| 6238.93 | |
| 6413 | |
| 7299.35 | L29 |
| 7894.7 | |
| 9564.1 | |

Table S46 shows the annotated ribosomal protein mass peaks of *Serratia proteamaculans* Proc5T6. Specifically, ribosomal protein L34 and L29 mapped to mass peaks 5396.01 Da and 7299.35 Da, respectively.

| **Table S47:** ***Serratia proteamaculans*** **Peak list of ProcB1** | **Ribosomal protein** |
|---|---|
| 2175.72 | |
| 2698.58 | |
| 2825.49 | |
| 3120.09 | |
| 3948.5 | |
| 4182.37 | |
| 4346.52 | |
| 4489.35 | |
| 4639.16 | |
| 4780.29 | |
| 4913.8 | |
| 5392.3 | L34 |
| 6055.52 | |
| 6234.91 | |
| 6408.48 | |
| 7283.1 | |
| 7891.92 | |
| 9553.46 | |

Table S47 shows the ribosomal protein mass peaks of *S. proteamaculans* ProcB1. Specifically, ribosomal protein L34 mapped to mass peak 5392.3 Da.

| **Table S48:** ***Serratia proteamaculans*** **Peak list of ProcB4** | **Ribosomal protein** |
|---|---|
| 2175.45 | |
| 2698.13 | |
| 2824.88 | |
| 3029.82 | |
| 3119.55 | |
| 3947.88 | |
| 4181.74 | |
| 4345.76 | |
| 4638.32 | |
| 4779.56 | |
| 4914.07 | |
| 5391.35 | L34 |
| 6054.42 | |
| 6233.85 | |
| 6407.22 | |
| 7283.28 | |
| 7890.62 | |
| 9552.23 | |

Table S48 shows the annotated ribosomal protein mass peak of *S. proteamaculans* ProcB4. Specifically, ribosomal protein L34 mapped to mass peak 5391.35 Da.

| **Table S49:** ***Staphylococcus aureus*** **Peak list of ATCC 9144** | **Ribosomal protein** |
|---|---|
| 2752.79 | |
| 2868.66 | |
| 3275.22 | |
| 3443.2 | |
| 4303.52 | L36 |
| 5029.81 | |
| 5416.9 | |
| 5504.14 | |
| 6549.01 | L30 |
| 6885.29 | |

Table S49 shows the annotated ribosomal protein mass peaks of *Staphylococcus aureus* ATCC 9144. Specifically, ribosomal protein L36 and L30 mapped to mass peaks 4303.52 Da and 6549.01 Da, respectively.

| **Table S50:** ***Staphylococcus aureus*** **Peak list of ATCC 35845** | **Ribosomal protein** |
|---|---|
| 2753.49 | |
| 2869.49 | |
| 3275.95 | |
| 3444.1 | |
| 4304.48 | L36 |
| 5031.13 | |
| 5505.55 | |
| 6550.61 | L30 |
| 6886.98 | |

Table S50 shows the annotated ribosomal protein mass peaks of *S. aureus* ATCC 35845. Specifically, ribosomal protein L36 and L30 mapped to mass peaks 4304.48 Da and 6550.61 Da, respectively.

| **Table S51:** ***Staphylococcus aureus*** **Peak list of Proc1010** | **Ribosomal protein** |
|---|---|
| 2278.68 | |
| 2288.72 | |
| 2306.88 | |
| 2607.93 | |
| 2636.06 | |
| 2674.16 | |
| 2979.17 | |
| 3007.3 | |
| 3444.47 | |
| 4303.99 | L36 |
| 5031.55 | |
| 5524.05 | |
| 6887.43 | |

Table S51 shows the annotated ribosomal protein mass peak of *S. aureus* Proc1010. Specifically, ribosomal protein L36 mapped to mass peak 4303.99 Da.

| **Table S52:** ***Stenotrophomonas maltophilia*** **Peak list of 5PC6** | **Ribosomal protein** |
|---|---|
| 2006.41 | |
| 2134.87 | |
| 2333.74 | |
| 2436.6 | |
| 2627.29 | |
| 2780.15 | |
| 3029.54 | |
| 3083.76 | |
| 3552.74 | |
| 3609.47 | |
| 3700.49 | |
| 4243.31 | |
| 4805.95 | |
| 4872.57 | |
| 4919.35 | |
| 5226.46 | |
| 5253.25 | |
| 5986.2 | |
| 6057.12 | |
| 6167.42 | |
| 6215.29 | |
| 7006.45 | |
| 7113.67 | |
| 7219.14 | |
| 7397.9 | |

Table S52 revealed that no ribosomal protein mass peak is present in the mass spectrum peak list of *Stenotrophomonas maltophilia* 5PC6

| **Table S53:** ***Stenotrophomonas maltophilia*** **Peak list of 15MF** | **Ribosomal protein** |
|---|---|
| 2005.81 | |
| 2135.08 | |
| 2266.95 | |
| 2392.6 | |
| 2428.95 | |
| 2635.06 | |
| 2780.14 | |
| 3049.99 | |
| 3521.91 | |
| 3540.97 | |
| 3586.11 | |
| 3686.97 | |
| 4242.56 | |
| 4435.47 | |
| 4447.42 | |
| 4796.13 | |
| 4856.24 | L36 |
| 5268.9 | |
| 5888.19 | |
| 5938.28 | |
| 6098.92 | |
| 6863.74 | |
| 7044.05 | |
| 7080.65 | |
| 7165.51 | L29 |
| 7372.67 | |
| 8484.04 | |
| 9315.71 | |
| 9582.44 | |

Table S53 shows the annotated ribosomal protein mass peaks of *S. maltophilia* 15MF. Specifically, ribosomal protein L36 and L29 mapped to mass peaks 4856.24 Da and 7165.51 Da, respectively.

| **Table S54:** ***Stenotrophomonas maltophilia*** **Peak list of 25MC6** | **Ribosomal protein** |
|---|---|
| 2004.93 | |
| 2061.68 | |
| 2082.36 | |
| 2101.74 | |
| 2134.09 | |
| 2263.09 | |
| 2391.93 | |
| 2428.14 | |
| 2634.05 | |
| 2778.97 | |
| 3035.27 | |
| 3530.65 | |
| 3583.78 | |
| 3685.58 | |
| 4222.56 | |
| 4240.39 | |
| 4789.41 | |
| 4836.77 | |
| 4854.7 | L36 |
| 5267.18 | |
| 5885.44 | |
| 6068.95 | |
| 6863.76 | |
| 7029.71 | |
| 7060.31 | |
| 7371.15 | |
| 8445.38 | |
| 9318.85 | |
| 9578.49 | |

Table S54 shows the annotated ribosomal protein mass peak of *S. maltophilia* 25MC6. Specifically, ribosomal protein L36 mapped to mass peak 4854.7 Da.

| **Table S55:** ***Stenotrophomonas maltophilia*** **Peak list of ATCC 13637** | **Ribosomal protein** |
|---|---|
| 2268.73 | |
| 2779.94 | |
| 3049.81 | |
| 4221.58 | |
| 4239.31 | |
| 4535.98 | |
| 4664.87 | |
| 4855.3 | L36 |
| 5157.45 | |
| 5267.81 | |
| 5884.05 | |
| 6097.53 | |
| 6240.64 | |
| 6858.62 | |
| 7040.89 | |
| 7080.04 | |
| 7147.27 | |
| 7169.2 | L29 or L32 |
| 7372.08 | |
| 8015.08 | |
| 8444.72 | |
| 9329 | |
| 9579.36 | |

Table S55 shows the annotated ribosomal protein mass peaks of *S maltophilia* ATCC 13637. Specifically, ribosomal protein L36 mapped to mass peak 4855.3 Da. On the other hand, ribosomal protein L29 or L32 could be mapped to mass peak 7169.2 Da.

| **Table S56:** ***Stenotrophomonas maltophilia*** **Peak list of Seab01** | **Ribosomal protein** |
|---|---|
| 2057.43 | |
| 2436.02 | |
| 2626.23 | |
| 2778.82 | |
| 2986.94 | |
| 3081.5 | |
| 3500.77 | |
| 3550.09 | |
| 3606.84 | |
| 3698.85 | |
| 4240.04 | |
| 4498.23 | |
| 4642.31 | |
| 4765.47 | |
| 4801.62 | |
| 4868.79 | |
| 4917.05 | |
| 5248.6 | |
| 5973.8 | |
| 6160.71 | |
| 6997.73 | |
| 7104.94 | |
| 7209.3 | |
| 7390.85 | |
| 8476.65 | |
| 8866.42 | |
| 8899.39 | |
| 8989.67 | |
| 9279.85 | |
| 9522.97 | |

Table S56 revealed that no ribosomal protein mass peak could be annotated for *S. maltophilia* Seab01.

| **Table S57:** ***Stenotrophomonas maltophilia*** **Peak list of Seab05** | **Ribosomal protein** |
|---|---|
| 2444.76 | |
| 2627.94 | |
| 2772.18 | |
| 3061.93 | |
| 3619.25 | |
| 3717.43 | |
| 4243.49 | |
| 4655.98 | |
| 4791.37 | |
| 4886.81 | |
| 5253.06 | |
| 5928.2 | |
| 6117.73 | |
| 6356.32 | |
| 6985.37 | |
| 7085.52 | |
| 7139.74 | |
| 7237.69 | |
| 7429.3 | |
| 8483.16 | |
| 9308.58 | |
| 9834.17 | |

Table S57 revealed that no ribosomal protein mass peak could be annotated for *S. maltophilia* Seab05.

| **Table S58:** ***Stenotrophomonas maltophilia*** **Peak list of Seab06** | **Ribosomal protein** |
|---|---|
| 2267.97 | |
| 2635.5 | |
| 2781.37 | |
| 3051.01 | |
| 4241.04 | |
| 4856.84 | L36 |
| 5229.91 | |
| 5268.82 | |
| 5884.12 | |
| 6099.94 | |
| 6228.65 | |
| 7043.99 | |
| 7165.54 | L29 or L32 |
| 7374.13 | |
| 8470.18 | |
| 8877.61 | |
| 9325.6 | |
| 9582.17 | |

Table S58 shows the annotated ribosomal protein mass peaks of *S. maltophilia* Seab06. Specifically, ribosomal protein L36 could be mapped to mass peak 4856.84 Da. On the other hand, either ribosomal protein L29 or L32 could be mapped to mass peak 7165.54 Da.

| **Table S59:** ***Stenotrophomonas maltophilia*** **Peak list of Seab08** | **Ribosomal protein** |
|---|---|
| 2267.68 | |
| 2780.83 | |
| 3050.46 | |
| 3687.24 | |
| 4242.23 | |
| 4787.85 | |
| 4797.62 | |
| 4856.03 | L36 |
| 5164.34 | |
| 5268.75 | |
| 5888.69 | |
| 5939.67 | |
| 6098.51 | |
| 6228.09 | |
| 6864.68 | |
| 7042.53 | |
| 7080.46 | |
| 7149.46 | |
| 7170.2 | L29 or L32 |
| 7372.81 | |
| 8015.99 | |
| 8462.74 | |
| 8879.75 | |
| 9317.65 | |
| 9580.76 | |

Table S59 shows the annotated ribosomal protein mass peaks of *S. maltophilia* Seab08. Specifically, ribosomal protein L36 mapped to mass peak 4856.03 Da. On the other hand, ribosomal protein L29 or L32 mapped to mass peak 7170.2 Da.

| **Table S60:** ***Bacillus cereus*** **Peak list of ATCC 9634** | **Ribosomal protein** |
|---|---|
| 2167.47 | |
| 2423.55 | |
| 2586.17 | |
| 2721.39 | |
| 2811.72 | |
| 3854.57 | |
| 4333.78 | L36 |
| 4994.33 | |
| 5171.29 | L34 |
| 5441.56 | |
| 5548.11 | |
| 5886.77 | |
| 6263.71 | |
| 6384.27 | |
| 6426.13 | |

Table S60 shows the annotated ribosomal protein mass peaks of *Bacillus cereus* ATCC 9634. Specifically, ribosomal protein L36 and L34 mapped to mass peaks 4333.78 Da and 5171.29 Da, respectively.

| **Table S61:** ***Bacillus cereus*** **Peak list of ATCC 14579** | **Ribosomal protein** |
|---|---|
| 2003.83 | |
| 2168.26 | |
| 2424.3 | |
| 2586.66 | |
| 2812.95 | |
| 3343.54 | |
| 3855.6 | |
| 4334.9 | L36 |
| 4995.39 | |
| 5172.04 | L34 |
| 5442.19 | |
| 5548.48 | |
| 5888.12 | |
| 6264.18 | |
| 6384.52 | |
| 6426.85 | |

| 6686.22 |
|---|

Table S61 shows the annotated ribosomal protein mass peaks of *B. cereus* ATCC 14579. Specifically, ribosomal protein L36 and L34 mapped to mass peaks 4334.9 Da and 5172.04 Da, respectively.

| **Table S62:** ***Bacillus cereus*** **Peak list of ATCC 14893** | **Ribosomal protein** |
|---|---|
| 2167.38 | |
| 2189 | |
| 2595.15 | |
| 2640.92 | |
| 2787.54 | |
| 3088.19 | |
| 3116.21 | |
| 3360.61 | |
| 3650.13 | |
| 3720.68 | |
| 3758.49 | |
| 4055.27 | |
| 4331.74 | L36 |
| 4548.99 | |
| 4991.31 | |
| 5293.82 | |
| 5446.26 | |
| 5546.57 | |

Table S62 shows the annotated ribosomal protein mass peak of *B. cereus* ATCC 14893. Specifically, ribosomal protein L36 mapped to mass peak 4331.74 Da.

| **Table S63:** ***Acinetobacter baumanii*** **Peak list of ATCC 15308** | **Ribosomal protein** |
|---|---|
| 2875.2 | |
| 4245.44 | |
| 4264.6 | L36 |
| 4662.53 | |
| 5176.07 | |
| 5748.39 | |
| 6092.22 | L33 |
| 6331.04 | |
| 6951.37 | |
| 7436.16 | L29 |
| 8325.19 | L31 |
| 8487.95 | |
| 9322.78 | |

Table S63 shows the annotated ribosomal protein mass peaks of *Acinetobacter baumanii* ATCC 15308. Specifically, ribosomal protein L36, L33, L29 and L31 mapped to mass peaks 4264.6 Da, 6092.22 Da, 7436.16 Da, 8325.19 Da, respectively.

| **Table S64:** ***Aeromonas hydrophila*** **Peak list of ATCC 7966** | **Ribosomal protein** |
|---|---|
| 2131.16 | |
| 2224.76 | |
| 2527.02 | |
| 3051.37 | |
| 3154.05 | |
| 3606.15 | |
| 3675.68 | |
| 3915.63 | |
| 4173.01 | |
| 4259.52 | |
| 4447.77 | |
| 4593.94 | |
| 4701.87 | |
| 5008.84 | |
| 5051.51 | L34 |
| 5157 | |
| 5316.3 | |
| 6100.64 | |
| 6305.38 | |
| 6481.4 | |
| 7210.33 | L29 |
| 7349.13 | |

| | |
|---|---|
| 7748.13 | L31 |
| 8344.02 | |
| 9185.71 | |
| 9401.75 | S17 |

Table S64 shows the annotated ribosomal protein mass peaks of *Aeromonas hydrophila* ATCC 7966. Specifically, ribosomal protein L34, L29, L31 and S17 mapped to mass peaks 5051.51 Da, 7210.33 Da, 7748.13 Da and 9401.75 Da, respectively.

| **Table S65: *Bacillus amyloliquefaciens* Peak list of ATCC 23842** | **Ribosomal protein** |
|---|---|
| 2186 | |
| 2202.78 | |
| 2287.12 | |
| 2293.97 | |
| 2445.25 | |
| 2495.14 | |
| 2502.15 | |
| 2647.21 | |
| 2687.69 | |
| 2729.6 | |
| 3051.35 | |
| 3662.96 | |
| 3755.78 | |
| 3860.49 | |
| 4289.3 | |
| 4307.27 | L36 |
| 4346.25 | |
| 4627.21 | |
| 5067.54 | |
| 5281.61 | |
| 5393.44 | |
| 5448.91 | |
| 5852.83 | |
| 6407.3 | |
| 6509.15 | |
| 6673.27 | |
| 7331.35 | |
| 7436.31 | L31 |
| 7714.92 | L29 |
| 9887.96 | |

Table S65 shows the annotated ribosomal protein mass peaks of *Bacillus amyloliquefaciens* ATCC 23842. Specifically, ribosomal protein L36, L31 and L29 mapped to mass peaks 4307.27 Da, 7436.31 Da, and 7714.92 Da, respectively.

| **Table S66:** ***Bacillus licheniformis*** **Peak list of ATCC 14580** | **Ribosomal protein** |
|---|---|
| 2057.59 | |
| 2946.81 | |
| 3021.98 | |
| 3040.3 | |
| 3252.33 | |
| 3289.98 | |
| 3725.92 | |
| 3864.18 | |
| 4304.36 | L36 |
| 4795.03 | |
| 4948.57 | |
| 5891.01 | |
| 6503.28 | |
| 7721.85 | L29 |
| 9585.41 | L31 Type B |
| 9899.77 | |

Table S66 shows the annotated ribosomal protein mass peaks of *Bacillus licheniformis* ATCC 14580. Specifically, ribosomal protein L36, L29 and L31 Type B mapped to mass peaks 4304.36 Da, 7721.85 Da, and 9585.41 Da, respectively.

| **Table S67:** ***Bacillus licheniformis*** **Peak list of ATCC 27811** | **Ribosomal protein** |
|---|---|
| 2056.34 | |
| 3021.59 | |
| 3039.44 | |
| 3252.4 | |
| 3289.37 | |
| 4795.72 | |
| 5893.43 | |
| 6503.32 | |
| 7038.26 | |
| 7418.63 | |
| 9581.65 | L31 Type B |
| 9898.53 | |

Table S67 shows the annotated ribosomal protein mass peaks of *B. licheniformis* ATCC 27811. Specifically, ribosomal protein L31 Type B mapped to mass peak 9581.65 Da.

| **Table S68:** ***Bacillus licheniformis*** **Peak list of 1Pesc** | **Ribosomal protein** |
|---|---|
| 2056.91 | |
| 2328.92 | |
| 2601.41 | |
| 2947.95 | |
| 3021.35 | |
| 3046 | |
| 3253.26 | |
| 3289.77 | |
| 3726.21 | |
| 4305.45 | L36 |
| 4817.06 | |
| 4984.85 | |
| 5894.26 | |
| 7258.66 | |
| 9902.06 | |

Table S68 shows the annotated ribosomal protein mass peaks of *B. licheniformis* 1Pesc. Specifically, ribosomal protein L36 mapped to mass peak 4305.45 Da.

| **Table S69: *Bacillus megaterium* Peak list of ATCC 14581** | **Ribosomal protein** |
|---|---|
| 2476.89 | |
| 3046.75 | |
| 3074.7 | |
| 3130.33 | |
| 3423.43 | |
| 3553.25 | |
| 4303.79 | |
| 4610.75 | |
| 4794.2 | |
| 5205.46 | L34 |
| 5830.22 | L33 |
| 6259.21 | |
| 6578.36 | |
| 6742.84 | |
| 7452.15 | |
| 7727.76 | L29 |
| 9346.43 | |

Table S69 shows the annotated ribosomal protein mass peaks of *Bacillus megaterium* ATCC 14581. Specifically, ribosomal protein L34, L33 and L29 mapped to mass peaks 5205.46 Da, 5830.22 Da, and 7727.76 Da, respectively.

| **Table S70:** ***Bacillus megaterium*** **Peak list of ATCC 25848** | **Ribosomal protein** |
|---|---|
| 2478.58 | |
| 3047.27 | |
| 3075.32 | |
| 3130.55 | |
| 3553.09 | |
| 4304.25 | |
| 4608.92 | |
| 4793.03 | |
| 4978.21 | |
| 5204.25 | L34 |
| 5829.56 | L33 |
| 6257.77 | |
| 6576.95 | |
| 6741.19 | |
| 7449.05 | |
| 7725.32 | L29 |
| 9349.63 | |

Table S70 shows the annotated ribosomal protein mass peaks of *B. megaterium* ATCC 25848. Specifically, ribosomal protein L34, L33 and L29 mapped to mass peaks 5204.25 Da, 5829.56 Da, 7725.32 Da, respectively.

| **Table S71:** ***Bacillus megaterium*** **Peak list of Rpesc** | **Ribosomal protein** |
|---|---|
| 2477.74 | |
| 3047.31 | |
| 3075.28 | |
| 3131.04 | |
| 3553.68 | |
| 4304.51 | |
| 4610.56 | |
| 4794.24 | |
| 4979.64 | |
| 5016.52 | |
| 5166.86 | |
| 5206.01 | L34 |
| 5831.56 | L33 |
| 6261.02 | |

| | |
|---|---|
| 6581.32 | |
| 6743.97 | |
| 7729.49 | L29 |
| 9348.91 | |

Table S71 shows the annotated ribosomal protein mass peaks of *B. megaterium* Rpesc. Specifically, ribosomal protein L34, L33 and L29 mapped to mass peaks 5206.01 Da, 5831.56 Da, and 7729.49 Da, respectively.

| **Table S72:** ***Bacillus pumilus*** **Peak list of ATCC 7061** | **Ribosomal protein** |
|---|---|
| 3022.31 | |
| 3045.46 | |
| 3619.52 | |
| 3761.22 | |
| 4066.56 | |
| 4303.61 | L36 |
| 4586.95 | |
| 5297.45 | |
| 6045.52 | |
| 6617.76 | |
| 6865.7 | |
| 7236.96 | |
| 7724.41 | L29 |
| 9821.49 | |

Table S72 shows the annotated ribosomal protein mass peaks of *Bacillus pumilus* ATCC 7061. Specifically, ribosomal protein L36 and L29 mapped to mass peaks 4303.61 Da and 7724.41 Da, respectively.

| **Table S73:** ***Bacillus pumilus*** **Peak list of ATCC 14884** | **Ribosomal protein** |
|---|---|
| 3016 | |
| 3043.96 | |
| 3605.25 | |
| 3620.43 | |
| 3760.13 | |
| 4302.11 | L36 |
| 4584.7 | |
| 5297.33 | |
| 6113.69 | |
| 6616.31 | |
| 6774.86 | |
| 6880.35 | |
| 7238.48 | |
| 7723.24 | L29 |
| 9819 | |

Table S73 shows the annotated ribosomal protein mass peaks of *B. pumilus* ATCC 14884. Specifically, ribosomal protein L36 and L29 mapped to mass peaks 4302.11 Da and 7723.24 Da, respectively.

| **Table S74:** ***Carnobacterium divergens*** **Peak list of ATCC 35677** | **Ribosomal protein** |
|---|---|
| 2172.69 | |
| 2718.73 | |
| 2860.78 | |
| 3230.26 | |
| 3348.71 | |
| 3420.77 | |
| 3764.16 | |
| 4344.88 | |
| 4838.75 | |
| 5721.73 | |
| 6460.23 | |
| 6697.89 | |

Table S74 revealed that no ribosomal protein mass peak could be annotated for *Carnobacterium divergens* ATCC 35677.

| **Table S75:** ***Carnobacterium gallinarum*** **Peak list of ATCC 49517** | **Ribosomal protein** |
|---|---|
| 2173.76 | |
| 2838.42 | |
| 3239.54 | |
| 3350.19 | |
| 3730.95 | |
| 4346.29 | |
| 5675.6 | |
| 6346.46 | |
| 6476.94 | L30 |
| 6699.44 | |
| 6963.96 | |
| 7427.78 | |

Table S75 shows the annotated ribosomal protein mass peaks of *Carnobacterium gallinarum* ATCC 49517. Specifically, ribosomal protein L30 mapped to mass peak 6476.94 Da. The annotation exercise was performed with a proteome of *Carnobacterium* sp. CP1 from UniProt as a corresponding proteome of *Carnobacterium gallinarum* could not be found on UniProt.

| **Table S76:** ***Citrobacter freundii*** **Peak list of ATCC 8090** | **Ribosomal protein** |
|---|---|
| 2183.63 | |
| 2620.86 | |
| 2705.95 | |
| 2826.39 | |
| 3128.53 | |
| 3150.7 | |
| 3192.38 | |
| 3545.79 | |
| 3631.29 | |
| 4185.61 | |
| 4364.73 | L36 |
| 4452.67 | |
| 4483.74 | |
| 4599.12 | |
| 4762.27 | |
| 4777.8 | |
| 5144.52 | |
| 5239.56 | |
| 5409.9 | L34 |
| 6254.96 | |
| 6299.39 | |
| 6383.34 | |
| 7089.74 | |
| 7260.89 | L29 |
| 7736.84 | |
| 7869.19 | |
| 8369.49 | |
| 8906.16 | |
| 8965.26 | |
| 9193.26 | S16 |
| 9526.77 | |

Table S76 shows the annotated ribosomal protein mass peaks of *Citrobacter freundii* ATCC 8090. Specifically, ribosomal protein mass peak L36, L34, L29 and S16 mapped to mass peaks 4364.73 Da, 5409.9 Da, 7260.89 Da, and 9193.26 Da, respectively.

| **Table S77:** ***Clostridium botulinum*** **Peak list of ATCC 19397** | **Ribosomal protein** |
|---|---|
| 2152.3 | |
| 2463.48 | |
| 2752.27 | |
| 2949.91 | |
| 3296.04 | |
| 3328.93 | |
| 3378.17 | |
| 3488.93 | |
| 3657.64 | |
| 3687.08 | |
| 3700.25 | |
| 3856.71 | |
| 3878.61 | |
| 4162.26 | |
| 4303.69 | |
| 5491.2 | |
| 5503.65 | L34 |
| 5899.63 | L33 |
| 6370.96 | |
| 6472.68 | |
| 6591.99 | |
| 6657.99 | |
| 6756.25 | |
| 6942.99 | |
| 6977.97 | |
| 7374.28 | |
| 8325.17 | L29 |

Table S77 shows the annotated ribosomal protein mass peaks of *Clostridium botulinum* ATCC 19397. Specifically, ribosomal protein L34, L33 and L29 mapped to mass peaks 5503.65 Da, 5899.63 Da, and 8325.17 Da, respectively.

| **Table S78: *Clostridium perfringens* Peak list of ATCC 10543** | **Ribosomal protein** |
|---|---|
| 2152.66 | |
| 2745.87 | |
| 2997.39 | |
| 3153.57 | |
| 3349.72 | |
| 3463.12 | |
| 3469.6 | |
| 3661.53 | |
| 3859.46 | |
| 3916.42 | |
| 4066.35 | |
| 4304.08 | L36 |
| 4684.12 | |
| 5490.91 | L34 |
| 5995.75 | |
| 6306.36 | L30 |
| 6699.01 | |
| 6937.27 | |
| 7322.45 | |
| 7718.7 | L31 |
| 7832.65 | |
| 8132.71 | L29 |
| 8965.47 | |
| 9368.1 | |

Table S78 shows the annotated ribosomal protein mass peaks of *Clostridium perfringens* ATCC 10543. Specifically, ribosomal protein L36, L34, L30, L31 and L29 mapped to mass peaks 4304.08 Da, 5490.91 Da, 6306.36 Da, 7718.7 Da, and 8132.71 Da, respectively.

| **Table S79: *Enterobacter aerogenes* Peak list of ATCC 13048** | **Ribosomal protein** |
|---|---|
| 2181.58 | |
| 2696.04 | |
| 2855.22 | |
| 3143.38 | |
| 3147.99 | |
| 3577.69 | |
| 3620.97 | |
| 3822.46 | |
| 4183.75 | |
| 4362.15 | |
| 4445.83 | |
| 4471.57 | |
| 4567.56 | |
| 4737.96 | |
| 4760.8 | |
| 5142.28 | |
| 5391.39 | |
| 6289.19 | |
| 6378.64 | L33 |
| 6822.06 | |
| 7154.22 | |
| 7239.91 | |
| 7352.47 | |
| 7645.81 | |
| 9478.24 | |

Table S79 shows the annotated ribosomal protein mass peak of *Enterobacter aerogenes* ATCC 13048. Specifically, ribosomal protein L33 mapped to mass peak 6378.64 Da. Annotation of the peak list of the bacterium was done with the proteome of *Enterobacter* sp. CC120223-11 from UniProt as no corresponding proteome for *E. aerogenes* could be found on UniProt.

| **Table S80:** ***Enterobacter cloacae*** **Peak list of ATCC 13047** | **Ribosomal protein** |
|---|---|
| 2183.08 | |
| 2478.43 | |
| 2558.16 | |
| 2691.03 | |
| 3137.08 | |
| 3165.97 | |
| 3579.87 | |
| 3622.46 | |
| 4185.71 | |
| 4364.45 | |
| 4448.15 | |
| 4569.9 | |
| 4756.71 | |
| 5115.21 | |
| 5152.04 | |
| 5380.26 | L34 |
| 6152.08 | |
| 6273.4 | |
| 6329.65 | |
| 6446.42 | |
| 7157.81 | |
| 7242.39 | L29 |
| 9139.18 | L27 |
| 9510.19 | |

Table S80 shows the annotated ribosomal protein mass peaks of *Enterobacter cloacae* ATCC 13047. Specifically, ribosomal protein L34, L29 and L27 mapped to mass peaks 5380.26 Da, 7242.39 Da, and 9139.18 Da, respectively.

| **Table S81:** ***Enterobacter hormaechei*** **Peak list of 10MC1** | **Ribosomal protein** |
|---|---|
| 2182.95 | |
| 2322.87 | |
| 2463.61 | |
| 2691.03 | |
| 2801.92 | |
| 2821.95 | |
| 3188.09 | |
| 3623.67 | |
| 3844.44 | |
| 4165.08 | |
| 4184.89 | |
| 4364.3 | |
| 4502.57 | |
| 4544.85 | |
| 4644.62 | |
| 4740.71 | |
| 4755.48 | |
| 5119.7 | |
| 5380.48 | L34 |
| 5643.17 | |
| 6255.65 | |
| 6329.9 | |
| 7242.77 | L29 |
| 7689.46 | |
| 8329.34 | |
| 9502.25 | |

Table S81 shows the annotated ribosomal protein mass peaks of *Enterobacter hormaechei* 10MC1. Specifically, ribosomal protein L34 and L29 mapped to mass peaks 5380.48 Da and 7242.77 Da, respectively.

| **Table S82:** ***Enterobacter sakazakii*** **Peak list of ATCC 29544** | **Ribosomal protein** |
|---|---|
| 2183.3 | |
| 2339.03 | |
| 2438.22 | |
| 2690.99 | |
| 2825.84 | |
| 3127.88 | |
| 3159.95 | |
| 3537.08 | |
| 3579.24 | |
| 3645.06 | |
| 3659.09 | |
| 4186.77 | |
| 4363.82 | |
| 4739.59 | |
| 5144.78 | |
| 5380.87 | |
| 6255.54 | |
| 6316.76 | |
| 6383.65 | |
| 7039.79 | |
| 7159 | |
| 7289.18 | |
| 8171.88 | |
| 9478.54 | |

Table S82 revealed that no ribosomal protein mass peak could be annotated for *Enterobacter sakazakii* ATCC 29544. The proteome used for the annotation exercise was from the UniProt entry of *Enterobacter* sp. CC120223-11 as no corresponding proteome could be found for *Enterobacter sakazakii* in UniProt.

| **Table S83:** ***Hafnia alvei*** **Peak list of ATCC 9760** | **Ribosomal protein** |
|---|---|
| 2189.47 | |
| 2698.41 | |
| 2811.51 | |
| 3111.1 | |
| 3127.67 | |
| 3199.71 | |
| 3545.2 | |
| 3616.99 | |
| 3889.33 | |
| 4185.06 | |
| 4342.7 | |
| 4376.91 | L36 |
| 4778.06 | |
| 4823.23 | |
| 5129.07 | |
| 5352.65 | |
| 5395.14 | L34 |
| 6223.63 | |
| 6253.2 | |
| 6397 | |
| 7087.68 | |
| 7232.04 | L29 |
| 7368.47 | |
| 7775.42 | |
| 9554.54 | |

Table S83 shows the annotated ribosomal protein mass peaks of *Hafnia alvei* ATCC 9760. Specifically, ribosomal protein L36, L34 and L29 mapped to mass peaks 4376.91 Da, 5395.14 Da, and 7232.04 Da, respectively.

| **Table S84:** ***Klebsiella oxytoca*** **Peak list of ATCC 13182** | **Ribosomal protein** |
|---|---|
| 2181.57 | |
| 2704.61 | |
| 2837.44 | |
| 3126.58 | |
| 3133.15 | |
| 3412.33 | |
| 3577.73 | |
| 3621.25 | |
| 3843.31 | |
| 4135.6 | |
| 4184.27 | |
| 4362.23 | L36 |
| 4447.21 | |
| 4568.61 | |
| 4689.32 | |
| 4731.42 | |
| 4775.84 | |
| 5050.7 | |
| 5081.38 | |
| 5142.55 | |
| 5407.96 | L34 |
| 5618.21 | |
| 6255.4 | |
| 6382.18 | L33 |
| 6822.89 | |
| 7111.14 | |
| 7151.31 | |
| 7241.09 | L29 |
| 7346 | |
| 7686.2 | |
| 8273.47 | |
| 8368.06 | |
| 9378.81 | |
| 9462.87 | |

Table S84 shows the annotated ribosomal protein mass peaks of *Klebsiella oxytoca* ATCC 13182. Specifically, ribosomal protein L36, L34, L33 and L29 mapped to mass peaks 4362.23 Da, 5407.96 Da, 6382.18 Da, and 7241.09 Da, respectively.

| **Table S85:** ***Klebsiella pneumoniae*** **Peak list of ATCC 10031** | **Ribosomal protein** |
|---|---|
| 2182.65 | |
| 2690.7 | |
| 2856.48 | |
| 3144.7 | |
| 3148.96 | |
| 3579.76 | |
| 3622.33 | |
| 4185.39 | |
| 4364.08 | |
| 4739.86 | |
| 4770.53 | |
| 5380.17 | L34 |
| 6291.81 | |
| 6382.06 | |
| 7158.62 | |
| 7242.06 | L29 |
| 7706.43 | |
| 9480.92 | |

Table S85 shows the annotated ribosomal protein mass peaks of *Klebsiella pneumoniae* ATCC 10031. Specifically, ribosomal protein L34 and L29 mapped to mass peaks 5380.17 Da and 7242.06 Da, respectively.

| **Table S86:** ***Listeria innocua*** **Peak list of ATCC 33090** | **Ribosomal protein** |
|---|---|
| 2162.69 | |
| 2756.44 | |
| 3004.48 | |
| 3181.91 | |
| 3194.04 | |
| 3358.48 | |
| 3431.08 | |
| 3508.13 | |
| 3701.88 | |
| 4322.86 | L36 |
| 4519.78 | |
| 4696.36 | |
| 4876.56 | |
| 5127.33 | |
| 5173.21 | |
| 5299.88 | L34 |
| 5597.96 | |
| 5856.87 | |
| 6007.16 | L33 |
| 6196.63 | |
| 6362.08 | L32 |
| 6386.2 | |
| 6423.77 | |
| 6715.29 | |
| 6860.47 | |
| 7014.05 | |
| 7402.41 | L29 |
| 9036.85 | |
| 9390.35 | |
| 9751.87 | |

Table S86 shows the annotated ribosomal protein mass peaks of *Listeria innocua* ATCC 33090. Specifically, ribosomal protein L36, L34, L33, L32 and L29 mapped to mass peaks 4322.86 Da, 5299.88 Da, 6007.16 Da, 6362.08 Da, and 7402.41 Da, respectively.

| **Table S87:** ***Listeria ivanovii*** **Peak list of ATCC 19119** | **Ribosomal protein** |
|---|---|
| 2134.24 | |
| 2162.57 | |
| 2722.39 | |
| 3004.22 | |
| 3013.42 | |
| 3181.77 | |
| 3193.81 | |
| 3251.44 | |
| 3430.89 | |
| 3442.75 | |
| 3448.61 | |
| 3508.31 | |
| 3701.58 | |
| 3993.67 | |
| 4043.74 | |
| 4322.64 | L36 |
| 4519.27 | |
| 4696.02 | |
| 4876.22 | |
| 5117.45 | |
| 5172.79 | |
| 5597.52 | |
| 6006.78 | |
| 6124.8 | |
| 6361.38 | |
| 6385.31 | |
| 6698.25 | |
| 6860.16 | |
| 7014.45 | |
| 7401.65 | L29 |
| 7986.17 | |
| 8085.84 | |
| 9036.79 | |
| 9390.25 | |
| 9750.85 | |

Table S87 shows the annotated ribosomal protein mass peaks of *Listeria ivanovii* ATCC 19119. Specifically, ribosomal protein L36 and L29 mapped to mass peaks 4322.64 Da and 7401.65 Da, respectively.

| **Table S88:** ***Listeria monocytogenes*** **Peak list of CECT 4032** | **Ribosomal protein** |
|---|---|
| 2163.72 | |
| 2757.23 | |
| 3005.3 | |
| 3182.56 | |
| 3195.6 | |
| 3213.51 | |
| 3359.13 | |
| 3431.75 | |
| 3509.5 | |
| 3702.51 | |
| 3973.21 | |
| 4324.63 | L36 |
| 4520.35 | |
| 4696.91 | |
| 4877.37 | |
| 5118.89 | |
| 5173.83 | |
| 5598.56 | |
| 6007.98 | L33 |
| 6362.58 | L32 |
| 6388.62 | |
| 6424.85 | |
| 6715.83 | |
| 6860.97 | |
| 7016.02 | |
| 7402.5 | L29 |
| 7943.86 | |
| 8087.25 | |
| 9036.02 | |
| 9389.26 | |
| 9751.72 | |

Table S88 shows the annotated ribosomal protein mass peaks of *Listeria monocytogenes* CECT 4032. Specifically, ribosomal protein L36, L33, L32 and L29 mapped to mass peaks 4323.63 Da, 6007.98 Da, 6362.58 Da and 7402.5 Da, respectively.

| **Table S89:** ***Listeria seeligeri*** **Peak list of ATCC 35967** | **Ribosomal protein** |
|---|---|
| 2028.17 | |
| 2071.51 | |
| 2277.18 | |
| 2432.74 | |
| 2474.45 | |
| 2722.66 | |
| 2828.69 | |
| 2982.69 | |
| 3004.45 | |
| 3181.87 | |
| 3701.92 | |
| 4323.25 | L36 |
| 4548.45 | |
| 4696.17 | |
| 4876.81 | |
| 4942.06 | |
| 5118.37 | |
| 5172.78 | |
| 5598.85 | |
| 6007.23 | L33 |
| 6361.3 | |
| 6521.49 | L32 |
| 6860.49 | |
| 7014.45 | |
| 7401.77 | L29 |
| 7929.17 | |
| 8095.29 | |
| 9094.1 | S18 |
| 9389 | |
| 9750.32 | |
| 9882.43 | |

Table S89 shows the annotated ribosomal protein mass peaks of *Listeria seeligeri* ATCC 35967. Specifically, ribosomal protein L36, L33, L32, L29 and S18 mapped to mass peaks 4323.25 Da, 6007.23 Da, 6521.49 Da, 7401.77 Da and 9094.1 Da, respectively.

| **Table S90:** ***Listeria welshimeri*** **Peak list of ATCC 35897** | **Ribosomal protein** |
|---|---|
| 2162.87 | |
| 2756.34 | |
| 3004.74 | |
| 3195.77 | |
| 3214.43 | |
| 3430.95 | |
| 3508.55 | |
| 3701.65 | |
| 3977.32 | |
| 4043.96 | |
| 4323.26 | L36 |
| 4512.29 | |
| 4696.03 | |
| 4876.22 | |
| 5117.87 | |
| 5172.8 | |
| 5597.81 | |
| 6006.74 | L33 |
| 6195.91 | |
| 6390.18 | |
| 6427.16 | |
| 6859.87 | |
| 7014.75 | |
| 7401.61 | L29 |
| 7952.55 | |
| 8086.26 | |
| 9021.54 | |
| 9388.35 | |
| 9750.59 | |

Table S90 shows the annotated ribosomal protein mass peaks of *Listeria welshimeri* ATCC 35897. Specifically, ribosomal protein L36, L33 and L29 mapped to mass peaks 4323.26 Da, 6006.74 Da, and 7401.61 Da, respectively.

| *Table S91: Morganella morganii* **Peak list of ATCC 8076** | **Ribosomal protein** |
|---|---|
| 2187.46 | |
| 2691.61 | |
| 3109.43 | |
| 3156.25 | |
| 3177.2 | |
| 3192.76 | |
| 3241.71 | |
| 3590.59 | |
| 3638.05 | |
| 3880.02 | |
| 4170.76 | |
| 4337.33 | |
| 4372.48 | L36 |
| 4454.86 | |
| 4642.51 | |
| 4733.74 | |
| 4963.2 | |
| 5138.23 | |
| 5381 | L34 |
| 5965.6 | |
| 6216.61 | |
| 6350.14 | L32 |
| 6481.24 | |
| 6615.61 | L30 |
| 6747.82 | |
| 7179.16 | |
| 7273.77 | L29 |
| 7758.06 | L31 |
| 7908.5 | |
| 8337.07 | |
| 8671.05 | |
| 9280.66 | |
| 9462.88 | |
| 9921.17 | |

Table S91 shows the annotated ribosomal protein mass peaks of *Morganella morganii* ATCC 8076. Specifically, ribosomal protein L36, L34, L32, L30, L29 and L31 mapped to mass peaks 4372.48 Da, 5381 Da, 6350.14 Da, 6615.61 Da, 7273.77 Da, and 7758.06 Da, respectively.

| **Table S92:** ***Morganella morganii*** **Peak list of ATCC BM65** | **Ribosomal protein** |
|---|---|
| 2186.57 | |
| 2690.45 | |
| 3108.37 | |
| 3155.21 | |
| 3175.8 | |
| 3192.47 | |
| 3240.5 | |
| 3589.28 | |
| 3636.59 | |
| 3878.45 | |
| 4335.02 | |
| 4371.15 | L36 |
| 4453.82 | |
| 4640.09 | |
| 4732.07 | |
| 5136.47 | |
| 5379.34 | L34 |
| 6214.54 | |
| 6347.6 | L32 |
| 6382.01 | |
| 6479.69 | L33 |
| 7175.84 | |
| 7270.22 | L29 |
| 8668.48 | |
| 9277.02 | |
| 9460.04 | |

Table S92 shows the annotated ribosomal protein mass peaks of *M. morganii* ATCC BM65. Specifically, ribosomal protein L36, L34, L32, L33 and L29 mapped to mass peaks 4371.15 Da, 5379.34 Da, 6347.6 Da, 6479.69 Da and 7270.22 Da, respectively.

| **Table S93:** ***Pantoea agglomerans*** **Peak list of ATCC 27155** | **Ribosomal protein** |
|---|---|
| 2510.23 | |
| 2699.25 | |
| 2771.15 | |
| 3128.05 | |
| 3590.45 | |
| 3695.83 | |
| 4005.29 | |
| 4108.72 | |
| 4176.64 | |
| 4182.2 | |
| 4420.52 | |
| 4660.21 | |
| 4754.21 | |
| 4890.73 | |
| 5018.4 | |
| 5396.3 | L34 |
| 6199.33 | |
| 6253.54 | |
| 6400.1 | |
| 7105.12 | |
| 7179.6 | |
| 7263.04 | L29 |
| 7390.27 | |
| 7986 | L31 |
| 8351.5 | |
| 9320.21 | |
| 9508.25 | |
| 9782.87 | |

Table S93 shows the annotated ribosomal protein mass peaks of *Pantoea agglomerans* ATCC 27155. Specifically, ribosomal protein L34, L29 and L31 mapped to mass peaks 5396.3 Da, 7263.04 Da, and 7986 Da, respectively.

| **Table S94:** ***Photobacterium damselae*** **Peak list of ATCC 33539** | **Ribosomal protein** |
|---|---|
| 2139.73 | |
| 2382.57 | |
| 2561.64 | |
| 2981.52 | |
| 3038.31 | |
| 3130.58 | |
| 3141.1 | |
| 3303.42 | |
| 3582.41 | |
| 3587.28 | |
| 4183.63 | |
| 4277.7 | L36 |
| 4517.77 | |
| 4723.07 | |
| 4772.76 | |
| 5121.43 | |
| 5161.36 | |
| 5485.5 | |
| 5690.25 | |
| 6075.32 | |
| 6259.52 | |
| 6280.49 | |
| 6582.65 | |
| 6605.32 | |
| 7168.09 | L29 |
| 7758.23 | L31 |
| 8365.37 | |
| 9033.06 | L28 |
| 9449.53 | |
| 9543.19 | |

Table S94 shows the annotated ribosomal protein mass peaks of *Photobacterium damselae* ATCC 33539. Specifically, ribosomal protein L36, L29, L31 and L28 mapped to mass peaks 4277.7 Da, 7168.09 Da, 7758.23 Da, and 9033.06 Da, respectively.

| **Table S95:** ***Photobacterium phosphoreum*** **Peak list of CECT 4172** | **Ribosomal protein** |
|---|---|
| 2140.52 | |
| 2433.12 | |
| 2578.7 | |
| 2982.52 | |
| 3010.37 | |
| 3131.5 | |
| 3138.37 | |
| 3560.05 | |
| 3581.59 | |
| 4184.69 | |
| 4278.41 | |
| 4481.23 | |
| 4490.72 | |
| 4575.63 | |
| 4737.57 | |
| 4803.05 | |
| 5073.23 | |
| 5156.34 | L34 |
| 5395.66 | |
| 5466.41 | |
| 5683.88 | |
| 6018.65 | |
| 6234.1 | |
| 6262.58 | |
| 6583.43 | |
| 7118.2 | L29 |
| 7161.16 | |
| 7862.98 | |
| 8367.83 | |
| 8806.01 | |
| 8975.52 | |
| 9148.31 | S16 |
| 9472.26 | |
| 9604.44 | S17 |

Table S95 shows the annotated ribosomal protein mass peaks of *Photobacterium phosphoreum* CECT 4172. Specifically, ribosomal protein L34, L29, S16 and S17 mapped to mass peaks 5156.34 Da, 7188.2 Da, 9148.31 Da, and 9604.44 Da, respectively.

| **Table S96:** ***Proteus mirabilis*** **Peak list of ATCC 14153** | **Ribosomal protein** |
|---|---|
| 2236.37 | |
| 2748.55 | |
| 2826.37 | |
| 2997.15 | |
| 3127.77 | |
| 3561.75 | |
| 3637.7 | |
| 3915.83 | |
| 3982.71 | |
| 4185.48 | |
| 4470.95 | L36 |
| 4738.01 | |
| 4796.59 | |
| 5131.01 | |
| 5495.52 | L34 |
| 5992.79 | |
| 6146.77 | |
| 6253.77 | |
| 6475.45 | |
| 7121.56 | |
| 7273.63 | L29 |
| 7829.73 | L31 |
| 7964.1 | |
| 8364.31 | |
| 9474.73 | |

Table S96 shows the annotated ribosomal protein mass peaks of *Proteus mirabilis* ATCC 14153. Specifically, ribosomal protein L36, L34, L29 and L31 mapped to mass peaks 4470.95 Da, 5495.52 Da, 7273.63 Da, and 7829.73 Da, respectively.

| **Table S97:** ***Proteus penneri*** **Peak list of ATCC 33519** | **Ribosomal protein** |
|---|---|
| 2241.71 | |
| 2747.31 | |
| 2825.18 | |
| 3028.37 | |
| 3130.59 | |
| 3136.46 | |
| 3195.99 | |
| 3553.49 | |
| 3636.47 | |
| 3905.65 | |
| 3981.9 | |
| 4184.18 | |
| 4482.28 | |
| 4794.57 | |
| 5129.53 | |
| 5493.54 | L34 |
| 6055.66 | |
| 6264.83 | |
| 6390.71 | L33 |
| 7106.43 | |
| 7270.68 | L29 |
| 7962.88 | |
| 8368.97 | |

Table S97 shows the annotated ribosomal protein mass peaks of *Proteus penneri* ATCC 33519. Specifically, ribosomal protein L34, L33 and L29 mapped to mass peaks 5493.54 Da, 6390.71 Da, and 7270.68 Da, respectively.

| **Table S98: *Providencia rettgeri* Peak list of ATCC 29944** | **Ribosomal protein** |
|---|---|
| 2219.17 | |
| 2734.9 | |
| 2854.46 | |
| 3115.78 | |
| 3158.57 | |
| 3554.44 | |
| 3622.94 | |
| 4029.43 | |
| 4185.01 | |
| 4435.36 | L36 |
| 4761.32 | |
| 4780.7 | |
| 5149.63 | |
| 5467.11 | L34 |
| 6186.41 | |
| 6229.49 | |
| 6314.47 | |
| 6440.37 | L33 |
| 6742.58 | |
| 6873.65 | |
| 7106.57 | |
| 7244.11 | L29 or L35 |
| 7786.93 | L31 |
| 8057.98 | |
| 8369.41 | |
| 8660.38 | |
| 9522.06 | |

Table S98 shows the annotated ribosomal protein mass peaks of *Providencia rettgeri* ATCC 29944. Specifically, ribosomal protein L36 L34, L33 and L31 mapped to mass peaks 4435.36 Da, 5467.11 Da, 6440.37 Da, and 7786.93 Da, respectively. On the other hand, ribosomal protein L29 or L35 could be mapped to mass peak 7244.11 Da.

| **Table S99:** ***Providencia stuartii*** **Peak list of ATCC 29914** | **Ribosomal protein** |
|---|---|
| 2218.13 | |
| 2734.1 | |
| 2886.45 | |
| 3122.49 | |
| 3167.13 | |
| 3554.21 | |
| 3622.79 | |
| 4143.85 | |
| 4185.47 | |
| 4435.29 | |
| 4489.45 | |
| 4774.35 | |
| 5467.31 | L34 |
| 6231.44 | |
| 6332.47 | |
| 6457.69 | L33 |
| 7107.03 | |
| 7243.38 | L29 or L35 |
| 8288.09 | |
| 8980.46 | S18 |

Table S99 shows the annotated ribosomal protein mass peaks of *Providencia stuartii* ATCC 29914. Specifically, ribosomal protein L34, L33 and S18 mapped to mass peaks 5467.31 Da, 6457.69 Da, and 8980.46 Da, respectively. On the other hand, ribosomal protein L29 or L35 could be mapped to mass peak 7243.38 Da.

| **Table S100:** ***Raoultella planticola*** **Peak list of ATCC 33531** | **Ribosomal protein** |
|---|---|
| 2182.61 | |
| 2705.61 | |
| 2838.33 | |
| 3127.71 | |
| 3150.43 | |
| 3190.81 | |
| 3578.71 | |
| 3622.32 | |
| 3660.75 | |
| 4183.77 | |
| 4363.68 | L36 |
| 4446.67 | |
| 4567.63 | |
| 4732.53 | |
| 4777.38 | |
| 5145.23 | |
| 5409.56 | L34 |
| 6254.15 | |
| 6297.93 | |
| 6381.19 | L33 |
| 7154.78 | |
| 7242.17 | L29 |
| 7521 | |
| 7913.81 | L31 |
| 8366.56 | |
| 9129.56 | S16 or L27 |
| 9464.08 | |
| 9552.5 | |

Table S100 shows the annotated ribosomal protein mass peaks of *Raoultella planticola* ATCC 33531. Specifically, ribosomal protein L36, L34, L33, L29 and L31 mapped to mass peaks 4363.68 Da, 5409.56 Da, 6381.19 Da, 7242.17 Da, and 7913.81 Da. On the other hand, ribosomal protein S16 or L27 could be mapped to mass peak 9129.56 Da.

| **Table S101:** ***Serratia liquefaciens*** **Peak list of ATCC 12926** | **Ribosomal protein** |
|---|---|
| 2176.63 | |
| 2700.07 | |
| 2827.1 | |
| 3031.97 | |
| 3121.9 | |
| 3189.17 | |
| 3605.9 | |
| 3652.38 | |
| 3962.8 | |
| 4186.13 | |
| 4350.66 | L36 |
| 4634.08 | |
| 4664.16 | |
| 4688.72 | |
| 4782.92 | |
| 5158.34 | |
| 5398.86 | L34 |
| 5536.11 | |
| 6062.54 | |
| 6241.87 | |
| 7210.58 | |
| 7303.02 | L29 |
| 7924.64 | |
| 8110.37 | |
| 8371.84 | |
| 9286.77 | |
| 9327.85 | |
| 9565.17 | |

Table S101 shows the annotated ribosomal protein mass peaks of *Serratia liquefaciens* ATCC 12926. Specifically, ribosomal protein L36, L34 and L29 mapped to mass peaks 4350.66 Da, 5398.86 Da, and 7303.02 Da, respectively.

| **Table S102:** ***Shewanella algae*** **Peak list of ATCC 51192** | **Ribosomal protein** |
|---|---|
| 2133.56 | |
| 2527.22 | |
| 3097.86 | |
| 3145.04 | |
| 3269.16 | |
| 3285.56 | |
| 3295.27 | |
| 3578.91 | |
| 3649.28 | |
| 4108.09 | |
| 4264.16 | |
| 4489.19 | |
| 4716.62 | |
| 4804.27 | |
| 4813.05 | |
| 5033.1 | |
| 5051.61 | |
| 5155.6 | |
| 5633.08 | |
| 6193.6 | |
| 6536.28 | |
| 6574.74 | |
| 6587.72 | |
| 6645.04 | |
| 7095.47 | |
| 7155.46 | L29 |
| 7295.94 | |
| 8215.4 | |
| 9431.51 | |
| 9622.25 | |

Table S102 shows the annotated ribosomal protein mass peak of *Shewanella algae* ATCC 51192. Specifically, ribosomal protein L29 mapped to mass peak 7155.46 Da.

| **Table S103: *Shewanella baltica* Peak list of CECT 323** | **Ribosomal protein** |
|---|---|
| 2132.85 | |
| 2512.15 | |
| 3027.5 | |
| 3084.79 | |
| 3276.55 | |
| 3308.64 | |
| 3574.29 | |
| 3626.47 | |
| 4107.32 | |
| 4263.21 | L36 |
| 4491.08 | |
| 4625.71 | |
| 4723.2 | |
| 4779.95 | |
| 5021.79 | |
| 5170.75 | |
| 5431.97 | |
| 5583.37 | |
| 5616.78 | |
| 6167.59 | |
| 6551.25 | |
| 6615.39 | |
| 7146.62 | L29 |
| 7251.07 | |
| 7560.92 | L31 |
| 8213.12 | |
| 8408.34 | |
| 9445.13 | |
| 9558.03 | |

Table S103 shows the annotated ribosomal protein mass peaks of *Shewanella baltica* CECT 323. Specifically, ribosomal protein L36, L29 and L31 mapped to mass peaks 4263.21 Da, 7146.62 Da, and 7560.92 Da, respectively.

| **Table S104:** ***Shewanella putrefaciens*** **Peak list of ATCC 8071** | **Ribosomal protein** |
|---|---|
| 2133.27 | |
| 2512.35 | |
| 3084.08 | |
| 3301.8 | |
| 3355.83 | |
| 3586.1 | |
| 3640.81 | |
| 4107.38 | |
| 4263.6 | L36 |
| 4491.07 | |
| 4626.04 | |
| 4723.78 | |
| 4795.53 | |
| 5022.15 | |
| 5171.48 | |
| 5432.86 | |
| 5590.89 | |
| 5638.89 | |
| 6168.05 | |
| 6552.37 | |
| 6601.8 | |
| 7170.32 | L29 |
| 7279.52 | |
| 7580.8 | |
| 8214.01 | |
| 9446.45 | |
| 9588.81 | |

Table S104 shows the annotated ribosomal protein mass peaks of *Shewanella putrefaciens* ATCC 8071. Specifically, ribosomal protein L36 and L29 mapped to mass peaks 4263.6 Da and 7170.32 Da, respectively.

| **Table S105:** ***Staphylococcus epidermidis*** **Peak list of ATCC 35983** | **Ribosomal protein** |
|---|---|
| 2556.18 | |
| 2838.22 | |
| 3739.55 | |
| 3777.36 | |
| 3823.73 | |
| 3952.08 | |
| 4289.04 | L36 |
| 5109.04 | |
| 5147.1 | |
| 5248.49 | |
| 5335.92 | |
| 6677.17 | |

Table S105 shows the annotated ribosomal protein mass peak of *Staphylococcus epidermidis* ATCC 35983. Specifically, ribosomal protein L36 mapped to mass peak 4289.04 Da.

| **Table S106:** ***Staphylococcus pasteuri*** **Peak list of 24MF** | **Ribosomal protein** |
|---|---|
| 2481.26 | |
| 2727.54 | |
| 3088.27 | |
| 3856.11 | |
| 4303.53 | L36 |
| 4962.04 | |
| 5454.75 | |
| 6176.51 | |

Table S106 shows the annotated ribosomal protein mass peak of *Staphylococcus pasteuri* 24MF. Specifically, ribosomal protein L36 mapped to mass peak 4303.53 Da.

| **Table S107:** ***Staphylococcus xylosus*** **Peak list of ATCC 29971** | **Ribosomal protein** |
|---|---|
| 2447.54 | |
| 2481.14 | |
| 2793.02 | |
| 3031.55 | |
| 3191.47 | |
| 3284.37 | |
| 4003.33 | |
| 4235.02 | L36 |
| 4893.63 | |
| 4960.9 | |
| 6062.03 | |
| 6381.52 | |
| 6496.51 | L32 |
| 6567.57 | |
| 6789.99 | |

Table S107 shows the annotated ribosomal protein mass peaks of *Staphylococcus xylosus* ATCC 29971. Specifically, ribosomal protein L36 and L32 mapped to mass peaks 4235.02 Da, and 6496.51 Da, respectively.

| **Table S108:** ***Vibrio alginolyticus*** **Peak list of ATCC 17749** | **Ribosomal protein** |
|---|---|
| 2094.68 | |
| 2331.03 | |
| 2591.26 | |
| 3151.38 | |
| 3279.83 | |
| 3597.63 | |
| 4046.66 | |
| 4179.05 | |
| 4277.79 | L36 |
| 4532.84 | |
| 4726.32 | |
| 4951.36 | |
| 5148.76 | |
| 5180.58 | L34 |
| 5524.28 | |
| 5664.42 | |
| 6167.36 | |
| 6407.06 | |
| 6467.1 | |
| 7121.49 | |
| 7193.93 | L29 |
| 8092.38 | |
| 8756.66 | |
| 9451.15 | |
| 9902.6 | |

Table S108 shows the annotated ribosomal protein mass peaks of *Vibrio alginolyticus* ATCC 17749. Specifically, ribosomal protein L36, L34 and L29 mapped to mass peaks 4277.79 Da, 5180.58 Da, and 7193.93 Da, respectively.

| **Table S109:** ***Vibrio parahaemolyticus*** **Peak list of ATCC 17802** | **Ribosomal protein** |
|---|---|
| 2140.7 | |
| 2400.81 | |
| 2591.53 | |
| 3045.76 | |
| 3084.55 | |
| 3181.37 | |
| 3204.3 | |
| 3227.23 | |
| 3599.47 | |
| 3756.22 | |
| 4178.41 | |
| 4278.14 | L36 |
| 4537.61 | |
| 4703.08 | |
| 4732.7 | |
| 5149.13 | |
| 5179.8 | L34 |
| 6166.16 | |
| 6360.07 | |
| 6405.11 | |
| 6451.26 | |
| 7195.59 | L29 |
| 7510.02 | |
| 8354.24 | |
| 9073.56 | S16 |
| 9466.19 | |
| 9934.56 | |

Table S109 shows the annotated ribosomal protein mass peaks of *Vibrio parahaemolyticus* ATCC 17802. Specifically, ribosomal protein L36, L34, L29 and S16 mapped to mass peaks 4278.14 Da, 5179.8 Da, 7195.59 Da, and 9073.56 Da, respectively.

| **Table S110:** ***Vibrio vulnificus*** **Peak list of ATCC 27562** | **Ribosomal protein** |
|---|---|
| 2140.18 | |
| 2605.15 | |
| 3064.92 | |
| 3085.05 | |
| 3170.54 | |
| 3224.19 | |
| 3322.94 | |
| 3566.34 | |
| 3596.73 | |
| 3723.41 | |
| 3994.31 | |
| 4046.77 | |
| 4178.86 | |
| 4278.5 | |
| 4322.83 | |
| 4532.21 | |
| 4754.99 | |
| 4879.14 | |
| 4958.72 | |
| 5024.65 | |
| 5156.38 | |
| 5207.57 | |
| 5404.17 | |
| 5844.2 | |
| 6167.28 | |
| 6223.53 | |
| 6446.06 | |
| 6495.37 | |
| 7191.17 | L29 |
| 8088.56 | |
| 8356.19 | |
| 9064.62 | |
| 9508.68 | S20 |
| 9909.61 | |

Table S110 shows the annotated ribosomal protein mass peaks of *Vibrio vulnificus* ATCC 27562. Specifically, ribosomal protein L29 and S20 mapped to mass peaks 7191.17 Da, and 9508.68 Da, respectively.

**Table S111: Bacterial species analyzed in this work**

| | | |
|---|---|---|
| *Acinetobacter baumanii* | *Aeromonas hydrophila* | *Bacillus amyloliquefaciens* |
| *Bacillus cereus* | *Bacillus licheniformis* | *Bacillus megaterium* |
| *Bacillus pumilus* | *Bacillus subtilis* | *Bacillus thuringiensis* |
| *Carnobacterium divergens* | *Carnobacterium gallinarum* | *Carnobacterium maltaromaticum* |
| *Citrobacter freundii* | *Clostridium botulinum* | *Clostridium perfringens* |
| *Enterobacter aerogenes* | *Enterobacter cloacae* | *Enterobacter hormaechei* |
| *Enterobacter sakazakii* | *Escherichia coli* | *Hafnia alvei* |
| *Klebsiella oxytoca* | *Klebsiella pneumoniae* | *Listeria innocua* |
| *Listeria ivanovii* | *Listeria monocytogenes* | *Listeria seeligeri* |
| *Listeria welshimeri* | *Morganella morganii* | *Pantoea agglomerans* |
| *Photobacterium damselae* | *Photobacterium phosphoreum* | *Proteus mirabilis* |
| *Proteus penneri* | *Proteus vulgaris* | *Providencia rettgeri* |
| *Providencia stuartii* | *Pseudomonas fluorescens* | *Pseudomonas fragi* |
| *Pseudomonas putida* | *Pseudomonas syringae* | *Raoultella planticola* |
| *Serratia liquefaciens* | *Serratia marcescens* | *Serratia proteamaculans* |
| *Shewanella algae* | *Shewanella baltica* | *Shewanella putrefaciens* |
| *Staphylococcus aureus* | *Staphylococcus epidermidis* | *Staphylococcus pasteuri* |
| *Staphylococcus xylosus* | *Stenotrophomonas maltophilia* | *Vibrio alginolyticus* |
| *Vibrio parahaemolyticus* | *Vibrio vulnificus* | |

**Table S112: Bacterial species selected for analyzing the phylogenetic potential of individual ribosomal protein**

| | | |
|---|---|---|
| *Aeromonas hydrophila* | *Bacillus subtilis* | *Carnobacterium maltaromaticum* |
| *Citrobacter freundii* | *Clostridium botulinum* | *Enterobacter cloacae* |
| *Escherichia coli* | *Hafnia alvei* | *Klebsiella pneumoniae* |
| *Listeria monocytogenes* | *Morganella morganii* | *Pantoea agglomerans* |
| *Photobacterium phosphoreum* | *Proteus vulgaris* | *Providencia rettgeri* |
| *Pseudomonas fluorescens* | *Raoultella planticola* | *Serratia marcescens* |
| *Shewanella putrefaciens* | *Staphylococcus aureus* | *Stenotrophomonas maltophilia* |
| *Vibrio parahaemolyticus* | | |

**Conflicts of interest**

The author declares no conflicts of interest.

**Funding**

No funding was used in this work.
